# An eQTL-based Approach Reveals Candidate Regulators of LINE-1 RNA Levels in Lymphoblastoid Cells

**DOI:** 10.1101/2023.08.15.553416

**Authors:** Juan I. Bravo, Chanelle R. Mizrahi, Seungsoo Kim, Lucia Zhang, Yousin Suh, Bérénice A. Benayoun

## Abstract

Long interspersed element 1 (L1) are a family of autonomous, actively mobile transposons that occupy ∼17% of the human genome. A number of pleiotropic effects induced by L1 (promoting genome instability, inflammation, or cellular senescence) have been observed, and L1’s contributions to aging and aging diseases is an area of active research. However, because of the cell type-specific nature of transposon control, the catalogue of L1 regulators remains incomplete. Here, we employ an eQTL approach leveraging transcriptomic and genomic data from the GEUVADIS and 1000Genomes projects to computationally identify new candidate regulators of L1 RNA levels in lymphoblastoid cell lines. To cement the role of candidate genes in L1 regulation, we experimentally modulate the levels of top candidates *in vitro*, including *IL16, STARD5, HSDB17B12,* and *RNF5*, and assess changes in TE family expression by Gene Set Enrichment Analysis (GSEA). Remarkably, we observe subtle but widespread upregulation of TE family expression following *IL16* and *STARD5* overexpression. Moreover, a short-term 24-hour exposure to recombinant human IL16 was sufficient to transiently induce subtle, but widespread, upregulation of *L1* subfamilies. Finally, we find that many L1 expression-associated genetic variants are co-associated with aging traits across genome-wide association study databases. Our results expand the catalogue of genes implicated in L1 RNA control and further suggest that L1-derived RNA contributes to aging processes. Given the ever-increasing availability of paired genomic and transcriptomic data, we anticipate this new approach to be a starting point for more comprehensive computational scans for transposon transcriptional regulators.

## Introduction

Transposable elements (TEs) constitute ∼45% of the human genome (Lander et al. 2001). Among these, the long interspersed element-1 (LINE-1 or L1) family of transposons is the most abundant, accounting for ∼16-17% (Lander et al. 2001; Venter et al. 2001), and remains autonomously mobile, with humans harboring an estimated 80-100 retrotransposition-competent L1 copies (Brouha et al. 2003). These retrotransposition-competent L1s belong to evolutionarily younger L1Hs subfamily, are ∼6 kilobases long, carry an internal promoter in their 5’-untranslated region (UTR), and encode two proteins — L1ORF1p and L1ORF2p — that are necessary for transposition (Moran et al. 1996). The remaining ∼500,000 copies are non-autonomous or immobile because of the presence of inactivating mutations or truncations (Lander et al. 2001) and include L1 subfamilies of all evolutionary ages, including the evolutionarily older L1P and L1M subfamilies. Though not all copies are transposition competent, L1s can nevertheless contribute to aspects of aging (Bravo et al. 2020; Della Valle et al. 2022) and aging-associated diseases (Liu et al. 2019; Simon et al. 2019; Zhao et al. 2021; Zhao et al. 2022).

Though mechanistic studies characterizing the role of L1 in aging and aging-conditions are limited, its effects are pleiotropic. For example, L1 can contribute to genome instability via insertional mutagenesis. Indeed, an expansion of L1 copy number with organismal aging (De Cecco et al. 2013b) and during cellular senescence (De Cecco et al. 2013a) have been documented, though many of these copies are likely cytosolic or extra-chromosomal. L1 can also play a contributing role in shaping inflammatory and cellular senescence phenotypes. The secretion of a panoply of pro-inflammatory factors is a hallmark of cell senescence, called the senescence associated secretory phenotype (SASP) (Campisi 2013). Importantly, the SASP is believed to stimulate the innate immune system and contribute to chronic, low-grade, sterile inflammation with age, a phenomenon referred to as “inflamm-aging” (Campisi 2013; Franceschi et al. 2018). During deep senescence, L1 are transcriptionally de-repressed and consequently generate cytosolic DNA that initiates an immune response consisting of the production and secretion of pro-inflammatory interferons (De Cecco et al. 2019). Finally, L1 is causally implicated in aging-associated diseases, including cancer. L1 may contribute to cancer by (i) serving as a source for chromosomal rearrangements that can lead to tumor-suppressor genes deletion (Rodriguez-Martin et al. 2020) or (ii) introducing its active promoter next to normally silenced oncogenes (Flasch et al. 2022). Thus, because of the pathological effects L1 can have on hosts, it is critical that hosts maintain precise control over L1 activity.

Eukaryotic hosts have evolved several pre- and post-transcriptional mechanisms for regulating TEs (Levin and Moran 2011; Rebollo et al. 2012). Nevertheless, our knowledge of regulatory genes remains incomplete because of cell type-specific regulation and the complexity of methods required to identify regulators. Indeed, one clustered regularly interspaced short palindromic repeats (CRISPR) screen in two cancer cell lines for regulators of L1 transposition identified >150 genes involved in diverse biological functions (Liu et al. 2018) (*e.g*. chromatin regulation, DNA replication, and DNA repair). However, only about ∼36% of the genes identified in the primary screen exerted the same effects in both cell lines (Liu et al. 2018), highlighting the potentially cell type-specific nature of L1 control. Moreover, given the complexities of *in vitro* screens, especially in non-standard cell lines or primary cells, *in silico* screens for L1 regulators may facilitate the task of identifying and cataloguing candidate regulators across cell and tissue types. One such attempt was made by generating gene-TE co-expression networks from RNA sequencing (RNA-seq) data generated from multiple cancer-adjacent tissue types (Chung et al. 2019). Although co-expression modules with known TE regulatory functions, such as interferon signaling, were correlated with TE modules, it is unclear whether other modules may harbor as of now uncharacterized TE-regulating properties, since no validation experiments were carried out. Additionally, this co-expression approach is limited, as no mechanistic directionality can be assigned between associated gene and TE clusters, complicating the prioritization of candidate regulatory genes for validation. Thus, there is a need for the incorporation of novel “omic” approaches to tackle this problem. Deciphering the machinery that controls TE activity in healthy somatic cells will be crucial, in order to identify checkpoints lost in diseased cells.

The 1000Genomes Project and GEUVADIS Consortium provide a rich set of genomic resources to explore the mechanisms of human TE regulation *in silico*. The 1000Genomes project generated a huge collection of genomic data from thousands of human subjects across the world, including single nucleotide variant (SNV) and structural variant (SV) data (Auton et al. 2015; Sudmant et al. 2015). To accomplish this, the project relied on lymphoblastoid cell lines (LCLs), which are generated by infecting resting B-cells in peripheral blood with Epstein-Barr virus (EBV). Several properties make them advantageous for use in large-scale projects (e.g. they can be generated relatively noninvasively, provide a means of obtaining an unlimited amount of a subject’s DNA and other biomolecules, and can serve as an *in vitro* model for studying the effects of genetic variation with phenotypes of interest) (Sie et al. 2009; Hussain and Mulherkar 2012). Naturally, the GEUVADIS Consortium generated transcriptomic data for a subset of subjects sampled by the 1000Genomes Project and carried out an expression quantitative trait locus (eQTL) analysis to define the effects of genetic variation on gene expression (Lappalainen et al. 2013). Later, in a series of landmark studies on TE biology, this collection of data was reanalyzed (i) to characterize the effects of polymorphic TE structural variation on gene expression (TE-eQTLs) (Wang et al. 2016; Spirito et al. 2019; Goubert et al. 2020) and (ii) to explore the potential impact of TE polymorphisms on human health and disease through genome-wide association study (GWAS) analysis (Wang et al. 2017). These results highlight the value of this data and the power of eQTL analysis in identifying genetic factors implicated in gene expression control and, potentially, disease susceptibility. Together, these resources provide a useful toolkit for investigating the genetic regulation of TEs, generally, and L1, specifically.

Much work on the mechanisms of L1 regulation has been carried out by looking exclusively at full-length, transposition-competent L1 elements, as this has allowed for the study of the whole L1 replication lifecycle, starting from transcription at the internal promoter and ending with transposition into a new genomic site (Liu et al. 2018; Mita et al. 2020). However, the total L1 RNA pools can be influenced by a number of other sources, including L1 copies residing in introns, L1 copies that are exonized, L1 copies that are co-transcribed because of nearby genes, and L1 copies that are independently transcribed from their own promoter regardless of their ability to mobilize. Thus, cellular L1 RNA levels are likely to be modulated by a number of transcriptional and post-transcriptional processes, including promoter-dependent transcription, RNA turnover, exonization, and/or intron retention (among others). However, the control mechanisms for non-full length transposition-competent L1 elements remain incompletely characterized, even though there is increasing evidence that these can have important regulatory and functional potential. For example, one study suggested that intronic L1s are part of an important regulatory network maintaining T-cell quiescence (Marasca et al. 2022). This is consistent with the increasing appreciation for the importance of alternative splicing in immune regulation and cell death pathways (Liao and Garcia-Blanco 2021) and with another study highlighting the importance of TE exonization in interferon signaling (Pasquesi et al. 2023). Additionally, L1 RNA may be sufficient to induce an interferon response and alter cellular viability, even in the absence of transposition (Ardeljan et al. 2020; Luqman-Fatah et al. 2023). Given these observations, there is a need to characterize the control mechanisms and functional effects of all L1 RNA sources, including truncated, non-autonomous or transposition-incompetent copies.

In this study, we (i) develop a new pipeline to identify novel candidate regulators of L1 RNA levels in lymphoblastoid cell lines, including intronic, intergenic, and exon-overlapping RNA levels, (ii) provide experimental evidence for the involvement of top candidates in L1 RNA level control, and (iii) expand and reinforce the catalog of diseases linked to differential L1 levels.

## Results

### In silico scanning for L1 subfamily candidate regulators by eQTL analysis

To identify new candidate regulators of L1 RNA levels, we decided to leverage publicly available human “omic” datasets with both genetic and transcriptomic information. For this analysis, we focused on samples for which the following data was available: (i) mRNA-seq data from the GEUVADIS project, (ii) SNVs called from whole-genome sequencing data overlayed on the hg38 human reference genome made available by the 1000Genomes project, and (iii) repeat structural variation data made available by the 1000Genomes project. This yielded samples from 358 European and 86 Yoruban individuals, all of whom declared themselves to be healthy at the time of sample collection (**Figure 1A**). Using the GEUVADIS data, we obtained gene and TE subfamily expression counts using TEtranscripts (Jin et al. 2015). As a quality control step, we checked whether mapping rates segregated with ancestry groups, which may bias results. However, the samples appeared to cluster by laboratory rather than by ancestry (**Figure S1A**). As additional quality control metrics, we also checked whether the SNV and SV data segregated by ancestry following principal component analysis (PCA). These analyses demonstrated that the top two and the top three principal components from the SNV and SV data, respectively, segregated ancestry groups (**Figure S1B, Figure S1C**).

**Figure 1.**
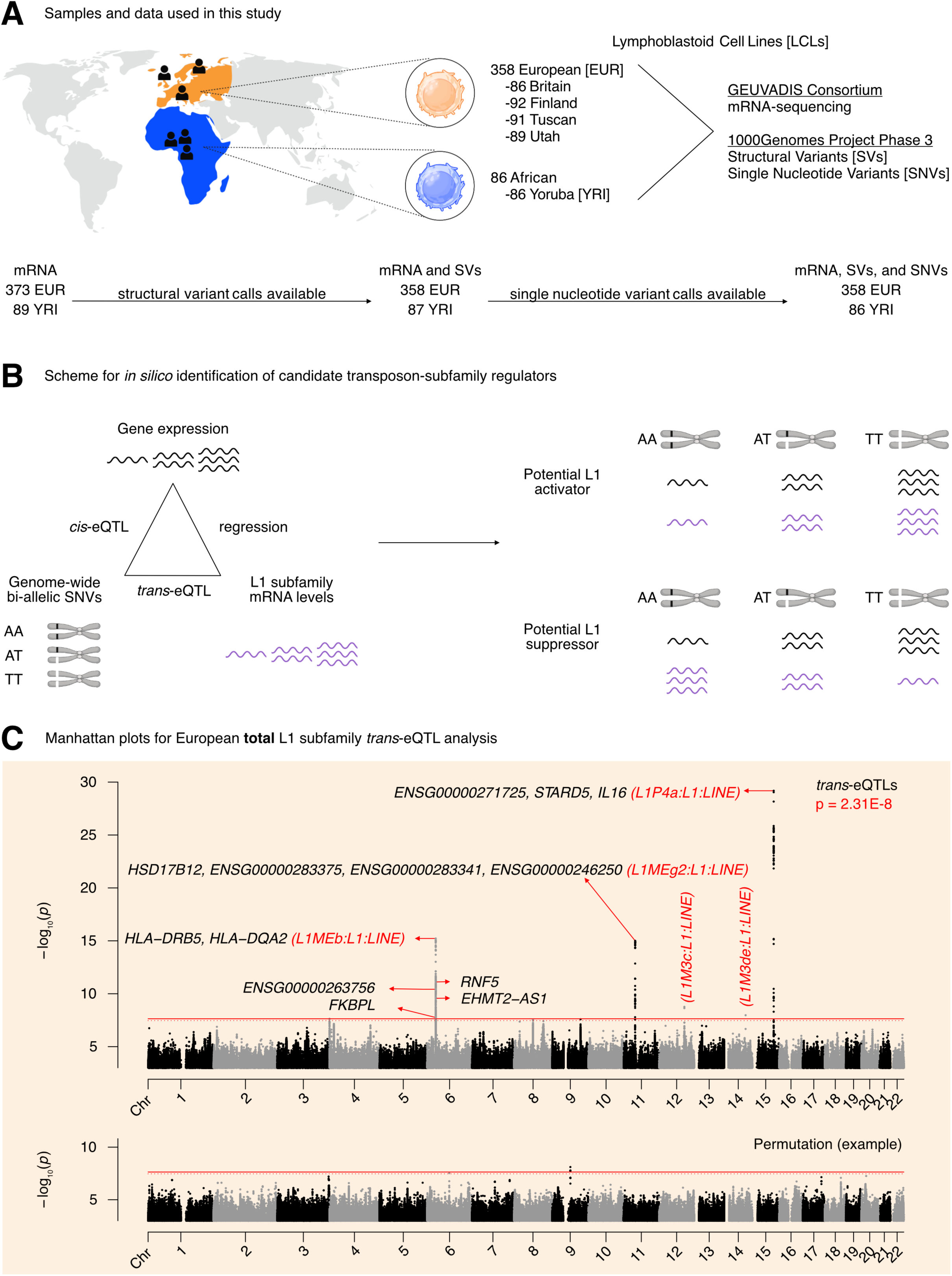
Overview of the pipeline developed to scan for L1 transcriptional regulators *in silico*. **(A)** An illustration of the samples and “omic” data used in this study. Of the 358 European individuals, 187 were female and 171 were male. Of the 86 African individuals, 49 were female and 37 were male. (Note that Utah subjects are of Northern European ancestry, and thus part of the European cohort for analytical purposes). **(B)** A schematic illustrating how genetic variants, gene expression, and TE expression can be integrated to identify highly correlated SNV-Gene-TE trios. **(C)** A Manhattan plot for the L1 subfamily *trans*-eQTL analysis in the European cohort. The genes that passed our three-part integration approach are listed next to the most significant *trans*-eQTL SNV they were associated with in *cis*. The dashed line at p = 3.44E-8 corresponds to an average empirical FDR < 0.05, based on 20 random permutations. One such permutation is illustrated in the bottom panel. The solid line at p = 2.31E-8 corresponds to a Benjamini-Hochberg FDR < 0.05. The stricter of the two thresholds, p = 2.31E-8, was used to define significant *trans*-eQTLs. FDR: False Discovery Rate. Some panels were created with BioRender.com.

We then chose to do a three-part integration of the available “omic” data (**Figure 1B**). Since TEtranscripts quantifies total TE RNA levels aggregated at TE subfamily resolution and discards TE position information, we chose to carry out a *trans*-eQTL analysis against global RNA levels of each L1 subfamily. We reasoned that there would have to be factors (i.e., miRNAs, proteins, non-coding RNAs) mediating at least a subset of the effects of SNVs on L1 subfamily RNA levels. Thus, to identify candidate genic mediators, we searched for genes with *cis*-eQTLs that overlapped with L1 *trans*-eQTLs. As a final filter, we reasoned that for a subset of regulators, L1 subfamily RNA levels would respond to changes in the expression of those regulators. Consequently, we chose to quantify the association between L1 subfamily RNA levels and candidate gene expression by linear regression. Importantly, to avoid confounding eQTL associations with extraneous technical and biological factors, the expression data was corrected for the following: laboratory, population category, genetic population structure, biological sex, net L1 and Alu copy number called from the SV data, and EBV expression levels. We hypothesized that this three-part integration would result in combinations of significantly correlated SNVs, genes, and L1 subfamilies (**Figure 1B**).

The *trans*-eQTL analysis for RNA levels against every detected L1 subfamily led to the identification of 499 *trans*-eQTLs distributed across chromosomes 6, 11, 12, 14, and 15 that passed genome-wide significance (**Figure 1C, Supplementary Table S1A**). The *cis*-eQTL analysis led to the identification of 845,260 *cis*-eQTLs that passed genome-wide significance (**Supplementary Figure S2A, Supplementary Table S1B**). After integrating the identified *cis*- and *trans*-eQTLs and running linear regression, we identified 1,272 SNV-Gene-L1 trios that fulfilled our three-part integration approach (**Supplementary Table S1C**). Among this pool of trios, we identified 7 unique protein-coding genes including (i) *IL16* and *STARD5* which were correlated with *L1P4a* levels, (ii) *HLA-DRB5, HLA-DQA2, RNF5,* and *FKBPL* which were correlated with *L1MEb* levels, and (iii) *HSD17B12* which was correlated with *L1MEg2* levels (**Figure 1C**). Although *EHMT2* did not pass our screening approach, it does overlap *EHMT2-AS1*, which did pass our screening thresholds. In contrast, we also identified “orphan” SNVs on chromosomes 12 and 14 which were associated with *L1M3c* and *L1M3de* levels in *trans* but to which were unable to attribute a candidate gene. These SNVs resided in intronic regions within *NTN4* and *STON2*, respectively. We note that detection of these gene and TE associations is unlikely to be mechanistically related to variations in EBV expression, as expression profiles were corrected for such differences before downstream analyses (**Supplementary Figure S2B**). We also note that several other unique non-coding genes, often overlapping the protein-coding genes listed, were also identified (**Figure 1C**). For simplicity of interpretation, we focused on protein-coding genes during downstream experimental validation.

Next, to define first and second tier candidate regulators, we clumped SNVs in linkage disequilibrium (LD) by L1 *trans*-eQTL p-value to identify the most strongly associated genetic variant in each genomic region (**Figure 2A, Supplementary Figure S3A**). LD-clumping identified the following index SNVs (*i.e*. the most strongly associated SNVs in a given region): rs11635336 on chromosome 15, rs9271894 on chromosome 6, rs1061810 on chromosome 11, rs112581165 on chromosome 12, and rs72691418 on chromosome 14 (**Supplementary Table S1D**). Genes linked to these SNVs were considered first tier candidate regulators and included *IL16*, *STARD5*, *HLA-DRB5*, *HLA-DQA2*, and *HSD17B12* (**Figure 2B, Supplementary Table S1E**). The remaining genes were linked to clumped, non-index SNVs and were consequently considered second tier candidates and included *RNF5*, *EHMT2-AS1*, and *FKBPL* (**Supplementary Figure S3B**). Additionally, for simplicity of interpretation, we considered only non-*HLA* genes during downstream experimental validation, since validation could be complicated by the highly polymorphic nature of *HLA* loci (Williams 2001) and their involvement in multi-protein complexes.

**Figure 2.**
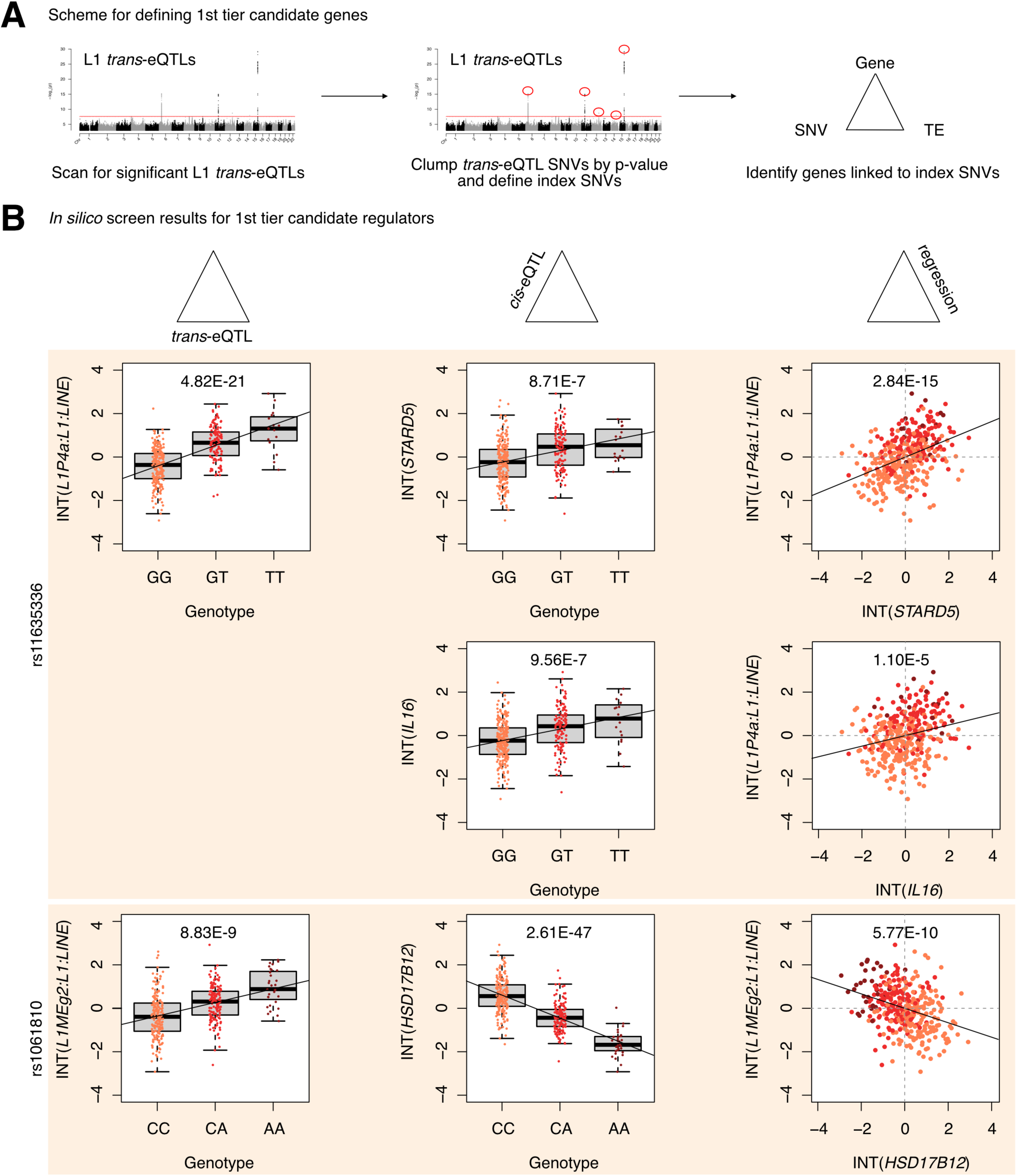
Identification of 1^st^ tier candidate L1 expression regulators in the European cohort. **(A)** A schematic for how 1^st^ tier candidate genes were defined. In short, these were genes in trios with index SNVs that were at the top of their respective peak. **(B)** The three-part integration results for three protein-coding genes—*STARD5*, *IL16*, *HSD17B12*—that we considered first tier candidates for functional, *in vitro* testing. In the left column are the *trans*-eQTLs, in the middle column are the *cis*-eQTLs, and in the right column are the linear regressions for gene expression against L1 subfamily expression. Expression values following an inverse normal transform (INT) are shown. The FDR for each analysis is listed at the top of each plot. FDR: False Discovery Rate.

Finally, to computationally determine whether candidate genes may causally influence L1 subfamily RNA levels, we carried out mediation analysis on all SNV-gene-L1 trios (**Supplementary Figure S4A**). Interestingly, 868 out of the 1,272 (68.2%) trios exhibited significant (FDR < 0.05) mediation effects (**Supplementary Table S1F**). Among the 1st tier candidate regulators, significant, partial, and consistent mediation effects could be attributed to *STARD5*, *IL16*, *HSD17B12*, and *HLA-DRB5* (**Supplementary Figure S4B, Supplementary Table S1F**). To note, while significant mediation could be attributed to the index SNV for *STARD5*, significant mediation could only be attributed to clumped SNVs for *IL16* and *HSD17B12*. Given that *STARD5* and *IL16* share *cis*-eQTL SNVs, this suggests that *STARD5* may be the more potent mediator. Among the 2nd tier candidate regulators, significant, partial, and consistent mediation effects could be attributed to *RNF5*, *EHMT2-AS1*, and *FKBPL* (**Supplementary Figure S4C, Supplementary Table S1F**). These results suggest that candidate genes may mediate the effects between linked SNVs and L1 subfamilies.

### In silico scanning for L1 subfamily candidate regulators in an African population

We sought to assess the cross-ancestry regulatory properties of candidate genes by repeating our scan using the Yoruban samples as a smaller but independent replication cohort. Here, rather than conduct a genome-wide scan for *cis*- and *trans*-associated factors, we opted for a targeted approach focusing only on gene *cis*-eQTLs and L1 subfamily *trans*-eQTLs that were significant in the analysis with European samples (**Supplementary Figure S5A**). The targeted *trans*-eQTL analysis led to the identification of 227 significant (FDR < 0.05) *trans*-eQTLs distributed across chromosomes 6 and 11 (**Supplementary Table S2A**). The targeted *cis*-eQTL analysis led to the identification of 1,248 significant (FDR < 0.05) *cis*-eQTLs (**Supplementary Table S2B**). After integrating the identified *cis*- and *trans*-eQTLs and running linear regression, we identified 393 SNV-Gene-L1 trios that fulfilled our three-part integration approach (**Supplementary Table S2C**). Among this pool of trios, we identified 2 unique protein-coding genes—*HSD17B12 and HLA-DRB6—*as well as several unique non-coding genes (**Supplementary Table S2C**). Again, we clumped SNVs in linkage disequilibrium by L1 *trans*-eQTL p-value. LD-clumping identified the following index SNVs: rs2176598 on chromosome 11 and rs9271379 on chromosome 6 (**Supplementary Table S2D**). Genes linked to these SNVs were considered first tier candidate regulators and included both *HSD17B12* and *HLA-DRB6* (**Supplementary Figure S5B, Supplementary Table S2E**). Finally, we carried out mediation analysis on all SNV-gene-L1 trios; however, no significant (FDR < 0.05) mediation was observed (**Supplementary Table S2F**). These results implicate *HSD17B12* and the *HLA* loci as candidate, cross-ancestry regulators of L1 RNA levels.

To assess why some candidate genes did not replicate in the Yoruba cohort, we manually inspected *cis*- and *trans*-eQTL results for trios with those genes (**Supplementary Figure S6A**). Interestingly, we identified rs9270493 and rs9272222 as significant (FDR < 0.05) *trans*-eQTLs for *L1MEb* RNA levels. However, those SNVs were not significant *cis*-eQTLs for *RNF5* and *FKBPL* expression, respectively. For trios involving *STARD5*, *IL16*, and *EHMT2-AS1*, neither the *cis-*eQTL nor the *trans*-eQTL were significant. We note that for most of these comparisons, although the two genotypes with the largest sample sizes were sufficient to establish a trending change in *cis* or *trans* RNA levels, this trend was often broken by the third genotype with spurious sample sizes. This suggests that replication in the Yoruba cohort may be limited by the small cohort sample size in the GEUVADIS project.

### Stratified in silico scanning for candidate regulators of intronic, intergenic, or exon-overlapping L1 subfamily RNA levels

One potential limitation with the approach undertaken thus far is the inability to distinguish different transposon RNA sources. For example, intergenic TEs may be transcribed using their own promoter or using a nearby gene’s promoter. In contrast, while intronic or exon-overlapping TEs may be independently transcribed if they have retained their promoter, they may also appear expressed due to intron retention or due to exonization events. To have further granularity in deciphering L1 RNA level regulators with respect to the genomic provenance of their RNA sources, we next (i) carried out locus-specific quantification for each TE locus, (ii) stratified loci by whether they were intronic, nearby intergenic (within 5 kb of a gene), distal intergenic (>5 kb from a gene) or exon-overlapping, (iii) aggregated counts within each category at the subfamily level to compare with our unstratified TEtranscripts results, and (iv) repeated our eQTL scan using the four stratified TE RNA profiles (**Supplementary Table S3A-S3D**; **Supplementary Figure S7A-D**).

First, the eQTL scan using the intronic L1 profiles recapitulated the results of our initial scan for the *IL16/STARD5* and *HSD17B12* loci (**Supplementary Figure S7A, Supplementary Table S3E)**. Interestingly, we recovered a dominant, new peak on chromosome 4 that was associated with *L1M3a* RNA levels but to which we could not attribute a protein-coding mediator. The index SNV for this locus, rs6819237, resides within an intron of *ZNF141*, which has been shown to bind L1PA elements (de Tribolet-Hardy et al. 2023). Second, the eQTL scan using the nearby intergenic L1 profiles recapitulated the results of our initial scan for the *HLA* loci (**Supplementary Figure S7B, Supplementary Table S3F)**. Third, for distal intergenic L1 expression, we identified a cluster of SNVs on chromosome 6 that were co-associated with *L1MC5* RNA levels and *ZSCAN26* expression (**Supplementary Figure S7C, Supplementary Table S3G)**. Interestingly, these SNVs reside in a genomic region with many other *ZSCAN* genes, including *ZKSCAN4* which is hypothesized to regulate L1PA5/PA6 transcripts (Helleboid et al. 2019) and *ZSCAN9* which was shown to bind L1 by MapRRCon (Sun et al. 2018) analysis. Finally, we also identified several loci associated with exon-overlapping L1 RNA levels (**Supplementary Figure S7D, Supplementary Table S3H)**.

Since intergenic L1s, as a potential source of independently transcribed L1 RNA, are of special interest, we repeated the mediation analysis for the *ZSCAN26*-associated SNV rs1361387 (**Supplementary Figure S8A**). Alternating the genotype of rs1361387 was associated with an increase in *L1MC5* RNA levels and a decrease in *ZSCAN26* expression (**Supplementary Figure S8B**). Mediation analysis revealed significant, but inconsistent mediation of *L1MC5* through *ZSCAN26* (**Supplementary Figure S8C, Supplementary Table S3I**). This may suggest that rs1361387 may exert both positive and negative control of *L1MC5* through uncharacterized mechanisms. Taken together, these results suggest that our approach can detect known L1 RNA regulators.

### TE families and known TE-associated pathways are differentially regulated across L1 trans-eQTL variants

Though our eQTL analysis identified genetic variants associated with the levels of specific, evolutionarily older L1 subfamilies, we reasoned that there may be more global but subtle differences in TE expression profiles among genotype groups, given that TE levels across subfamilies is highly correlated (Chung et al. 2019). Thus, for each gene-associated index SNV identified in the European eQTL analysis, we carried out differential expression analysis for all expressed genes and TEs (**Figure 3A**). At the individual gene level, we detected few significant (FDR < 0.05) changes: 4 genes/TEs varied with rs11635336 genotype (*IL16*/*STARD5*), 4 genes/TEs varied with rs9271894 genotype (*HLA*), and 5 genes/TEs varied with rs1061810 genotype (*HSD17B12*) (**Supplementary Table S4A-S4C**). Importantly, however, these genes/TEs overlapped the genes/TEs identified in the *cis-* and *trans*-eQTL analyses, providing an algorithm-independent link among candidate SNV-gene-TE trios.

**Figure 3.**
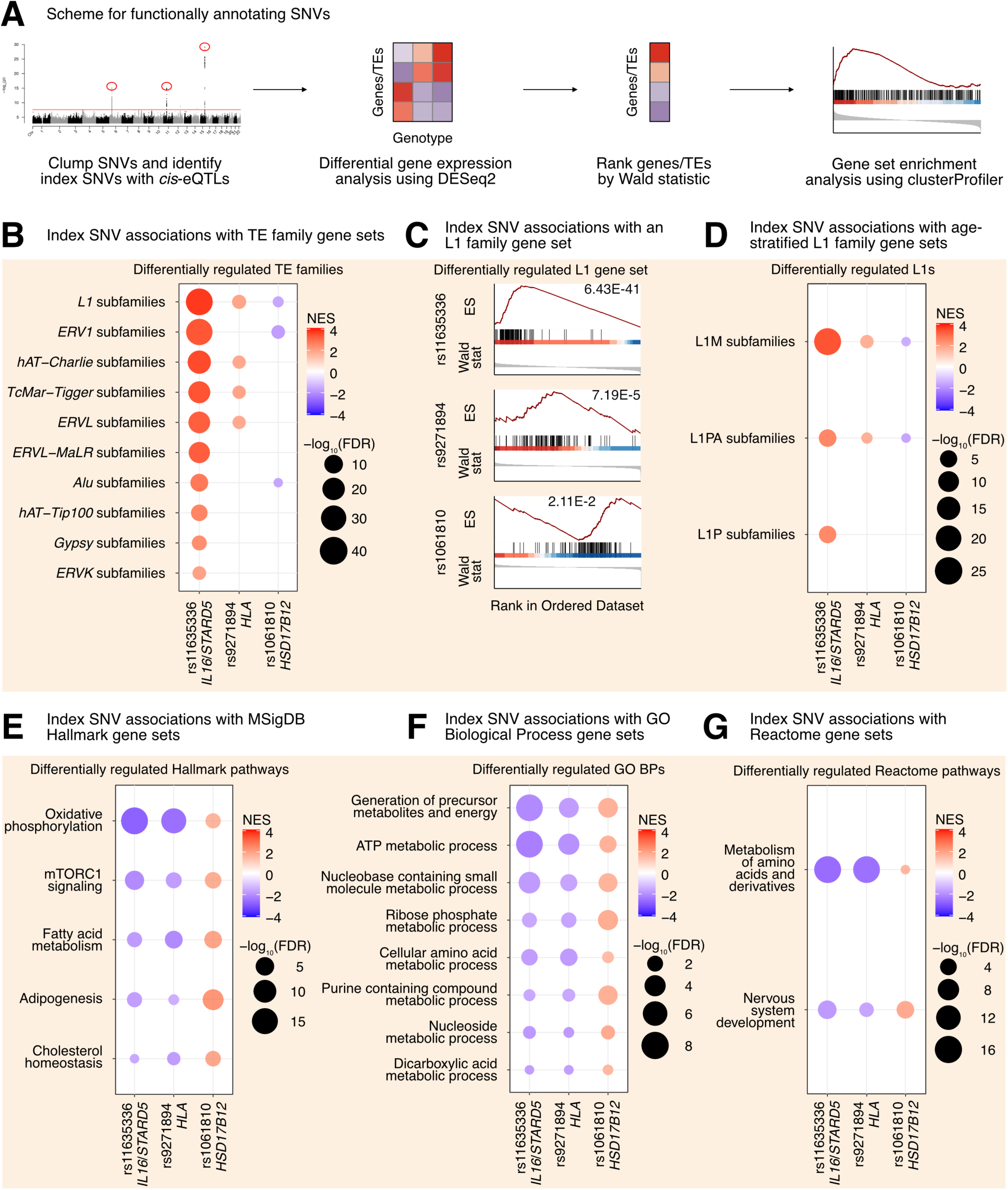
L1 *trans*-eQTLs are associated with subtle, widespread differences in TE families and known TE-associated pathways. **(A)** Scheme for functionally annotating gene-linked index SNVs by GSEA. **(B)** GSEA analysis for shared, significantly regulated TE family gene sets across genotypes for rs11635336 (*IL16/STARD5*), rs9271894 (*HLA*), and rs1061810 (*HSD17B12*). **(C)** GSEA plots for the L1 family gene set results summarized in **(B)**. For these plots, the FDR value is listed. **(D)** GSEA analysis for shared, significantly regulated, evolutionary-age-stratified L1 gene sets across genotypes for rs11635336 (*IL16/STARD5*), rs9271894 (*HLA*), and rs1061810 (*HSD17B12*). L1M subfamilies are the oldest, L1P subfamilies are intermediate, and L1PA subfamilies are the youngest. GSEA analysis for top, shared, concomitantly regulated **(E)** MSigDB Hallmark pathway, **(F)** GO Biological Process, and **(G)** Reactome pathway gene sets across genotypes for rs11635336 (*IL16/STARD5*), rs9271894 (*HLA*), and rs1061810 (*HSD17B12*). Shared gene sets were ranked by combining p-values from each individual SNV analysis using Fisher’s method. In each bubble plot, the size of the dot represents the -log_10_(FDR) and the color reflects the normalized enrichment score. FDR: False Discovery Rate.

In contrast to gene-level analyses, Gene Set Enrichment Analysis (GSEA) provides increased sensitivity to subtle, but consistent and widespread, transcriptomic changes at the level of gene sets (*e.g*. TE families, biological pathways, etc.) (Subramanian et al. 2005). Specifically, GSEA was developed in response to (i) the lack of reproducibility of individual, significant gene changes across studies and (ii) the need to summarize biological changes in a functionally meaningful way through the use of gene sets containing biologically-related genes. For the TEs, we opted to aggregate TE subfamilies into gene sets corresponding to TE families on the basis that (i) broad changes in individual TE subfamilies may be hard to detect but changes across many subfamilies would be easier to detect, (ii) the expression of many different TE subfamilies was previously found to be highly correlated in an analysis of tumor-adjacent tissue (Chung et al. 2019), (iii) we were searching for factors influencing global TE RNA levels and not just specific TE loci, and (iv) GSEA has previously been applied to summarize TE changes (Gu et al. 2021; Wong et al. 2021; Zhang et al. 2023). We leveraged our differential expression analysis in combination with GSEA to identify repeat family and biological pathway gene sets impacted by SNV genotype in the GEUVADIS dataset (**Supplementary Table S4D-S4O**; **Figure 3A**). Interestingly, changes in the genotype of rs11635336 (*IL16*/*STARD5*), rs9271894 (*HLA*), and rs1061810 (*HSD17B12*) were associated with an upregulation, upregulation, and downregulation, respectively, of multiple TE family gene sets (**Figure 3B, Supplementary Table S4P**). Differentially regulated TE family gene sets included DNA transposons, such as the hAT-Charlie family, and long terminal repeat (LTR) transposons, such as the endogenous retrovirus-1 (ERV1) family (**Figure 3B, Supplementary Table S4P**). Noteworthy, the L1 family gene set was the only TE gene set whose expression level was significantly altered across all three SNV analyses (**Figure 3B, Supplementary Table S4P**). Consistent with their relative significance in the L1 *trans*-eQTL analysis, the L1 family gene set was most strongly upregulated by alternating the *IL16*/*STARD5* SNV (NES = 3.74, FDR = 6.43E-41), intermediately upregulated by alternating the *HLA* SNV (NES = 1.90, FDR = 7.19E-5), and least strongly changed by alternating the *HSD17B12* SNV (NES = -1.57, FDR = 2.11E-2) (**Figure 3C**). Among these changes, both older (L1M) and younger (L1PA) elements were differentially regulated across all three SNVs (**Figure 3D, Supplementary Table S4Q**). Overall, we observed similar effects on TE family RNA levels when, as an alternative and orthogonal approach, we applied a one-sample Wilcoxon test to determine whether TE family changes were significantly different than 0 across the three SNV DESeq2 analyses (**Supplementary Figure S9A-C**). We briefly note here that rs9270493, a clumped SNV linked to *RNF5*, was also linked to upregulation of the L1 family gene set (**Supplementary Table S4R-S4S**). These results suggest that TE subfamily *trans*-eQTLs are associated with subtle but global differences in TE RNA levels beyond a lone TE subfamily.

To determine the origin of these global TE RNA level differences, we repeated our differential expression analysis and GSEA using the genomic-region-stratified TE RNA profiles. Similar to the unstratified analysis, changes in the genotype of rs11635336 (*IL16*/*STARD5*), rs9271894 (*HLA*), and rs1061810 (*HSD17B12*) were associated with an upregulation, upregulation, and downregulation, respectively, of multiple TE family gene sets of varied genomic (**Supplementary Table S4T-S4V**; **Supplementary Figure S9D-F**). Across genotype for rs11635336 (*IL16*/*STARD5*), the most upregulated TEs were of intronic origin (**Supplementary Figure S9D**). However, distal intergenic TE RNA levels, including L1 RNA levels, were also upregulated, suggesting that TE RNA level differences are not solely due to TE exonization or co-expression with genes. In contrast, most of the TE upregulation across genotypes for rs9271894 (*HLA*) was due to intronic TE RNA, though intergenic TEs near genes were also upregulated (**Supplementary Figure S9E**). Finally, L1 RNA levels, and TE RNA levels more generally, were downregulated in the distal intergenic, nearby intergenic, and exonic categories (**Supplementary Figure S9F**). These results suggest that TE subfamily *trans*-eQTLs are associated with subtle but global differences in TE expression of varying genomic origin (intronic, exonic, or intergenic).

Next, we asked if other biological pathways were regulated concomitantly with TE gene sets in response to gene-linked index SNVs, reasoning that such pathways would act either upstream (as regulatory pathways) or downstream (as response pathways) of TE alterations. GSEA with the MSigDB Hallmark pathway gene sets (Subramanian et al. 2005; Liberzon et al. 2015) identified 5 gene sets fitting this criterion, including “oxidative phosphorylation”, “mTORC1 signaling”, “fatty acid metabolism”, “adipogenesis”, and “cholesterol homeostasis” (**Figure 3E, Supplementary Table S4W**). Interestingly, several of these pathways or genes in these pathways have been implicated in TE regulation before. Rapamycin, which acts through mTORC1, has been shown to alter the expression of L1 and other repeats (Wahl et al. 2020; Marasca et al. 2022). Estrogens, which are involved in cholesterol and lipid metabolism, have been found to drive changes in repeat expression, and the receptors for both estrogens and androgens are believed to bind repeat DNA (Sampathkumar et al. 2020; Wahl et al. 2020). Pharmacological inhibition of the mitochondrial respiratory chain and pharmacological reduction of endogenous cholesterol synthesis have also been shown to induce changes in L1 protein levels or repeat expression more broadly (Baeken et al. 2020; Valdebenito-Maturana et al. 2023). GSEA with the GO Biological Process gene sets (**Figure 3F, Supplementary Table S4X**) and the Reactome gene sets (**Figure 3F, Supplementary Table S3Y**) also identified several metabolism-related pathways including “ATP metabolic process”, “Generation of precursor metabolites and energy”, and “metabolism of amino acids and derivatives”. These results add to the catalogue of pathways associated with differences in L1 expression.

In our eQTL analysis, we also identified two orphan index SNVs, rs112581165 and rs72691418, to which we could not attribute a protein-coding gene mediator. To determine whether these SNVs also regulate any transposon families or biological pathways, we repeated the differential expression analysis (with all expressed genes and TEs) (**Supplementary Table S5A-S5B**) and the GSEA (**Supplementary Table S5C-S5J**) with these SNVs (**Supplementary Figure S10A**). At the individual gene level, we detected 3193 genes/TEs that varied significantly (FDR < 0.05) with rs112581165 genotype and 1229 genes/TEs that varied significantly with rs72691418 genotype (**Supplementary Table S5A-S5B**). Similar to above, we next carried out GSEA to identify changes in functionally relevant gene sets. Like the gene-linked index SNVs, changes in the genotype of rs112581165 and rs72691418 were both associated with a downregulation and upregulation, respectively, of 10 TE families (**Supplementary Figure S10B, Supplementary Table S5K**). Noteworthy, the L1 family gene set was among the most strongly dysregulated TE family gene sets for both rs112581165 (NES = -4.32, FDR = 5.18E-89) and rs72691418 (NES = 4.01, FDR = 5.38E-79) (**Supplementary Figure S10C**). Among L1 changes, older (L1M), intermediate (L1P), and younger (L1PA) elements were differentially regulated across both SNVs (**Supplementary Figure S10D, Supplementary Table S5L**).

Overall, we observed similar changes in TE RNA levels when we applied the alternative one-sample Wilcoxon test approach to determine whether TE family changes were significantly different than 0 across both SNV DESeq2 analyses (**Supplementary Figure S11A-B**). After stratifying TE RNA levels by genomic origin, we observed that intronic TEs were strongly differentially regulated with genotype for both SNVs. However, differential regulation of intergenic TEs (both near and far from genes) and exonic TEs were also observed (**Supplementary Table S5M-S5N**; **Supplementary Figure S11C-D**). These results suggest that TE subfamily *trans*-eQTLs are associated with subtle differences in TE expression beyond the lone TE subfamily, even in the absence of a protein-coding gene *cis*-eQTL. Additionally, the data also suggests that TE RNA changes are not solely due to exonization or intron retention events.

Like before, we asked if other biological pathways were regulated concomitantly with TE gene sets in response to orphan index SNVs. The top 10 Hallmark pathway gene sets identified by GSEA included gene sets that were previously identified (“oxidative phosphorylation”, “fatty acid metabolism”, and “mTORC1 signaling”), as well as several new pathways (**Supplementary Figure S10E, Supplementary Table S5O**). Among the new pathways, “DNA repair” (Liu et al. 2018) and the “P53 pathway” (Ardeljan et al. 2020; Tiwari et al. 2020) have also been linked to L1 control, and proteins in the “Myc targets v1” gene set interact with L1 ORF1p (Luqman-Fatah et al. 2023). GSEA with the GO Biological Process gene sets (**Supplementary Figure S10F, Supplementary Table S5P**) and the Reactome gene sets (**Supplementary Figure S10G, Supplementary Table S5Q**) identified several metabolism-related pathways and several translation-related pathways, such as “cytoplasmic translation”, “eukaryotic translation initiation”, and “eukaryotic translation elongation”. Importantly, proteins involved in various aspects of proteostasis have been shown to be enriched among L1 ORF1p-interacting proteins (Luqman-Fatah et al. 2023). Again, these results add to the catalogue of pathways associated with differences in TE expression, even in the absence of a candidate *cis* mediator.

Finally, we carried out our DESeq2 and GSEA analysis against the lone index SNV, rs1361387, associated with distal intergenic L1 RNA levels (**Supplementary Figure S12A, Supplementary Table S5R-S5U**). For this SNV, we did not detect significant changes in any TE family gene set. However, GSEA will all three pathway gene sets revealed a strong suppression of immune related processes, including “interferon gamma response”, “interferon alpha response”, “response to virus”, and “interferon alpha/beta signaling” (**Supplementary Figure S12B-S12D**). These observations are consistent with the role of L1 as a stimulator and target of the interferon pathway (De Cecco et al. 2019; Luqman-Fatah et al. 2023), as well as the notion that transposons can mimic viruses and stimulate immune responses from their hosts (Lindholm et al. 2023).

### Modulation of top candidate gene activity in a lymphoblastoid cell line induces small but widespread TE RNA level changes

We decided to validate the L1 regulatory properties of top candidate genes associated with L1 *trans*-eQTLs. For experimental purposes, we selected the GM12878 lymphoblastoid cell line, because (i) it is of the same cell type as the transcriptomic data used here for our eQTL analysis, and (ii) its epigenomic landscape and culture conditions have been well well-characterized as part of the ENCODE project (The 2011; The 2012). For validation purposes, we selected *IL16*, *STARD5*, *HSD17B12*, and *RNF5* out of the 7 protein-coding gene candidates. We chose these genes for validation because the first 3 are associated with top *trans*-eQTL SNVs and the fourth one had very strong predicted mediation effects. To note, although GM12878 was part of the 1000Genomes Project, it was not included in the GEUVADIS dataset. However, based on its genotype, we can predict the relative expression of candidate regulators (**Supplementary Figure S13A**), which suggest that GM12878 may be most sensitive to modulations in *IL16* and *STARD5* expression, given their relatively low endogenous expression. Interestingly, examination of the ENCODE epigenomic data in GM12878 cells (The 2011) demonstrated that the region near the *IL16*/*STARD5*-linked index SNV (rs11635336) was marked with H3K4Me1 and H3K27Ac, regulatory signatures of enhancers (**Supplementary Figure S13C**). Similarly, the region near the *HLA*-linked index SNV (rs9271894) was marked with H3K4Me1, marked with H3K27Ac, and accessible by DNase, suggesting regulatory properties of the region as an active enhancer (**Supplementary Figure S13C**). These results further highlight the regulatory potential of the *IL16-*, *STARD5-*, and *HLA*-linked SNVs.

First, we tested the transcriptomic impact of overexpressing our top candidates in GM12878 LCLs. Cells were electroporated with overexpression plasmids (or corresponding empty vector), and RNA was isolated after 48h (**Figure 4A, Supplementary Figure S14A**). Differential expression analysis comparing control and overexpression samples confirmed the overexpression of candidate genes (**Supplementary Figure S14B, Supplementary Table S6A-S6D**). We note that no significant differences in EBV expression were identified in any of the four conditions (all FDR > 0.05; **Supplementary Figure S14C**). Intriguingly, we observed that *IL16* was significantly upregulated following *STARD5* overexpression (**Supplementary Figure S14D, Supplementary Table S6B**), although the inverse was not observed (**Supplementary Table S6A**), suggesting that *IL16* may act downstream of *STARD5*. We note here that, consistent with the use of a high expression vector, the *IL16* upregulation elicited by *STARD5* overexpression (log_2_ fold change = 0.45) was weaker than the upregulation from the *IL16* overexpression (log_2_ fold change = 1.89) (**Supplementary Table S6A-S6B**).

**Figure 4.**
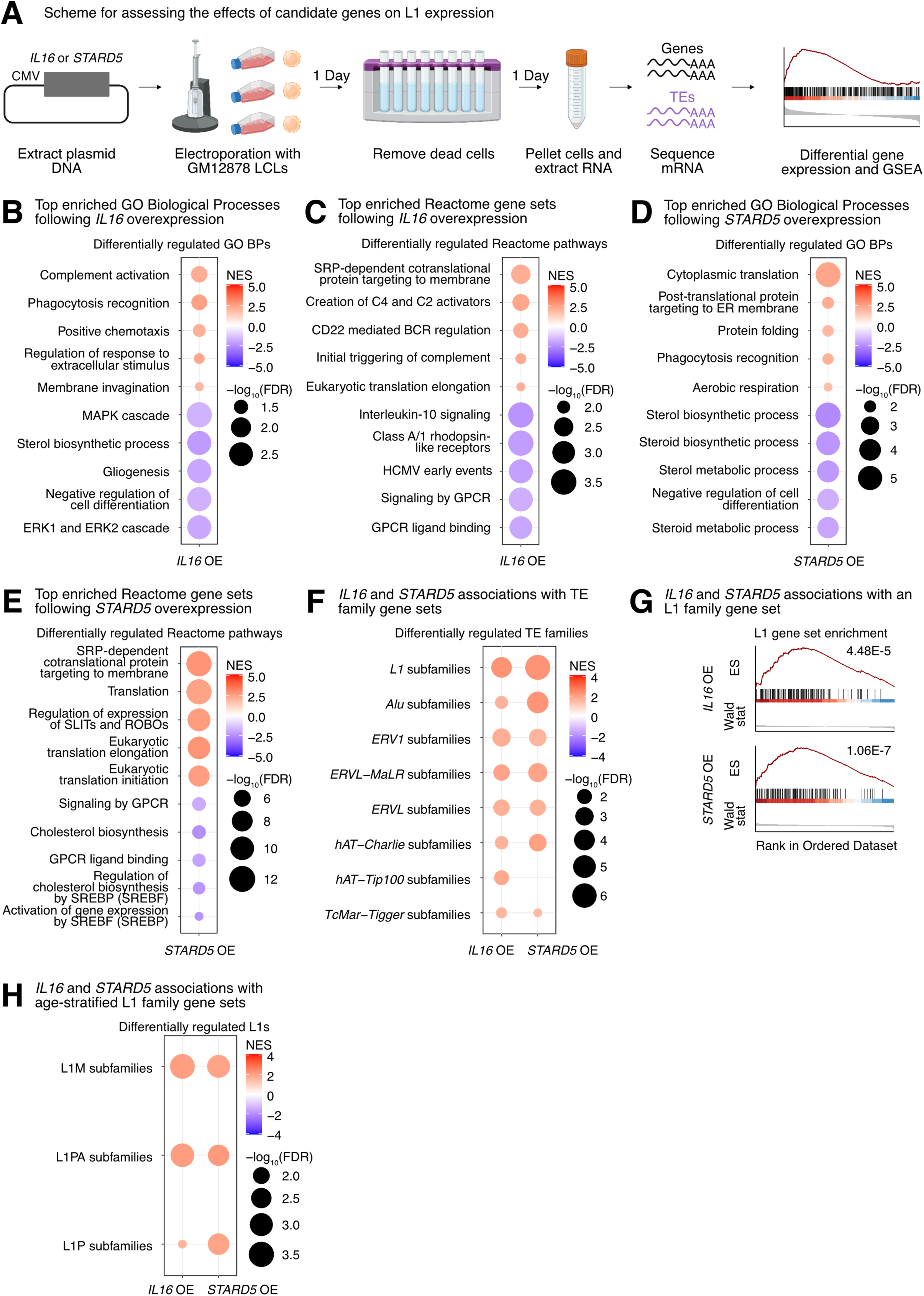
Impact of *IL16* and *STARD5* overexpression on LCL gene and TE expression landscapes. *IL16* and *STARD5* overexpression induce changes consistent with their known biology, as well as subtle but widespread upregulation of TE families. **(A)** Scheme for experimentally validating the roles of *IL16* and *STARD5* in L1 regulation. GSEA analysis for top, differentially regulated **(B)** GO Biological Process and **(C)** Reactome pathway gene sets following *IL16* overexpression. GSEA analysis for top, differentially regulated **(D)** GO Biological Process and **(E)** Reactome pathway gene sets following *STARD5* overexpression. **(F)** GSEA analysis for shared, significantly regulated TE family gene sets following *IL16* and *STARD5* overexpression. **(G)** GSEA plots for the L1 family gene set results summarized in **(F)**. For these plots, the FDR value is listed. **(H)** GSEA analysis for shared, significantly regulated, evolutionary-age-stratified L1 gene sets across *IL16* and *STARD5* overexpression. L1M subfamilies are the oldest, L1P subfamilies are intermediate, and L1PA subfamilies are the youngest. In each bubble plot, the size of the dot represents the -log_10_(FDR) and the color reflects the normalized enrichment score. FDR: False Discovery Rate. Some panels were created with BioRender.com.

To further assess the biological relevance of each overexpression, we carried out GSEA using the GO Biological Process, Reactome pathway, and Hallmark pathway gene sets (**Supplementary Table S6E-S6P**). Importantly, GSEA using GO Biological Process and Reactome pathway gene sets highlighted differences that were consistent with the known biology of our candidate genes. Firstly, *IL16* is involved in regulating T-cell activation, B-cell differentiation, and functions as a chemoattractant (Center and Cruikshank 1982; Cruikshank et al. 1994; Center et al. 1996; Cruikshank et al. 1996; Theodore et al. 1996; Wilson et al. 2004). Moreover, it modulates macrophage polarization by regulating *IL-10* expression (Huang et al. 2019). *IL16* overexpressing cells showed upregulation for “phagocytosis recognition” and “positive chemotaxis”, downregulation for “negative regulation of cell differentiation”, and downregulation for “Interleukin 10 signaling” (**Figure 4B-4C**). Secondly, *STARD5* encodes a cholesterol transporter and is upregulated in response to endoplasmic reticulum (ER) stress (Soccio et al. 2005; Rodriguez-Agudo et al. 2012; Rodriguez-Agudo et al. 2019). *STARD5* overexpressing cells showed downregulation of various cholesterol-related gene sets such as “sterol biosynthetic process”, “sterol metabolic process”, and “regulation of cholesterol biosynthesis by SREBP (SREBF)” (**Figure 4D-4E**). Thirdly, *HSD17B12* encodes a steroid dehydrogenase involved in converting estrone into estradiol and is essential for proper lipid homeostasis (Luu-The et al. 2006; Nagasaki et al. 2009; Heikelä et al. 2020). *HSD17B12* overexpressing cells showed downregulation of cholesterol-related gene sets, including “sterol biosynthetic process” and “sterol metabolic process” (**Supplementary Figure S14E**). Finally, *RNF5* encodes an ER and mitochondrial-bound E3 ubiquitin-protein ligase that ubiquitin-tags proteins for degradation (Didier et al. 2003; Tcherpakov et al. 2009; Zhong et al. 2009; Zhong et al. 2010). *RNF5* overexpressing cells demonstrated alterations in gene sets involved in proteostasis and ER biology, including upregulation of “ERAD pathway” and “response to endoplasmic reticulum stress” (**Supplementary Figure S14F**). These results suggest that our approach leads to biological changes consistent with the known biological impact of the genes being overexpressed.

Next, we sought to determine whether modulation of candidate genes had any impact on TE RNA levels in general, and L1 in particular. Although there were no significant changes for individual TE subfamilies following *IL16* and *STARD5* overexpression (**Supplementary Table S6A-S6B**), we identified subtle but widespread upregulation of various TE families across both conditions by GSEA (**Figure 4F, Supplementary Table S6Q-S6R**). Interestingly, 7 families, including L1, ERV1, ERVL-MaLR, Alu, ERVL, TcMar-Tigger, and hAT-Charlie families, were commonly upregulated under both conditions (**Figure 4F**). In contrast, cells overexpressing *HSD17B12* or *RNF5* did not drive widespread changes in L1 family expression as assessed by GSEA (**Supplementary Table S6S-S6T**), suggesting specificity of the *IL16/STARD5*-L1 relationships. Noteworthy, the L1 family gene set was more significantly upregulated following *STARD5* overexpression (NES = 2.27, FDR = 1.06E-7) compared to *IL16* overexpression (NES = 2.27, FDR = 4.48E-5) (**Figure 4G, Supplementary Table S6Q-S6R**). Since *IL16* is upregulated in response to *STARD5* overexpression, this suggests that *STARD5* may synergize with *IL16* for the regulation of L1 RNA levels.

Overall, we observed similar changes in TE RNA levels when we applied the alternative one-sample Wilcoxon test approach to determine whether TE family changes were significantly different than 0 across overexpression conditions (**Supplementary Figure S15A-B**). Among L1 changes, older (L1M), intermediate (L1P), and younger (L1PA) elements were differentially regulated across both overexpression conditions (**Figure 4H**). To gain insight into the mechanism of *IL16/STARD5*-mediated TE mis-regulation, we again stratified the TE expression profiles by genomic origin (**Supplementary Table S6U-S6V**). Though intronic TEs of various families were strongly upregulated following *IL16* and *STARD5* overexpression, distal intergenic L1 RNA levels were also upregulated in both conditions (**Supplementary Figure S15C-D**). These results further suggest that *IL16* and *STARD5* influence the repetitive RNA pools, including elements that are unlikely to be transcribed by neighboring genes.

Then, we decided to further characterize the impact of IL16 activity on TEs, since (i) its overexpression led to a global upregulation of TE transcription, and (ii) it was itself upregulated in response to *STARD5* overexpression, which also led to increased TE expression. Thus, since IL16 is a soluble cytokine, we independently assessed its regulatory properties by exposing GM12878 cells to recombinant human IL16 peptide [rhIL16] for 24 hours (**Figure 5A, Supplementary Figure S16A**). Differential gene expression analysis (**Supplementary Table S7A**) and comparison with the *IL16* overexpression results demonstrated that differentially expressed genes were weakly but significantly correlated (**Supplementary Figure S16B**). As with the overexpression conditions, no significant differences in EBV expression were identified (FDR > 0.05; **Supplementary Figure S16C**). Additionally, we carried out GSEA using the GO Biological Process, Reactome pathway, and Hallmark pathway gene sets (**Supplementary Table S7B-S7E**) and compared those results with the GSEA from the *IL16* overexpression (**Supplementary Table S7F-S7H**). Consistent with the known biology of *IL16*, GSEA highlighted a downregulation of many immune cell-related gene sets such as “leukocyte differentiation” and “mononuclear cell differentiation” (**Figure 5B-5C, Supplementary Table S7F-S7H**). Similar to our overexpression results, exposure of GM12878 to rhIL16 for 24 hours led to the upregulation of an L1 family gene set by GSEA, although the effect was less pronounced than with the overexpression (**Figure 5D**). Again, we observed similar changes in TE RNA levels when we applied the one-sample Wilcoxon test alternative approach to determine whether TE family changes were significantly different than 0 following rhIL16 treatment for 24 hours (**Supplementary Figure S16D**). Similar to the overexpression, intermediate (L1P) and younger (L1PA) age L1 elements were upregulated following rhIL16 treatment for 24 hours (**Figure 5E, Supplementary Table S7J**). After stratifying the TE RNA profiles by genomic origin and running GSEA, we again observed an upregulation of intronic and distal intergenic L1 RNA, as in the overexpression (**Figure 5F, Supplementary Table S7I**). Even though treatment of GM12878 with rhIL16 for 48 hours exhibited known features of IL16 biology (**Supplementary Figure S16B, S16E-F, Supplementary Table S7K-S7S**), the L1 upregulation was no longer detectable, though other TEs remained upregulated (**Supplementary Figure S16G, Supplementary Table S7S**). These results further support the notion that *IL16* acts as a modulator of L1 RNA levels, including for both intronic and distal intergenic copies.

**Figure 5.**
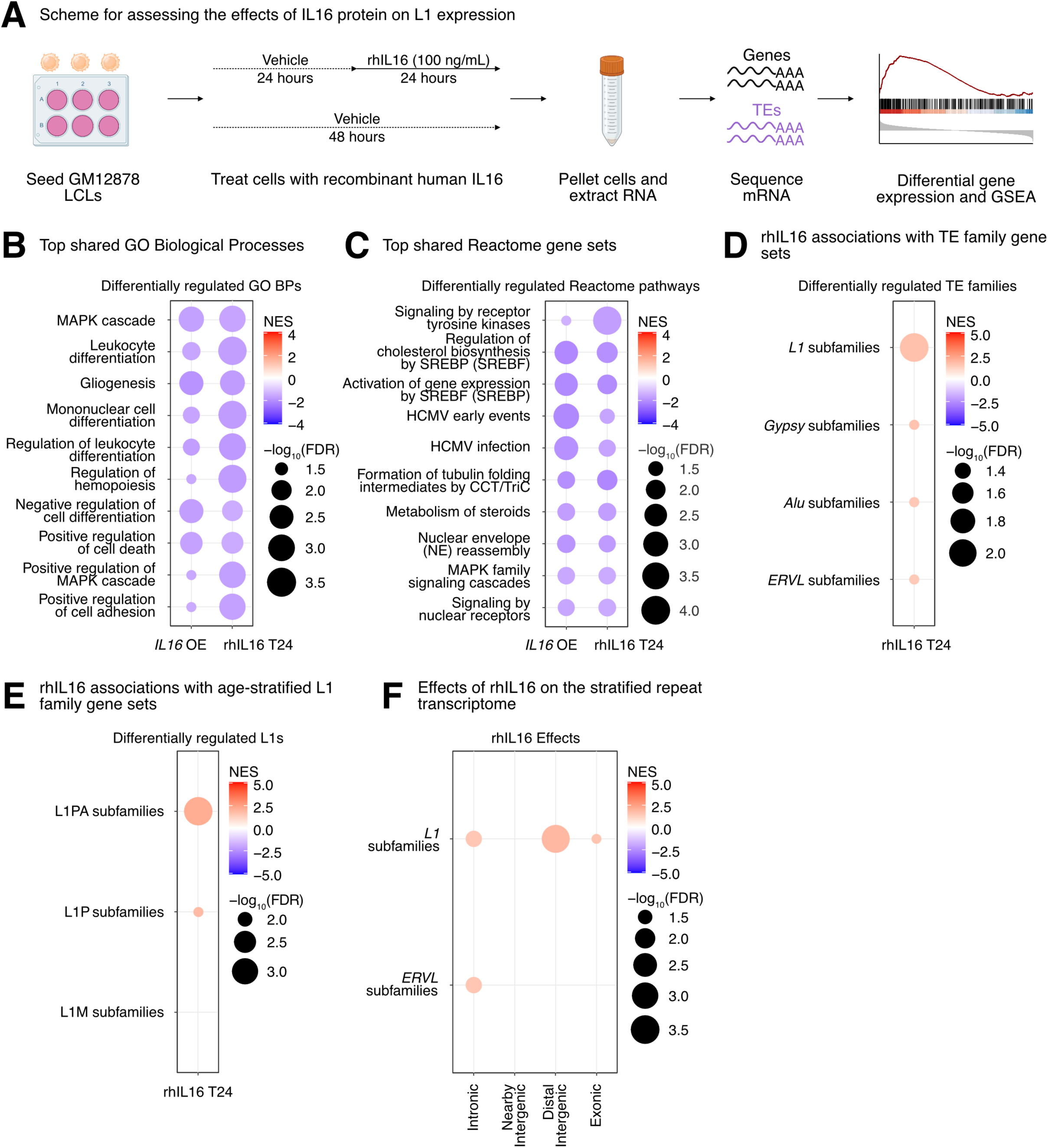
rhIL16 treatment is sufficient to transiently upregulate an L1 family gene set. **(A)** Scheme for experimentally validating the role of rhIL16 in L1 regulation. GSEA analysis for top, shared, concomitantly regulated **(B)** GO Biological Process and **(C)** Reactome pathway gene sets following *IL16* overexpression and rhIL16 exposure for 24 hours. Shared gene sets were ranked by combining p-values from each individual treatment analysis using Fisher’s method. **(D)** GSEA analysis for top, differentially regulated TE family gene sets following rhIL16 exposure for 24 hours. **(E)** GSEA analysis for significantly regulated evolutionary-age-stratified L1 gene sets following rhIL16 exposure. L1M subfamilies are the oldest, L1P subfamilies are intermediate, and L1PA subfamilies are the youngest. **(F)** GSEA analysis for top, differentially regulated TE family gene sets in different genomic locations following rhIL16 exposure for 24 hours. In each bubble plot, the size of the dot represents the -log_10_(FDR) and the color reflects the normalized enrichment score. FDR: False Discovery Rate. Some panels were created with BioRender.com.

Finally, we sought to define the biological pathways regulated concomitantly with the L1 family gene set under all experimental conditions where it was upregulated (i.e., *IL16* overexpression, *STARD5* overexpression, and 24 hours of rhIL16 exposure) (**Figure 6A**, **Figure 6B, Supplementary Table S8A**). Again, we reasoned that such pathways would act either upstream (as regulatory pathways) or downstream (as response pathways) of TE alterations. GSEA with the Hallmark pathway gene sets identified 7 gene sets fitting this criterion, including “TNFα signaling via NF-κB”, “IL2 STAT5 signaling”, “inflammatory response”, “mTORC1 signaling”, “estrogen response early”, “apoptosis”, and “UV response up” (**Figure 6C, Supplementary Table S7B**). GSEA with the GO Biological Process gene sets (**Figure 6D, Supplementary Table S7C**) and the Reactome pathway gene sets (**Figure 6E, Supplementary Table S7D**) also identified MAPK signaling, virus-related pathways like “HCMV early events”, pathways involved in cell differentiation, and pathways involved in cholesterol and steroid metabolism like “signaling by nuclear receptors”. These results further cement the catalogue of pathways associated with differences in TE RNA levels.

**Figure 6.**
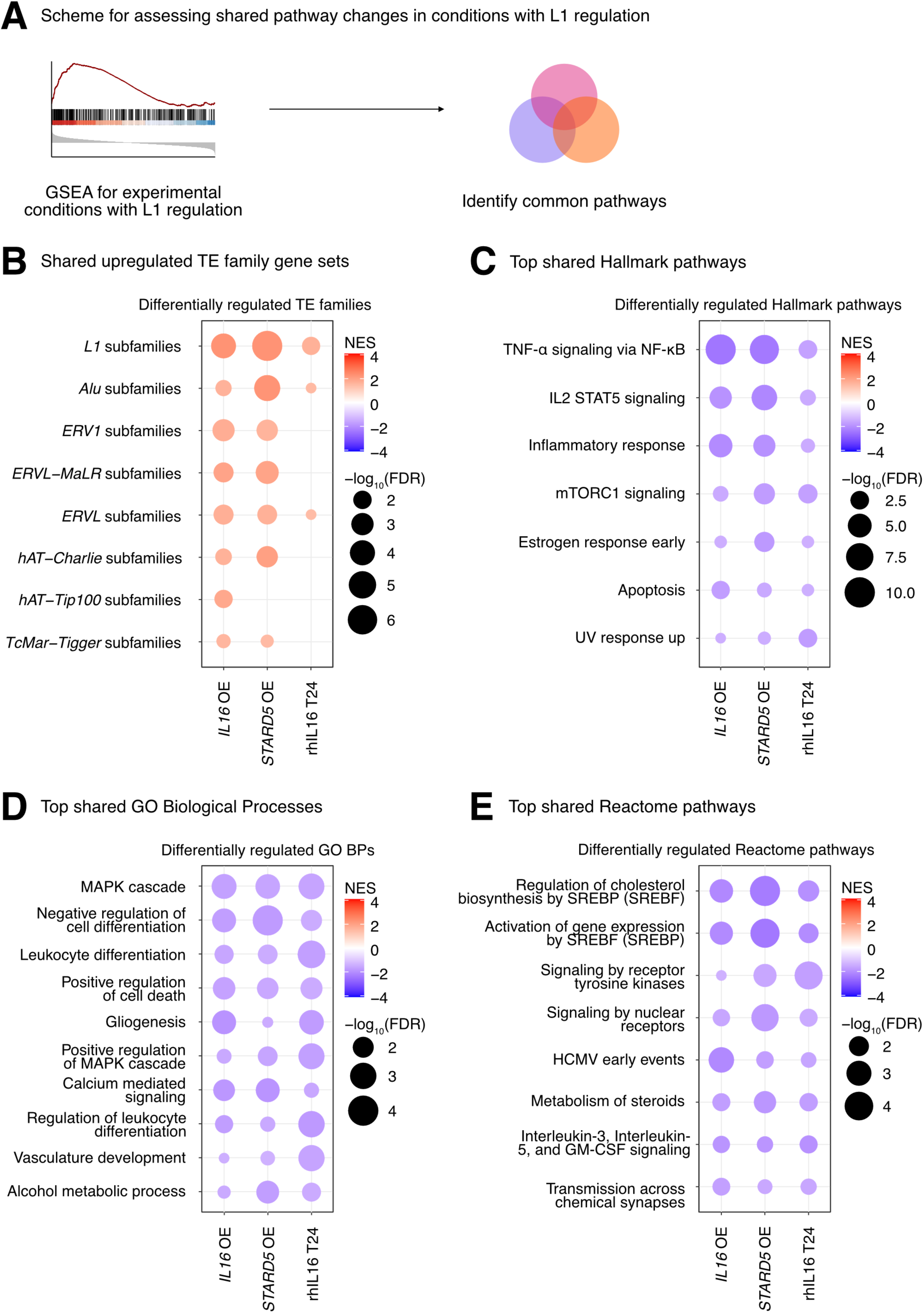
Consistent cellular responses to *IL16* overexpression, *STARD5* overexpression, and rhIL16 exposure for 24 hours. *IL16* overexpression, *STARD5* overexpression, and rhIL16 exposure for 24 hours are associated with subtle but widespread differences in TE families and known TE-associated pathways. **(A)** Scheme for assessing concordantly regulated TE family and pathway gene sets across conditions where an L1 gene set is upregulated. GSEA analysis for top, shared, concomitantly regulated **(B)** TE family, **(C)** MSigDB Hallmark pathway, **(D)** GO Biological Process, and **(E)** Reactome pathway gene sets following *IL16* overexpression, *STARD5* overexpression, and rhIL16 exposure for 24 hours. Shared gene sets were ranked by combining p-values from each individual treatment analysis using Fisher’s method. In each bubble plot, the size of the dot represents the - log_10_(FDR) and the color reflects the normalized enrichment score. FDR: False Discovery Rate.

### L1 trans-eQTLs are co-associated with aging traits in GWAS databases

Although TE de-repression has been observed broadly with aging and age-related disease (Lai et al. 2019; Bravo et al. 2020), whether this de-repression acts as a causal driver, or a downstream consequence, of aging phenotypes remains unknown. We reasoned that if increased TE expression at least partially drives aging phenotypes, L1 *trans*-eQTLs should be enriched for associations to aging traits in genome-wide association studies [GWAS] or phenome-wide association studies [PheWAS].

To test our hypothesis, we queried the Open Targets Genetics platform with our initial 499 *trans*-eQTL SNVs, mapped traits to standardized MeSH IDs, and then manually curated MeSH IDs related to aging-related traits (**Figure 7A**). Consistent with our hypothesis, a large proportion of L1 *trans*-eQTL SNVs (222/499 or 44.5%) were either (i) associated with an aging MeSH trait by PheWAS or (ii) LD-linked to a lead variant associated with an aging MeSH trait (**Figure 7B**). Moreover, among the 222 SNVs with significant aging-trait associations, we observed frequent mapping to more than a single age-related trait by PheWAS, with many SNVs associated with 10-25 traits (**Figure 7C, Supplementary Table S9A**). Additionally, many of the 222 SNVs mapped to 1-5 aging traits through a proxy lead variant (**Figure 7D, Supplementary Table S9A**). Among the most frequently associated or linked traits, we identified type 2 diabetes mellitus, hyperparathyroidism, thyroid diseases, coronary artery disease, hypothyroidism, and psoriasis, among many others (**Figure 7E, Supplementary Table S9B**).

**Figure 7.**
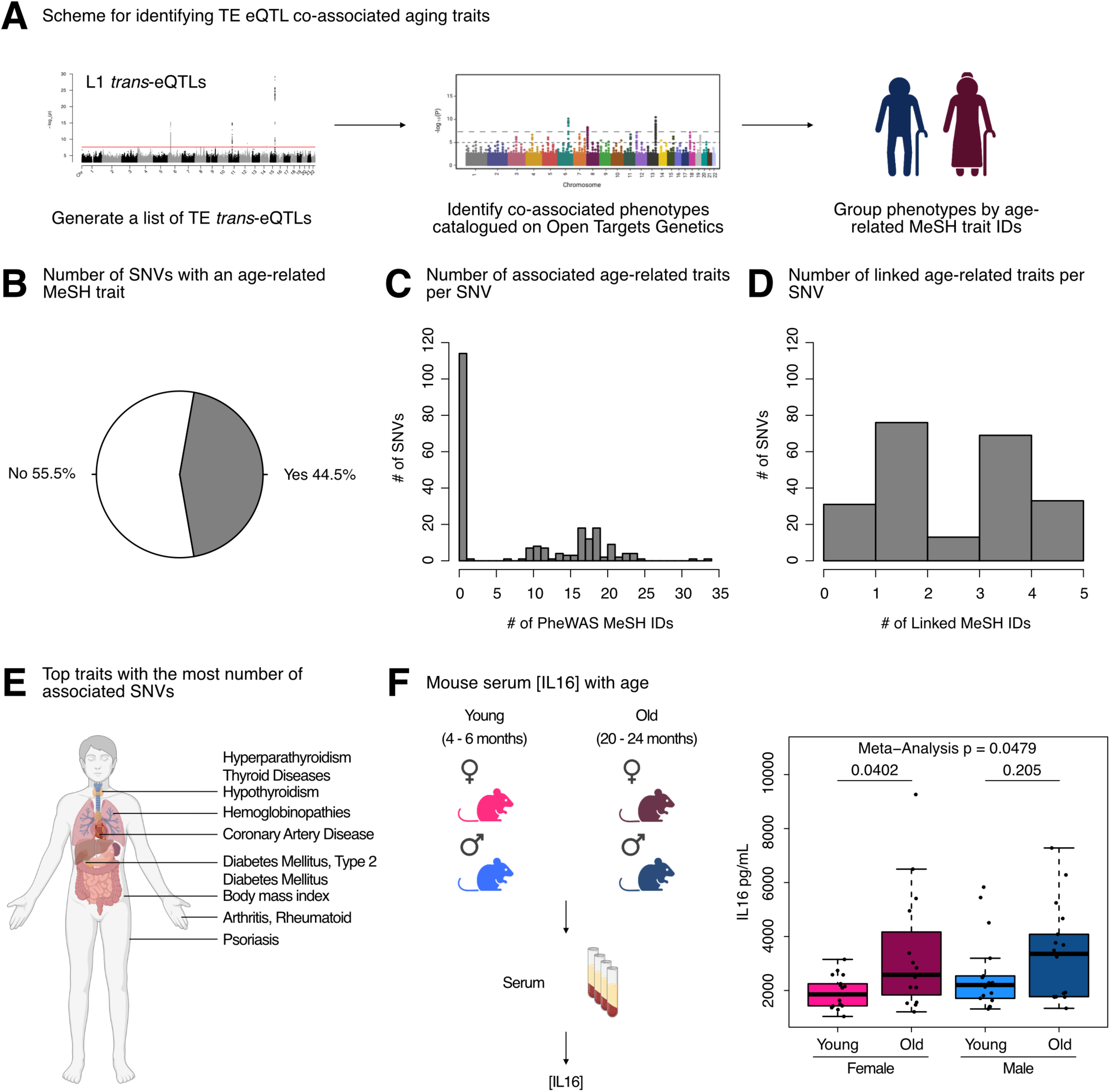
L1 *trans*-eQTLs are co-associated with aging traits in GWAS databases. **(A)** Scheme for obtaining *trans*-eQTL SNV-associated aging phenotypes from the Open Targets Genetics platform. **(B)** A pie chart representing the number of SNVs (222/499) associated with an aging-related MeSH trait, either by PheWAS or indirectly linked to the phenotype through a proxy lead SNP in LD with the SNV. **(C)** Histogram depicting the distribution of number of aging MeSH traits associated with the 222/499 SNVs by PheWAS. **(D)** Histogram depicting the distribution of number of aging MeSH traits linked with the 222/499 SNVs through a proxy lead SNP in LD with the SNVs. **(E)** A diagram highlighting the organ targets of the top 10 most frequently associated aging traits. **(F)** The concentrations of circulating IL16 in aging mice of both sexes was assessed by ELISA. Each dot represents an independent animal, with n = 15 - 17 per group. Significance across age in each sex was assessed using a Wilcoxon test. The p-values from each sex (females in pink and males in blue) were combined by meta-analysis using Fisher’s method. Any p-value < 0.05 was considered significant. Some panels were created with BioRender.com.

As a parallel approach, we queried the Open Targets Genetics platform with our L1 *trans*-eQTL SNVs, as well as 500 combinations of random SNVs sampled from all SNVs used in the eQTL analyses. We then leveraged broader phenotype categories annotated by the platform, including 14 disease categories that we considered aging-related, to determine whether L1 eQTL associations were enriched for any disease categories (**Supplementary Figure S17A**). L1 eQTL associations were significantly enriched (FDR < 0.05 and ES > 1) for 13 out of 14 disease categories, including cell proliferation disorders, immune system diseases, and musculoskeletal diseases (**Supplementary Figure S17B-N**). The cardiovascular diseases category was the only disease category for which we did not observe a significant enrichment (**Supplementary Figure S17O**). The enrichment for cell proliferation disorders is consistent with the associations of L1 activity with cellular senescence (De Cecco et al. 2013a; De Cecco et al. 2019) and cancer (Rodić et al. 2014; Sato et al. 2023). The enrichment for immune system diseases is consistent with the role of L1 as a stimulator of the interferon pathway, inflammation, and senescence (De Cecco et al. 2019), as well as the more general notion that transposons can mimic viruses and stimulate immune responses from their hosts (Lindholm et al. 2023). The enrichment for musculoskeletal diseases is consistent with an increase in L1 expression and copy number with age in muscle tissue from aging mice (De Cecco et al. 2013b). These results reinforce the notion that L1 activity is strongly and non-randomly associated with an assortment of age-related diseases.

Intriguingly, a large fraction of co-associated SNVs were on chromosome 6 near the HLA locus, which has previously been shown to be a hotspot of age-related disease traits (Jeck et al. 2012). Despite its association to our strongest L1 trans-eQTL SNV, little is known about the regulation and impact of IL16 during aging. One study, however, found that *IL16* expression increases with age in ovarian tissue, and the frequency of *IL16* expressing cells is significantly higher in ovarian tissue from women at early and late menopause, compared to premenopausal women (Ramirez et al. 2022). Given these findings, and since L1 expression levels and copy number have been found to increase with age [reviewed in (Bravo et al. 2020)], we asked whether circulating IL16 levels may also change with age, using C57BL/6JNia mice as a model (**Figure 7F, Supplementary Table S9C**). Consistent with the notion that increased IL16 levels may, at least partially, drive age-related TE de-repression, we observed a significant increase in circulating IL16 levels in female mice with age, and a trending increase with age in male mice (although the levels showed more animal-to-animal variability). By meta-analysis, circulating IL16 levels changed significantly with age across sexes (**Figure 7F**). These results further support the hypothesis that *IL16* is involved in L1 biology and may modulate L1 age-related changes. In sum, our results provide one of the first pieces of evidence of a causal link between L1 RNA levels and age-related decline.

## Discussion

### A new approach to identify regulators of TE expression

In this work, we developed a pipeline to computationally identify candidate L1 RNA level regulators by eQTL analysis. We provide experimental evidence for the involvement of top candidates in regulating L1 RNA levels, demonstrating as a proof-of-principle that this approach can be broadly used on other large “omic”-characterized cohorts with human (i.e. GTEx (Lonsdale et al. 2013; Carithers et al. 2015) or HipSci (Streeter et al. 2016)) or mouse (i.e. DO mice (Chick et al. 2016)) subjects to identify other regulators of L1 activity. These datasets, combined with our approach, could be utilized to rigorously characterize conserved or group-specific TE regulatory mechanisms on multiple layers, such as across TE families (like Alu or ERVs), across cell or tissue types, across ancestry groups, and across species. This approach, which leverages existing datasets to perform *in silico* screening, could be a powerful method to expand our knowledge of TE regulation in non-diseased cells and tissues.

Though our initial scan identified genetic variants associated with expression differences in specific L1 subfamilies, secondary analyses by GSEA suggest that genetic variants are associated with subtle but global differences in the expression of many TE families of varying genomic context, including intronic, intergenic, and exonic TE RNA levels. Our pipeline identified candidate genes, including *HSD17B12* and *HLA* genes, that likely play a conserved role in L1 regulation across human populations of different ancestries. Though some top candidates from the European cohort scan, such as *IL16, STARD5*, and *RNF5*, were not significant in the African cohort analysis, it is likely that some of these genes would appear in cross-ancestry scans with larger samples sizes. To note, none of our top candidates were associated with L1 polymorphisms in the landmark TE-eQTL study that first used this data to study TE biology (Wang et al. 2016), suggesting that our findings are likely mechanistically independent.

After repeating our computational scan using the TE profiles stratified by genomic context, we made a number of additional insights, such as (i) the association of *IL16* and *STARD5* with intronic L1 RNA levels, (ii) the association of *HLA* with nearby intergenic L1 RNA levels, (iii) the identification of an additional candidate regulator, *ZSCAN26*, which may influence distal intergenic L1 RNA levels, and (iv) a number of SNVs associated with exon-overlapping L1 RNA levels. Moreover, the secondary GSEA analyses suggest that the SNVs tested exert broad effects on TE family RNA levels, regardless of genomic context.

As an important aspect of this study, we experimentally validated our top candidate genes. We detected subtle but global differences in L1 family RNA levels following *IL16* overexpression, *STARD5* overexpression, and rhIL16 treatment for 24 hours, further suggesting that our top candidates have regulatory potential. Although *IL16/STARD5* were mainly associated to intronic L1 levels in our trans-eQTL analysis, these treatments affected both intronic and distal intergenic L1 RNA levels, suggesting that these differences are not solely due to intron retention or co-expression with neighboring genes. Surprisingly, our treatments exerted effects on other TE families as well, suggesting broad alterations that promote TE RNA differences. Thus, *IL16* and *STARD5* are likely to be *bona fide* regulators that can be prioritized for follow-up study.

Much of the work on L1 regulation relies on overexpressing full-length L1 elements in cell lines that can tolerate these manipulations (Liu et al. 2018; Mita et al. 2020). However, there are some approaches aimed at characterizing transcription factors that bind endogenous L1 promoters (Sun et al. 2018; Briggs et al. 2021), in addition to those that implement gene network approaches to find potential regulators that may act through alternative mechanisms (Chung et al. 2019). These, in our view, complement plasmid-based approaches through a more physiological study of L1 regulators. We think that our approach, which relies on endogenous TE profiles, adds to this category of tools.

### New candidate L1 regulators are involved in viral response

As another, theoretical line of evidence for the potential involvement of our top candidate genes in L1 regulation, we highlight known interactions between tested candidate genes and viral infections, which may be relevant under conditions where transposons are recognized as viral mimics (Lindholm et al. 2023). Indeed, *IL16* has been extensively studied for its ability to inhibit human immunodeficiency virus (HIV) replication, partly by suppressing mRNA expression (Baier et al. 1995; Zhou et al. 1997; Idziorek et al. 1998). Additionally, but in contrast to its HIV-suppressive properties, *IL16* can enhance the replication of influenza A virus (IAV) and facilitate its infection of hosts, potentially through its repression of type I interferon beta and interferon-stimulated genes (Jia et al. 2021). *IL16* can also contribute to the establishment of lifelong gamma herpesvirus infection (Liu et al. 2020). *STARD5* is another candidate implicated in the influenza virus replication cycle (Watanabe et al. 2010). *HSD17B12* promotes the replication of hepatitis C virus via the very-long-chain fatty acid (VLCFA) synthesis pathway and the production of lipid droplets important for virus assembly (Germain et al. 2014; Mohamed et al. 2020). Additionally, HSD17B12 has been found interacting with the coronavirus disease 2019 (COVID-19) protein nonstructural protein 13 (NSP13), which is thought to antagonize interferon signaling (Feng et al. 2021). Finally, *RNF5* has been implicated in both promoting and antagonizing severe acute respiratory syndrome coronavirus 2 (SARS-CoV-2) by either stabilizing the interactions of membrane protein (M) (Yuan et al. 2022) or inducing degradation of structural protein envelope (E) (Li et al. 2023), respectively. Fundamentally, *RNF5* regulates virus-triggered interferon signaling by targeting the stimulator of interferon genes (STING) or mitochondrial antiviral signaling protein (MAVS) for ubiquitin-mediated protein degradation (Zhong et al. 2009; Zhong et al. 2010). These studies reinforce the roles of tested candidate regulators in virus-associated processes, including interferon-mediated signaling.

### Future considerations for the use of trans-eQTL analysis in identification of L1 regulators

While we believe this approach can readily be applied to other datasets, we would like to note potential further considerations with the approach implemented here, some of which were simply beyond the scope of this paper. Firstly, though it is common to use probabilistic estimation of expression residuals (PEER) (Stegle et al. 2012) to enhance detection of *cis*-eQTLs, PEER was not implemented in our analysis as a precautionary measure, in order to avoid potentially blurring global TE signals, which likely led to a more conservative list of candidate *cis* gene mediators. Second, given the technical complexity in generating the vast amount of mRNA-seq data used for the eQTL analysis, it is possible that technical covariates introduced non-linear effects that would not be easily removed by approaches like PEER or SVA (Leek 2014). For that reason, we opted to supplement our computational predictions with experimental data. Third, the L1 *trans*-eQTLs identified were specific to older L1 subfamilies (L1P and L1M) and were not shared across subfamilies. One factor that may partially explain this is the heightened difficulty of quantifying the expression of evolutionarily younger L1 subfamilies using short-read sequencing (Savytska et al. 2022).

More generally, significant single gene differences are often difficult to reproduce across studies, and it is for this reason that methods like GSEA were developed, to robustly identify broader changes in sets of genes (Subramanian et al. 2005). Consistently, GSEA suggests that many TE families, beyond the single L1 subfamilies identified in the eQTL analysis, are differentially regulated among samples with different genotypes for *trans*-eQTL SNVs and among samples where *IL16/*IL16 and *STARD5* were manipulated. We note that although *HLA* and *HSD17B12* loci were significant in both the European and African cohorts, we were not able to independently identify all of the same candidate regulators. This is likely due to a combination of small sample size for the African cohort and the existence of population-specific L1 regulation. Future studies with larger sample sizes may be useful for expanding the catalogue of loci that are biologically meaningful for L1 expression across more than one population. Importantly, our computational scan is limited to loci exhibiting genomic variation among tested individuals. This will vary with factors like the ancestry groups of the populations being studied. Moreover, variants that confer extreme fitness defects may not exist at a sufficiently high level in a population to allow for an assessment of their involvement as eQTLs.

Finally, although we focused on protein-coding candidate regulators, it is possible that the non-coding genes identified in our scan may also causally drive differences in L1 expression. Though not explored here, other regulatory factors like small RNAs may also act as partial mediators. Since the GEUVADIS Consortium generated small RNA data in parallel to the mRNA data used in this study (Lappalainen et al. 2013), a future application of our pipeline could be to scan for *cis* small RNA mediators in the same biological samples. These unexplored factors may explain the associations between orphan SNV genotypes and TE family gene set changes.

### L1 trans-eQTLs are enriched for genetic variants linked to aging and age-related disease

Consistent with the notion that L1 is associated with aging and aging phenotypes (Lai et al. 2019; Bravo et al. 2020), we observed that L1 *trans*-eQTL SNVs were associated with aging phenotypes in GWAS/PheWAS databases. This is very surprising, but interesting, given that all 1000Genomes Project participants declared themselves to be healthy at the time of sample collection. Assuming this to be true, our results suggest that L1 RNA level differences exist in natural, healthy human populations, and these RNA level differences precede onset of aging diseases. Importantly, we note that the SNVs tested were associated with intronic and nearby intergenic L1 subfamily RNA levels, which may have been discarded in studies focusing on full-length intergenic L1 elements. Thus, these results reiterate the notion that intronic L1s and intergenic L1s near genes can potentially exert functional consequences on hosts and therefore merit further study. Though it is often unclear whether L1 mis-regulation is a consequence or driver of aging phenotypes, our results suggest that L1 RNA levels may drive aging phenotypes. As we continue to expand the catalogue of L1 regulators, especially in healthy cells and tissues, the L1 regulatory processes that are disrupted over the course of aging will become increasingly clear. To that end, this work may serve as a guide for conducting more comprehensive scans for candidate TE regulators.

In summary, we developed an eQTL-based pipeline that leverages genomic and transcriptomic data to scan the human genome for novel candidate regulators of L1 subfamily RNA levels. Though the initial scan identified genetic variants associated with RNA level differences in specific L1 subfamilies, secondary analyses by GSEA suggest that genetic variants are associated with subtle but global differences in the RNA levels of many TE families. Our pipeline identified candidate genes, including *HSD17B12* and *HLA* genes, that likely play a conserved role in L1 regulation across human populations of different ancestries. Though some top candidates from the European cohort scan, such as *IL16, STARD5*, and *RNF5*, were not significant in the African cohort analysis, it is likely that some of these genes would appear in cross-ancestry scans with larger samples sizes. We detected subtle but global differences in L1 family RNA levels following *IL16* overexpression, *STARD5* overexpression, and rhIL16 treatment for 24 hours, further suggesting that some candidate genes have regulatory potential. We generate a list of pathways, such as mTORC1 signaling and cholesterol metabolism, that may act upstream of L1 regulation. Finally, the co-association of some genetic variants with both L1 RNA level differences and various age-related diseases suggests that L1 differences may precede and contribute to the onset of disease. Our results expand the potential mechanisms by which L1 RNA levels are regulated and by which L1 may influence aging-related phenotypes.

## Material and Methods

### Publicly available data acquisition

The eQTL analysis was carried out on 358 European (EUR) individuals and 86 Yoruban (YRI) individuals for which paired single nucleotide variant, structural variant, and transcriptomic data were available from Phase 3 of the 1000 Genomes Project (Auton et al. 2015; Sudmant et al. 2015) and from the GEUVADIS consortium (Lappalainen et al. 2013). Specifically, Phase 3 autosomal SNVs called on the GRCh38 reference genome were obtained from The International Genome Sample Resource (IGSR) FTP site (http://ftp.1000genomes.ebi.ac.uk/vol1/ftp/data_collections/1000_genomes_project/release/20190312_biallelic_SNV_and_INDEL/). Structural variants were also obtained from the IGSR FTP site (http://ftp.1000genomes.ebi.ac.uk/vol1/ftp/phase3/integrated_sv_map/). mRNA-sequencing fastq files generated by the GEUVADIS consortium were obtained from ArrayExpress under accession E-GEUV-1.

### Aggregating and pre-processing genotype data for eQTL analyses

To prepare SNVs for association analyses, all SNVs were first annotated with rsIDs from dbSNP build 155 using BCFtools v1.10.2 (Danecek et al. 2021). VCFtools v0.1.17 (Danecek et al. 2011) was then used to remove indels and keep variants with the following properties in each of the two populations: possessed a minimum and maximum of two alleles, possessed a minor allele frequency (MAF) of at least 1%, passed Hardy-Weinberg equilibrium thresholding at p < 1e-6, with no missing samples, and located on an autosome. We note here that sex chromosomes were not included in the analysis since (i) Y chromosome SNVs were not available and (ii) analyses with X chromosome SNVs require unique algorithms and cannot simply be incorporated into traditional association pipelines (Gao et al. 2015; Keur et al. 2022). VCF files containing these filtered SNVs were then converted to PLINK BED format using PLINK v1.90b6.17 (Purcell et al. 2007), keeping the allele order. PLINK BED files were subsequently used to generate preliminary 0/1/2 genotype matrices using the ‘--recodeA’ flag in PLINK. These preliminary matrices were manipulated in terminal, using the gcut v9.0 function to remove unnecessary columns and datamash v1.7 to transpose the data, to generate the final 0/1/2 matrices used for the eQTL analyses. Finally, PLINK was used to prune the list of filtered SNVs, using the “--indep-pairwise 50 10 0.1” flag, and to generate principal components (PCs) from the pruned genotypes.

To control for inter-individual differences in genomic transposon copy number load, we applied 1 of 2 approaches, depending on the analysis. For approach 1, the net number of L1 and Alu insertions was quantified across the 444 samples. We chose to aggregate the L1 and Alu copy numbers, since Alu relies on L1 machinery for mobilization (Ahl et al. 2015), and so the aggregate number may provide a finer view of L1-associated copy number load. Briefly, VCFTools was used to extract autosomal structural variants from the 1000Genomes structural variant calls. L1 and Alu insertions and deletions were then extracted with BCFtools by keeping entries with the following expressions: ‘SVTYPE=“LINE1”’, ‘SVTYPE=“ALU”’, ‘SVTYPE=“DEL_LINE1”’, and ‘SVTYPE=“DEL_ALU”’. The resulting VCF files were then transformed to 0/1/2 matrices in the same manner as the SNVs. A net copy number score was obtained for each sample by adding the values for the L1 and Alu insertions and subtracting the values for the L1 and Alu deletions. For approach 2, the complete structural variant matrix was filtered with VCFtools using the same parameters as with the SNV matrices. The filtered structural variant matrix was then pruned with PLINK, and these pruned structural variant genotypes were used to generate principal components, in the same fashion as with the SNV matrix. The net copy number score or the structural variant principal components, depending on the analysis, were included as covariates.

### mRNA-seq read trimming, mapping, and quantification

Fastq files were first trimmed using fastp v0.20.1 (Chen et al. 2018) with the following parameters: detect_adapter_for_pe, disable_quality_filtering, trim_front1 17, trim_front2 17, cut_front, cut_front_window_size 1, cut_front_mean_quality 20, cut_tail, cut_tail_window_size 1, cut_tail_mean_quality 20, cut_right, cut_right_window_size 5, cut_right_mean_quality 20, and length_required 36. Read quality was then inspected using fastqc v0.11.9.

Next, the GRCh38 primary human genome assembly and comprehensive gene annotation were obtained from GENCODE release 33 (Frankish et al. 2018). Since LCLs are generated by infecting B-cells with Epstein-Barr virus, the EBV genome (GenBank ID V01555.2) was included as an additional contig in the human reference genome. The trimmed reads were aligned to this modified reference genome using STAR v2.7.3a (Dobin et al. 2012) with the following parameters: outFilterMultimapNmax 100, winAnchorMultimapNmax 100, and outFilterMismatchNoverLmax 0.04. The TEcount function in the TEtranscripts v2.1.4 (Jin et al. 2015) package was employed to obtain gene and TE counts, using the GENCODE annotations to define gene boundaries and a repeat GTF file provided on the Hammell lab website (downloaded on February 19 2020 from https://labshare.cshl.edu/shares/mhammelllab/www-data/TEtranscripts/TE_GTF/GRCh38_GENCODE_rmsk_TE.gtf.gz) to define repeat boundaries.

Similarly, the TElocal package (https://github.com/mhammell-laboratory/TElocal), from the same software suite as TEtranscripts, was employed to obtain gene and TE locus-specific counts using the same GENCODE annotations and a repeat file provided on the Hammell lab website (downloaded on October 31 2023 from https://labshare.cshl.edu/shares/mhammelllab/www-data/TElocal/prebuilt_indices/).

### Gene *cis*-eQTL and L1 t*rans*-eQTL analyses

Gene and TE count files were loaded into R v4.2.1. Lowly expressed genes were first filtered out if 323/358 European samples and 78/86 Yoruban samples did not have over 0.44 counts per million (cpm) or 0.43 cpm, respectively. These fractions were selected because they corresponded to expression in ∼90% of samples and thus helped maintain maximal statistical power by focusing on genes ubiquitously expressed across each entire population. The cpm thresholds were selected because they corresponded to 10 reads in the median-length library within each set of samples. For the locus-specific quantifications, repeat counts were loaded into R and stratified into the following categories: (i) ‘distal intergenic’ TEs that were >5 kb from a gene, (ii) ‘nearby intergenic’ TEs that were within 5 kb of a gene, (iii) ‘exonic’ TEs that overlapped any annotated exon, and (iv) ‘intronic’ TEs for TEs that were in a gene but did not overlap an annotated exon. The stratification was carried out in order to separately characterize the influences and responses of each TE type to our analytical groups. After stratifying, repeat counts were aggregated at the subfamily level in order to compare results with the unstratified TEtranscripts results. After aggregating, lowly expressed genes were filtered as specified above.

Then, counts underwent a variance stabilizing transformation (vst) using DESeq2 v1.36.0 (Love et al. 2014). The following covariates were regressed out from vst normalized expression data using the ‘removeBatchEffect’ function in Limma v3.52.2 (Ritchie et al. 2015): lab, population category, principal components 1-2 of the pruned SNVs, biological sex, net L1/Alu copy number, and EBV expression levels. Since the Yoruban samples were all from the same population, the population variable was omitted in their batch correction. Here, we note several things. First, EBV expression was included as a covariate because heightened TE expression is often a feature of viral infections (Macchietto et al. 2020). Secondly, although PEER (Stegle et al. 2012) is often used to remove technical variation for *cis*-eQTL analysis, this can come at the expense of correcting out genome-wide biological effects. This can be problematic in some settings, such as *trans*-eQTL analysis. Thus, PEER factors were not included. The batch-corrected data underwent a final inverse normal transformation (INT), using the RankNorm function in the R package RNOmni v1.0.1, to obtain normally distributed gene expression values.

The INT expression matrices were split into genes and L1 subfamilies, which were used to identify gene *cis*-eQTLs and L1 subfamily *trans*-eQTLs in the European superpopulation using MatrixEQTL v2.3 (Shabalin 2012). For gene *cis*-eQTLs, SNVs were tested for association with expressed genes within 1 million base pairs. We opted to use a *trans*-eQTL approach using aggregate subfamily-level TE expression since the *trans* approach should allow us to identify regulators of many elements rather than one. The Benjamini-Hochberg false discovery rate (FDR) was calculated in each analysis, and we used the p-value corresponding to an FDR of < 5% as the threshold for eQTL significance. In addition, the *cis*-eQTL and *trans*-eQTL analyses were also repeated using 20 permuted expression datasets in which the sample names were scrambled, and the p-value corresponding to an average empirical FDR of < 5% was used as a secondary threshold. To note, we calculated the average empirical FDR at a given p-value p_i_ by (i) counting the total number of null points with p ≤ p_i_, (ii) dividing by the number of permutations, to obtain an average number of null points with p ≤ p_i_, and (iii) dividing the average number of null points with p ≤ p_i_ by the number of real points with p ≤ p_i_. eQTLs were called as significant if they passed the stricter of the two thresholds. SNV-gene and SNV-L1 associations that were significant in the European superpopulation were then targeted and tested in the Yoruban population using R’s built-in linear modelling functions. In this case, only the Benjamini-Hochberg FDR was calculated, and significant eQTLs were called if they possessed an FDR < 0.05.

### Defining SNV-gene-L1 trios and mediation analysis

For each population, the *cis-* and *trans*-eQTL results were integrated to identify SNVs associated with both gene and L1 subfamily expression. We reasoned that L1 expression would respond to differences in expression of *bona fide* regulators. Consequently, gene expression and L1 subfamily expression associations were assessed by linear regression, and the p-values from this analysis were Benjamini-Hochberg FDR-corrected. Candidate SNV-gene-L1 trios were defined as those with *cis*- eQTL, *trans*-eQTL, and expression regression FDRs < 0.05. To identify top, index SNVs in regions of linkage disequilibrium (LD), SNVs within 500 kilobases of each other with an R^2^ > 0.10 were clumped together by *trans*-eQTL p-value using PLINK v1.90b6.17. Mediation analysis was carried out using the ‘gmap.gpd’ function in eQTLMAPT v0.1.0 (Wang et al. 2020) on all candidate SNV-gene-L1 trios. Empirical p-values were calculated using 30,000 permutations, and Benjamini-Hochberg FDR values were calculated from empirical p-values. Mediation effects were considered significant for trios with FDR < 0.05.

### Differential expression analysis across *trans*-eQTL SNV genotypes

Transcriptomic changes associated with alternating the allele of each SNV of interest were evaluated using DESeq2 v1.36.0. Using the same filtered counts prepared for the eQTL analysis, a linear model was constructed with the following covariates for each SNV: SNV genotype in 0/1/2 format, biological sex, lab, population category, principal components 1-2 of the pruned SNVs, and principal components 1-3 of the pruned SVs (to account for structural variant population structure). As before, the population label was omitted from the Yoruban population analysis. Significant genes and TEs were those with an FDR < 0.05.

### Functional enrichment analyses

We used the Gene Set Enrichment Analysis (GSEA) paradigm as implemented in the R package clusterProfiler v4.4.4 (Wu et al. 2021). Gene Ontology, Reactome, and Hallmark pathway gene sets were obtained from the R package msigdbr v7.5.1, an Ensembl ID-mapped collection of gene sets from the Molecular Signature Database (Subramanian et al. 2005; Liberzon et al. 2015). Additionally, TE subfamilies were aggregated into TE family gene sets using the TE family designations specified in the TE GTF file (downloaded on February 19 2020 from https://labshare.cshl.edu/shares/mhammelllab/www-data/TEtranscripts/TE_GTF/GRCh38_GENCODE_rmsk_TE.gtf.gz) used during the RNA-seq quantification step. The DESeq2 v1.36.0 Wald-statistic was used to generate a combined ranked list of genes and TEs for functional enrichment analysis. All gene sets with an FDR < 0.05 were considered significant. For plots with a single analysis, the top 5 downregulated and top 5 upregulated gene sets were plotted, at most. For plots with multiple analyses, shared gene sets with the desired expression patterns in each individual analysis were first identified. Then, the p-values for shared gene sets were combined using Fisher’s method, and this meta-analysis p-value was used to rank shared gene sets. Finally, the top 5 gene sets with one expression pattern and the top 5 gene sets with the opposite expression pattern were plotted. If there were less than 5 gene sets in either group, those were replaced with gene sets exhibiting the opposite regulation, in order to plot 10 shared gene sets whenever possible.

### Cell lines and cell culture conditions

GM12878 (RRID: CVCL_7526) lymphoblastoid cells were purchased from the Coriell Institute. We opted to use GM12878 as a well-characterized representative cell line for candidate validation, given that (i) it is of the same cell type as the transcriptomic data used here for our eQTL analysis, and (ii) its epigenomic landscape and culture conditions are well-characterized as part of the ENCODE project (The 2011; The 2012).

GM12878 cells were maintained in RPMI (Corning cat. 15-040-CV) containing 15% FBS and 1X Penicillin-Streptomycin-Glutamine (Corning cat. 30-009-CI). Cells were cultured in a humidified incubator at 37°C and 5% CO_2_, subculturing cells 1:5 once cells reached a density of ∼10^6^ mL^-1^. All cells used were maintained below passage 30 and routinely tested for mycoplasma contamination using the PlasmoTest Mycoplasma Detection Kit (InvivoGen).

### Plasmids

The empty pcDNA3.1(+) backbone (Invitrogen cat. V79020) was a kind gift from the lab of Dr. Changhan David Lee at the University of Southern California Leonard Davis School of Gerontology. Overexpression vectors for *IL16* (CloneID OHu48263C), *STARD5*-FLAG (CloneID OHu07617D), *HSD17B12*-FLAG (CloneID OHu29918D), and *RNF5*-FLAG (CloneID OHu14875D) on a pcDNA3.1 backbone were purchased from GenScript. Plasmid sequences were verified for accuracy using Plasmidsaurus’s whole plasmid sequencing service.

### Transfections

*Escherichia coli* were cultured in LB Broth (ThermoFischer Scientific) supplemented with 50 μg/mL carbenicillin to an optical density 600 (OD_600_) of 2 – 4. Plasmid extractions were carried out using the Nucleobond Xtra Midi Plus EF kit (Macherey-Nagel) following manufacturer recommendations. Plasmids were aliquoted and stored at -20°C until the time of transfection. On the day of transfection, GM12878 cells were collected in conical tubes, spun down (100xG, 5 minutes, room temperature), resuspended in fresh media, and counted by trypan blue staining using a Countess II FL automated cell counter (Thermo Fisher). The number of cells necessary for the experiment were then aliquoted, spun down, and washed with Dulbecco’s phosphate-buffered saline (DPBS)(Corning, cat. #21-031-CV).

GM12878 cells were transfected by electroporation using the Neon Transfection System (Invitrogen) with the following parameters: 1200 V, 20 ms, and 3 pulses for GM12878 cells in Buffer R. Per reaction, we maintained a plasmid mass:cell number ratio of 10 μg : 2*10^6^ cells. For mRNA-sequencing, 8*10^6^ GM12878 cells were independently transfected for each biological replicate, with 4 replicates per overexpression condition, and cultured in a T25 flask. Immediately after transfection, cells were cultured in Penicillin-Streptomycin-free media for ∼24 hours.

Afterwards, to promote selection of viable and healthy transfected GM12878 cells, we enriched for viable cells using the EasySep Dead Cell Removal (Annexin V) Kit (STEMCELL Technologies) before seeding 2*10^6^ live cells in the same media used for cell maintenance. After another 24 hours, cell viability was measured by trypan blue staining on a Countess automated cell counter and cells were spun down (100xG, 5 min, room temperature) and lysed in TRIzol Reagent (Invitrogen) for downstream total RNA isolation (see below).

### Recombinant human IL16 (rhIL16) peptide treatment

Human rIL16 was obtained from PeproTech (cat. #200-16) and resuspended in 0.1% bovine serum albumin (BSA) solution (Akron, cat. #AK8917-0100). GM12878 cells were seeded at a concentration of 500,000 live cells per mL of media on 6-well suspension plates with 3 independent replicates per condition. Cells were exposed to 0, 24, or 48 hours of 100 ng mL^-1^ of rhIL16. To replace or exchange media 24 hours after seeding, cells were transferred to conical tubes, spun down (100xG, 5 min, room temperature), resuspended in 5 mL of the appropriate media, and transferred back to 6-well suspension plates. After 48 hours, cell viability was measured by trypan blue staining and cells were spun down (100xG, 5 min, room temperature) and lysed in TRIzol Reagent (Invitrogen).

### RNA extractions and mRNA sequencing

RNA was extracted using the Direct-zol RNA Miniprep kit (Zymo Research) following manufacturer recommendations. The integrity of RNA samples was evaluated using an Agilent High Sensitivity RNA ScreenTape assay (Agilent Technologies), ensuring that all samples had a minimum eRIN score of 8 before downstream processing. We then submitted total RNA samples to Novogene (Sacramento, California) for mRNA library preparation and sequencing on the NovaSeq 6000 platform as paired-end 150 bp reads.

### Analysis of overexpression and rhIL16 exposure mRNA-seq

mRNA-seq reads were trimmed, mapped, and quantified like for the eQTL analysis, except for the overexpression sample data. For this data, one modification was made: the EBV-inclusive reference genome was further modified to include the pcDNA3.1 sequence as an additional contig. Lowly expressed genes were filtered using a cpm threshold as in the eQTL processing, but that cpm threshold had to be satisfied by as many samples as the size of the smallest biological group. For the overexpression data, surrogate variables were estimated with the ‘svaseq’ function (Leek 2014) in the R package ‘sva’ v3.44.9, and they were regressed out from the raw read counts using the ‘removeBatchEffect’ function in the R package Limma v3.52.2. DESeq2 was used to identify significantly (FDR < 0.05) differentially expressed genes and TEs between groups. Functional enrichment analysis was carried out as previously described.

### PheWAS analysis

To gather the known associated traits for the 499 TE-related SNVs, we used Open Targets Genetics (https://genetics.opentargets.org/), a database of GWAS summary statistics (Ghoussaini et al. 2021). First, we queried the database using the 499 TE-related SNVs and collected traits that were directly associated (with P < 5x10^-8^) with the SNVs, as well as traits associated with lead variants that were in linkage disequilibrium (LD) with the queried SNPs (with R^2^ > 0.6). For age-related traits (ARTs), we used the comprehensive list of 365 Medical Subject Headings (MeSH) terms reported by (Kim et al. 2021) (downloaded from https://github.com/kisudsoe/Age-related-traits). To identify known age-related traits, the known associated traits were translated into the equivalent MeSH terms using the method described by (Kim et al. 2021). Then, the MeSH-translated known associated traits for the 499 TE-related SNVs were filtered by the MeSH terms for age-related traits.

As a parallel approach, we mapped the RsIDs for all SNVs used during the eQTL analyses to their corresponding bi-allelic Open Targets variant IDs, when available. The variant IDs corresponding to L1 *trans*-eQTL SNVs were extracted, and 500 different equal-length combinations of random SNVs were generated. Next, we queried the Open Targets database using the lists of L1-associated and random SNVs and collected the associated traits (with P < 5x10^-8^). Importantly, the database assigns traits to broader categories, including 14 disease categories that we considered age-related. We counted the number of L1-associated or random SNVs mapping to each category, and we used the random SNV counts to generate an empirical cumulative distribution function (ecdf) for each category. We calculated enrichment p-values using the formula p = 1- ecdf(mapped eQTLs) and then Benjamini-Hochberg FDR-corrected all p-values. An enrichment score (ES) was also calculated for each category using the formula ES = number of mapped L1 eQTLs / median number of randomly mapping SNVs. Categories with an ES > 1 and FDR < 0.05 were considered significantly enriched.

### Mouse husbandry

All animals were treated and housed in accordance with the Guide for Care and Use of Laboratory Animals. All experimental procedures were approved by the University of Southern California’s Institutional Animal Care and Use Committee (IACUC) and are in accordance with institutional and national guidelines. Samples were derived from animals on approved IACUC protocol #20770.

### Quantification of mouse serum IL16 by ELISA

Serum was collected from male and female C57BL/6JNia mice (4-6 and 20-24 months old) obtained from the National Institute on Aging (NIA) colony at Charles Rivers. All animals were euthanized between 8-11 am in a “snaking order” across all groups to minimize batch-processing confounds due to circadian processes. All animals were euthanized by CO_2_ asphyxiation followed by cervical dislocation. Circulating IL16 levels were quantitatively evaluated from mouse serum by enzyme-linked immunosorbent assay (ELISA). Serum was diluted 1/10 before quantifying IL16 concentrations using Abcam’s Mouse IL-16 ELISA Kit (ab201282) in accordance with manufacturer instructions. Technical replicates from the same sample were averaged to one value before statistical analysis and plotting. P-values across age within each sex were calculated using a non-parametric 2-sided Wilcoxon test, and p-values from each sex-specific analysis were combined using Fisher’s method.

## Data availability

New sequencing data generated in this study is accessible through the Sequence Read Archive (SRA) under BioProject PRJNA937306. All code is available on the Benayoun lab GitHub (https://github.com/BenayounLaboratory/TE-eQTL_LCLs). Analyses were conducted using R version 4.2.1. Code was re-run independently on R version 4.3.0 to check for reproducibility.

## Acknowledgements

Some panels were created with BioRender.com. We would like to thank Prof. Rachel Brem for her feedback and insights on the eQTL analyses. We would also like to thank Dr. Minhoo Kim for her feedback on the manuscript.

This work was supported by NSF Graduate Research Fellowship Program (NSF GRFP) DGE-1842487 (J.I.B.), NIA T32 AG052374 (J.I.B.), the University of Southern California with a Provost Fellowship (J.I.B.), NIA R25 AG076400 (C.R.M.), and NIGMS R35 GM142395 (to B.A.B).

## Author contributions

J.I.B. and B.A.B designed the study. J.I.B., L.Z., and S.K. performed data analyses, with guidance from Y.S. and B.A.B. J.I.B. and C.R.M. carried out experiments. J.I.B., B.A.B., S.K., and Y.S. wrote the manuscript. All authors contributed to the editing of the manuscript.

## References

Ahl V, Keller H, Schmidt S, Weichenrieder O. 2015. Retrotransposition and Crystal Structure of an Alu RNP in the Ribosome-Stalling Conformation. Molecular Cell 60: 715–727.

Ardeljan D, Steranka JP, Liu C, Li Z, Taylor MS, Taylor MS, Payer LM, Gorbounov M, Sarnecki JS, Deshpande V et al. 2020. Cell fitness screens reveal a conflict between LINE-1 retrotransposition and DNA replication. Nat Struct Mol Biol 27: 168–178.

Auton A Abecasis GR Altshuler DM Durbin RM Abecasis GR Bentley DR Chakravarti A Clark AG Donnelly P Eichler EE et al. 2015. A global reference for human genetic variation. Nature 526: 68–74.

Baeken MW, Moosmann B, Hajieva P. 2020. Retrotransposon activation by distressed mitochondria in neurons. Biochemical and Biophysical Research Communications 525: 570–575.

Baier M, Werner A, Bannert N, Metzner K, Kurth R. 1995. HIV suppression by interleukin-16. Nature 378: 563.

Bravo JI, Nozownik S, Danthi PS, Benayoun BA. 2020. Transposable elements, circular RNAs and mitochondrial transcription in age-related genomic regulation. Development 147.

Briggs EM, Mita P, Sun X, Ha S, Vasilyev N, Leopold ZR, Nudler E, Boeke JD, Logan SK. 2021. Unbiased proteomic mapping of the LINE-1 promoter using CRISPR Cas9. Mobile DNA 12: 21.

Brouha B, Schustak J, Badge RM, Lutz-Prigge S, Farley AH, Moran JV, Kazazian HH. 2003. Hot L1s account for the bulk of retrotransposition in the human population. Proceedings of the National Academy of Sciences 100: 5280–5285.

Campisi J. 2013. Aging, Cellular Senescence, and Cancer. Annual Review of Physiology 75: 685–705.

Carithers LJ, Ardlie K, Barcus M, Branton PA, Britton A, Buia SA, Compton CC, DeLuca DS, Peter-Demchok J, Gelfand ET et al. 2015. A Novel Approach to High-Quality Postmortem Tissue Procurement: The GTEx Project. Biopreservation and Biobanking 13: 311–319.

Center DM, Cruikshank W. 1982. Modulation of lymphocyte migration by human lymphokines. I. Identification and characterization of chemoattractant activity for lymphocytes from mitogen-stimulated mononuclear cells. J Immunol 128: 2563–2568.

Center DM, Kornfeld H, Cruikshank WW. 1996. Interleukin 16 and its function as a CD4 ligand. Immunol Today 17: 476–481.

Chen S, Zhou Y, Chen Y, Gu J. 2018. fastp: an ultra-fast all-in-one FASTQ preprocessor. Bioinformatics 34: i884–i890.

Chick JM, Munger SC, Simecek P, Huttlin EL, Choi K, Gatti DM, Raghupathy N, Svenson KL, Churchill GA, Gygi SP. 2016. Defining the consequences of genetic variation on a proteome-wide scale. Nature 534: 500–505.

Chung N, Jonaid GM, Quinton S, Ross A, Sexton CE, Alberto A, Clymer C, Churchill D, Navarro Leija O, Han MV. 2019. Transcriptome analyses of tumor-adjacent somatic tissues reveal genes co-expressed with transposable elements. Mobile DNA 10: 39.

Cruikshank WW, Center DM, Nisar N, Wu M, Natke B, Theodore AC, Kornfeld H. 1994. Molecular and functional analysis of a lymphocyte chemoattractant factor: association of biologic function with CD4 expression. Proc Natl Acad Sci U S A 91: 5109–5113.

Cruikshank WW, Lim K, Theodore AC, Cook J, Fine G, Weller PF, Center DM. 1996. IL-16 inhibition of CD3-dependent lymphocyte activation and proliferation. J Immunol 157: 5240–5248.

Danecek P, Auton A, Abecasis G, Albers CA, Banks E, DePristo MA, Handsaker RE, Lunter G, Marth GT, Sherry ST et al. 2011. The variant call format and VCFtools. Bioinformatics 27: 2156–2158.

Danecek P, Bonfield JK, Liddle J, Marshall J, Ohan V, Pollard MO, Whitwham A, Keane T, McCarthy SA, Davies RM et al. 2021. Twelve years of SAMtools and BCFtools. GigaScience 10.

De Cecco M, Criscione SW, Peckham EJ, Hillenmeyer S, Hamm EA, Manivannan J, Peterson AL, Kreiling JA, Neretti N, Sedivy JM. 2013a. Genomes of replicatively senescent cells undergo global epigenetic changes leading to gene silencing and activation of transposable elements. Aging Cell 12: 247–256.

De Cecco M, Criscione SW, Peterson AL, Neretti N, Sedivy JM, Kreiling JA. 2013b. Transposable elements become active and mobile in the genomes of aging mammalian somatic tissues. Aging (Albany NY) 5: 867–883.

De Cecco M, Ito T, Petrashen AP, Elias AE, Skvir NJ, Criscione SW, Caligiana A, Brocculi G, Adney EM, Boeke JD et al. 2019. L1 drives IFN in senescent cells and promotes age-associated inflammation. Nature 566: 73–78.

de Tribolet-Hardy J, Thorball CW, Forey R, Planet E, Duc J, Coudray A, Khubieh B, Offner S, Pulver C, Fellay J et al. 2023. Genetic features and genomic targets of human KRAB-zinc finger proteins. Genome Research 33: 1409–1423.

Della Valle F, Reddy P, Yamamoto M, Liu P, Saera-Vila A, Bensaddek D, Zhang H, Prieto Martinez J, Abassi L, Celii M et al. 2022. *LINE-1* RNA causes heterochromatin erosion and is a target for amelioration of senescent phenotypes in progeroid syndromes. Science Translational Medicine 14: eabl6057.

Didier C, Broday L, Bhoumik A, Israeli S, Takahashi S, Nakayama K, Thomas SM, Turner CE, Henderson S, Sabe H et al. 2003. RNF5, a RING Finger Protein That Regulates Cell Motility by Targeting Paxillin Ubiquitination and Altered Localization. Molecular and Cellular Biology 23: 5331–5345.

Dobin A, Davis CA, Schlesinger F, Drenkow J, Zaleski C, Jha S, Batut P, Chaisson M, Gingeras TR. 2012. STAR: ultrafast universal RNA-seq aligner. Bioinformatics 29: 15–21.

Feng K, Min Y-Q, Sun X, Deng F, Li P, Wang H, Ning Y-J. 2021. Interactome profiling reveals interaction of SARS-CoV-2 NSP13 with host factor STAT1 to suppress interferon signaling. Journal of Molecular Cell Biology 13: 760–762.

Flasch DA, Chen X, Ju B, Li X, Dalton J, Mulder HL, Easton J, Wang L, Baker SJ, Chiang J et al. 2022. Somatic LINE-1 promoter acquisition drives oncogenic FOXR2 activation in pediatric brain tumor. Acta Neuropathologica 143: 605–607.

Franceschi C, Garagnani P, Parini P, Giuliani C, Santoro A. 2018. Inflammaging: a new immune–metabolic viewpoint for age-related diseases. Nature Reviews Endocrinology 14: 576–590.

Frankish A, Diekhans M, Ferreira A-M, Johnson R, Jungreis I, Loveland J, Mudge JM, Sisu C, Wright J, Armstrong J et al. 2018. GENCODE reference annotation for the human and mouse genomes. Nucleic Acids Research 47: D766–D773.

Gao F, Chang D, Biddanda A, Ma L, Guo Y, Zhou Z, Keinan A. 2015. XWAS: A Software Toolset for Genetic Data Analysis and Association Studies of the X Chromosome. J Hered 106: 666–671.

Germain MA, Chatel-Chaix L, Gagné B, Bonneil É, Thibault P, Pradezynski F, de Chassey B, Meyniel-Schicklin L, Lotteau V, Baril M et al. 2014. Elucidating novel hepatitis C virus-host interactions using combined mass spectrometry and functional genomics approaches. Mol Cell Proteomics 13: 184–203.

Ghoussaini M, Mountjoy E, Carmona M, Peat G, Schmidt EM, Hercules A, Fumis L, Miranda A, Carvalho-Silva D, Buniello A et al. 2021. Open Targets Genetics: systematic identification of trait-associated genes using large-scale genetics and functional genomics. Nucleic Acids Res 49: D1311–d1320.

Goubert C, Zevallos NA, Feschotte C. 2020. Contribution of unfixed transposable element insertions to human regulatory variation. Philosophical Transactions of the Royal Society B: Biological Sciences 375: 20190331.

Gu Z, Liu Y, Zhang Y, Cao H, Lyu J, Wang X, Wylie A, Newkirk SJ, Jones AE, Lee M et al. 2021. Silencing of LINE-1 retrotransposons is a selective dependency of myeloid leukemia. Nature Genetics 53: 672–682.

Heikelä H, Ruohonen ST, Adam M, Viitanen R, Liljenbäck H, Eskola O, Gabriel M, Mairinoja L, Pessia A, Velagapudi V et al. 2020. Hydroxysteroid (17β) dehydrogenase 12 is essential for metabolic homeostasis in adult mice. Am J Physiol Endocrinol Metab 319: E494–e508.

Helleboid P-Y, Heusel M, Duc J, Piot C, Thorball CW, Coluccio A, Pontis J, Imbeault M, Turelli P, Aebersold R et al. 2019. The interactome of KRAB zinc finger proteins reveals the evolutionary history of their functional diversification. The EMBO Journal 38: e101220.

Huang Y, Du KL, Guo PY, Zhao RM, Wang B, Zhao XL, Zhang CQ. 2019. IL-16 regulates macrophage polarization as a target gene of mir-145-3p. Mol Immunol 107: 1–9.

Hussain T, Mulherkar R. 2012. Lymphoblastoid Cell lines: a Continuous in Vitro Source of Cells to Study Carcinogen Sensitivity and DNA Repair. Int J Mol Cell Med 1: 75–87.

Idziorek T, Khalife J, Billaut-Mulot O, Hermann E, Aumercier M, Mouton Y, Capron A, Bahr GM. 1998. Recombinant human IL-16 inhibits HIV-1 replication and protects against activation-induced cell death (AICD). Clin Exp Immunol 112: 84–91.

Jeck WR, Siebold AP, Sharpless NE. 2012. Review: a meta-analysis of GWAS and age-associated diseases. Aging Cell 11: 727–731.

Jia R, Jiang C, Li L, Huang C, Lu L, Xu M, Xu J, Liang X. 2021. Interleukin 16 Enhances the Host Susceptibility to Influenza A Virus Infection. Frontiers in Microbiology 12.

Jin Y, Tam OH, Paniagua E, Hammell M. 2015. TEtranscripts: a package for including transposable elements in differential expression analysis of RNA-seq datasets. Bioinformatics 31: 3593–3599.

Keur N, Ricaño-Ponce I, Kumar V, Matzaraki V. 2022. A systematic review of analytical methods used in genetic association analysis of the X-chromosome. Briefings in Bioinformatics 23.

Kim S-S, Hudgins AD, Gonzalez B, Milman S, Barzilai N, Vijg J, Tu Z, Suh Y. 2021. A Compendium of Age-Related PheWAS and GWAS Traits for Human Genetic Association Studies, Their Networks and Genetic Correlations. Frontiers in Genetics 12.

Lai RW, Lu R, Danthi PS, Bravo JI, Goumba A, Sampathkumar NK, Benayoun BA. 2019. Multi-level remodeling of transcriptional landscapes in aging and longevity. BMB Rep 52: 86–108.

Lander ES Linton LM Birren B Nusbaum C Zody MC Baldwin J Devon K Dewar K Doyle M FitzHugh W et al. 2001. Initial sequencing and analysis of the human genome. Nature 409: 860-921.

Lappalainen T, Sammeth M, Friedländer MR, ‘t Hoen PAC, Monlong J, Rivas MA, Gonzàlez-Porta M, Kurbatova N, Griebel T, Ferreira PG, et al. 2013. Transcriptome and genome sequencing uncovers functional variation in humans. Nature 501: 506–511.

Leek JT. 2014. svaseq: removing batch effects and other unwanted noise from sequencing data. Nucleic Acids Research 42: e161–e161.

Levin HL, Moran JV. 2011. Dynamic interactions between transposable elements and their hosts. Nature Reviews Genetics 12: 615–627.

Li Z, Hao P, Zhao Z, Gao W, Huan C, Li L, Chen X, Wang H, Jin N, Luo Z-Q et al. 2023. The E3 ligase RNF5 restricts SARS-CoV-2 replication by targeting its envelope protein for degradation. Signal Transduction and Targeted Therapy 8: 53.

Liao KC, Garcia-Blanco MA. 2021. Role of Alternative Splicing in Regulating Host Response to Viral Infection. Cells 10.

Liberzon A, Birger C, Thorvaldsdóttir H, Ghandi M, Mesirov JP, Tamayo P. 2015. The Molecular Signatures Database (MSigDB) hallmark gene set collection. Cell Syst 1: 417–425.

Lindholm HT, Chen R, De Carvalho DD. 2023. Endogenous retroelements as alarms for disruptions to cellular homeostasis. Trends in Cancer 9: 55–68.

Liu EY, Russ J, Cali CP, Phan JM, Amlie-Wolf A, Lee EB. 2019. Loss of Nuclear TDP-43 Is Associated with Decondensation of LINE Retrotransposons. Cell Reports 27: 1409–1421.e1406.

Liu N, Lee CH, Swigut T, Grow E, Gu B, Bassik MC, Wysocka J. 2018. Selective silencing of euchromatic L1s revealed by genome-wide screens for L1 regulators. Nature 553: 228–232.

Liu S, Lei Z, Li J, Wang L, Jia R, Liu Z, Jiang C, Gao Y, Liu M, Kuang L et al. 2020. Interleukin 16 contributes to gammaherpesvirus pathogenesis by inhibiting viral reactivation. PLoS Pathog 16: e1008701.

Lonsdale J Thomas J Salvatore M Phillips R Lo E Shad S Hasz R Walters G Garcia F Young N et al. 2013. The Genotype-Tissue Expression (GTEx) project. Nature Genetics 45: 580–585.

Love MI, Huber W, Anders S. 2014. Moderated estimation of fold change and dispersion for RNA-seq data with DESeq2. Genome Biology 15: 550.

Luqman-Fatah A, Watanabe Y, Uno K, Ishikawa F, Moran JV, Miyoshi T. 2023. The interferon stimulated gene-encoded protein HELZ2 inhibits human LINE-1 retrotransposition and LINE-1 RNA-mediated type I interferon induction. Nature Communications 14: 203.

Luu-The V, Tremblay P, Labrie F. 2006. Characterization of type 12 17beta-hydroxysteroid dehydrogenase, an isoform of type 3 17beta-hydroxysteroid dehydrogenase responsible for estradiol formation in women. Mol Endocrinol 20: 437–443.

Macchietto MG, Langlois RA, Shen SS. 2020. Virus-induced transposable element expression up-regulation in human and mouse host cells. Life Sci Alliance 3.

Marasca F, Sinha S, Vadalà R, Polimeni B, Ranzani V, Paraboschi EM, Burattin FV, Ghilotti M, Crosti M, Negri ML et al. 2022. LINE1 are spliced in non-canonical transcript variants to regulate T cell quiescence and exhaustion. Nature Genetics 54: 180–193.

Mita P, Sun X, Fenyö D, Kahler DJ, Li D, Agmon N, Wudzinska A, Keegan S, Bader JS, Yun C et al. 2020. BRCA1 and S phase DNA repair pathways restrict LINE-1 retrotransposition in human cells. Nat Struct Mol Biol 27: 179–191.

Mohamed B, Mazeaud C, Baril M, Poirier D, Sow AA, Chatel-Chaix L, Titorenko V, Lamarre D. 2020. Very-long-chain fatty acid metabolic capacity of 17-beta-hydroxysteroid dehydrogenase type 12 (HSD17B12) promotes replication of hepatitis C virus and related flaviviruses. Scientific Reports 10: 4040.

Moran JV, Holmes SE, Naas TP, DeBerardinis RJ, Boeke JD, Kazazian HH, Jr. 1996. High Frequency Retrotransposition in Cultured Mammalian Cells. Cell 87: 917–927.

Nagasaki S, Miki Y, Akahira J, Suzuki T, Sasano H. 2009. Transcriptional regulation of 17beta-hydroxysteroid dehydrogenase type 12 by SREBP-1. Mol Cell Endocrinol 307: 163–168.

Pasquesi GIM, Allen H, Ivancevic A, Barbachano-Guerrero A, Joyner O, Guo K, Simpson DM, Gapin K, Horton I, Nguyen L et al. 2023. Regulation of human interferon signaling by transposon exonization. bioRxiv doi:10.1101/2023.09.11.557241.

Purcell S, Neale B, Todd-Brown K, Thomas L, Ferreira MAR, Bender D, Maller J, Sklar P, de Bakker PIW, Daly MJ et al. 2007. PLINK: A Tool Set for Whole-Genome Association and Population-Based Linkage Analyses. The American Journal of Human Genetics 81: 559–575.

Ramirez J, Bitterman P, Basu S, Barua A. 2022. Changes in IL-16 Expression in the Ovary during Aging and Its Potential Consequences to Ovarian Pathology. J Immunol Res 2022: 2870389.

Rebollo R, Romanish MT, Mager DL. 2012. Transposable Elements: An Abundant and Natural Source of Regulatory Sequences for Host Genes. Annual Review of Genetics 46: 21–42.

Ritchie ME, Phipson B, Wu D, Hu Y, Law CW, Shi W, Smyth GK. 2015. limma powers differential expression analyses for RNA-sequencing and microarray studies. Nucleic Acids Research 43: e47–e47.

Rodić N, Sharma R, Sharma R, Zampella J, Dai L, Taylor MS, Hruban RH, Iacobuzio-Donahue CA, Maitra A, Torbenson MS et al. 2014. Long Interspersed Element-1 Protein Expression Is a Hallmark of Many Human Cancers. The American Journal of Pathology 184: 1280–1286.

Rodriguez-Agudo D, Calderon-Dominguez M, Medina MA, Ren S, Gil G, Pandak WM. 2012. ER stress increases StarD5 expression by stabilizing its mRNA and leads to relocalization of its protein from the nucleus to the membranes. J Lipid Res 53: 2708–2715.

Rodriguez-Agudo D, Malacrida L, Kakiyama G, Sparrer T, Fortes C, Maceyka M, Subler MA, Windle JJ, Gratton E, Pandak WM et al. 2019. StarD5: an ER stress protein regulates plasma membrane and intracellular cholesterol homeostasis. Journal of Lipid Research 60: 1087–1098.

Rodriguez-Martin B Alvarez EG Baez-Ortega A Zamora J Supek F Demeulemeester J Santamarina M Ju YS Temes J Garcia-Souto D et al. 2020. Pan-cancer analysis of whole genomes identifies driver rearrangements promoted by LINE-1 retrotransposition. Nature Genetics 52: 306–319.

Sampathkumar NK, Bravo JI, Chen Y, Danthi PS, Donahue EK, Lai RW, Lu R, Randall LT, Vinson N, Benayoun BA. 2020. Widespread sex dimorphism in aging and age-related diseases. Hum Genet 139: 333–356.

Sato S, Gillette M, de Santiago PR, Kuhn E, Burgess M, Doucette K, Feng Y, Mendez-Dorantes C, Ippoliti PJ, Hobday S et al. 2023. LINE-1 ORF1p as a candidate biomarker in high grade serous ovarian carcinoma. Scientific Reports 13: 1537.

Savytska N, Heutink P, Bansal V. 2022. Transcription start site signal profiling improves transposable element RNA expression analysis at locus-level. Frontiers in Genetics 13.

Shabalin AA. 2012. Matrix eQTL: ultra fast eQTL analysis via large matrix operations. Bioinformatics 28: 1353–1358.

Sie L, Loong S, Tan EK. 2009. Utility of lymphoblastoid cell lines. Journal of Neuroscience Research 87: 1953–1959.

Simon M, Van Meter M, Ablaeva J, Ke Z, Gonzalez RS, Taguchi T, De Cecco M, Leonova KI, Kogan V, Helfand SL et al. 2019. LINE1 Derepression in Aged Wild-Type and SIRT6-Deficient Mice Drives Inflammation. Cell Metabolism 29: 871–885.e875.

Soccio RE, Adams RM, Maxwell KN, Breslow JL. 2005. Differential Gene Regulation of StarD4 and StarD5 Cholesterol Transfer Proteins: ACTIVATION OF StarD4 BY STEROL REGULATORY ELEMENT-BINDING PROTEIN-2 AND StarD5 BY ENDOPLASMIC RETICULUM STRESS *. Journal of Biological Chemistry 280: 19410–19418.

Spirito G, Mangoni D, Sanges R, Gustincich S. 2019. Impact of polymorphic transposable elements on transcription in lymphoblastoid cell lines from public data. BMC Bioinformatics 20: 495.

Stegle O, Parts L, Piipari M, Winn J, Durbin R. 2012. Using probabilistic estimation of expression residuals (PEER) to obtain increased power and interpretability of gene expression analyses. Nat Protoc 7: 500–507.

Streeter I, Harrison PW, Faulconbridge A, The HipSci Consortium, Flicek P, Parkinson H, Clarke L. 2016. The human-induced pluripotent stem cell initiative—data resources for cellular genetics. Nucleic Acids Research 45: D691–D697.

Subramanian A, Tamayo P, Mootha VK, Mukherjee S, Ebert BL, Gillette MA, Paulovich A, Pomeroy SL, Golub TR, Lander ES et al. 2005. Gene set enrichment analysis: A knowledge-based approach for interpreting genome-wide expression profiles. Proceedings of the National Academy of Sciences 102: 15545–15550.

Sudmant PH, Rausch T, Gardner EJ, Handsaker RE, Abyzov A, Huddleston J, Zhang Y, Ye K, Jun G, Hsi-Yang Fritz M et al. 2015. An integrated map of structural variation in 2,504 human genomes. Nature 526: 75–81.

Sun X, Wang X, Tang Z, Grivainis M, Kahler D, Yun C, Mita P, Fenyö D, Boeke JD. 2018. Transcription factor profiling reveals molecular choreography and key regulators of human retrotransposon expression. Proc Natl Acad Sci U S A 115: E5526–e5535.

Tcherpakov M, Delaunay A, Toth J, Kadoya T, Petroski MD, Ronai ZeA. 2009. Regulation of Endoplasmic Reticulum-associated Degradation by RNF5-dependent Ubiquitination of JNK-associated Membrane Protein (JAMP) *. Journal of Biological Chemistry 284: 12099–12109.

The EPC. 2011. A User’s Guide to the Encyclopedia of DNA Elements (ENCODE). PLOS Biology 9: e1001046.

The EPC. 2012. An integrated encyclopedia of DNA elements in the human genome. Nature 489: 57–74.

Theodore AC, Center DM, Nicoll J, Fine G, Kornfeld H, Cruikshank WW. 1996. CD4 ligand IL-16 inhibits the mixed lymphocyte reaction. J Immunol 157: 1958–1964.

Tiwari B, Jones AE, Caillet CJ, Das S, Royer SK, Abrams JM. 2020. p53 directly represses human LINE1 transposons. Genes & Development 34: 1439–1451.

Valdebenito-Maturana B, Valdebenito-Maturana F, Carrasco M, Tapia JC, Maureira A. 2023. Activation of Transposable Elements in Human Skeletal Muscle Fibers upon Statin Treatment. International Journal of Molecular Sciences 24: 244.

Venter JC Adams MD Myers EW Li PW Mural RJ Sutton GG Smith HO Yandell M Evans CA Holt RA et al. 2001. The Sequence of the Human Genome. Science 291: 1304–1351.

Wahl D, Cavalier AN, Smith M, Seals DR, LaRocca TJ. 2020. Healthy Aging Interventions Reduce Repetitive Element Transcripts. The Journals of Gerontology: Series A 76: 805–810.

Wang L, Norris ET, Jordan IK. 2017. Human Retrotransposon Insertion Polymorphisms Are Associated with Health and Disease via Gene Regulatory Phenotypes. Frontiers in Microbiology 8.

Wang L, Rishishwar L, Mariño-Ramírez L, Jordan IK. 2016. Human population-specific gene expression and transcriptional network modification with polymorphic transposable elements. Nucleic Acids Research 45: 2318–2328.

Wang T, Peng Q, Liu B, Liu X, Liu Y, Peng J, Wang Y. 2020. eQTLMAPT: Fast and Accurate eQTL Mediation Analysis With Efficient Permutation Testing Approaches. Frontiers in Genetics 10.

Watanabe T, Watanabe S, Kawaoka Y. 2010. Cellular Networks Involved in the Influenza Virus Life Cycle. Cell Host & Microbe 7: 427–439.

Williams TM. 2001. Human Leukocyte Antigen Gene Polymorphism and the Histocompatibility Laboratory. The Journal of Molecular Diagnostics 3: 98–104.

Wilson KC, Center DM, Cruikshank WW. 2004. Mini ReviewThe Effect of Interleukin-16 and its Precursor on T Lymphocyte Activation and Growth. Growth Factors 22: 97–104.

Wong C-J, Whiddon JL, Langford AT, Belleville AE, Tapscott SJ. 2021. Canine DUXC: implications for DUX4 retrotransposition and preclinical models of FSHD. Human Molecular Genetics 31: 1694–1704.

Wu T, Hu E, Xu S, Chen M, Guo P, Dai Z, Feng T, Zhou L, Tang W, Zhan L et al. 2021. clusterProfiler 4.0: A universal enrichment tool for interpreting omics data. The Innovation 2: 100141.

Yuan Z, Hu B, Xiao H, Tan X, Li Y, Tang K, Zhang Y, Cai K, Ding B. 2022. The E3 Ubiquitin Ligase RNF5 Facilitates SARS-CoV-2 Membrane Protein-Mediated Virion Release. mBio 13: e03168–03121.

Zhang M, Sun W, You X, Xu D, Wang L, Yang J, Li E, He S. 2023. LINE-1 repression in Epstein–Barr virus-associated gastric cancer through viral–host genome interaction. Nucleic Acids Research 51: 4867–4880.

Zhao H, Ji Q, Wu Z, Wang S, Ren J, Yan K, Wang Z, Hu J, Chu Q, Hu H et al. 2022. Destabilizing heterochromatin by APOE mediates senescence. Nature Aging 2: 303–316.

Zhao Y, Oreskovic E, Zhang Q, Lu Q, Gilman A, Lin YS, He J, Zheng Z, Lu JY, Lee J et al. 2021. Transposon-triggered innate immune response confers cancer resistance to the blind mole rat. Nature Immunology 22: 1219–1230.

Zhong B, Zhang L, Lei C, Li Y, Mao A-P, Yang Y, Wang Y-Y, Zhang X-L, Shu H-B. 2009. The Ubiquitin Ligase RNF5 Regulates Antiviral Responses by Mediating Degradation of the Adaptor Protein MITA. Immunity 30: 397–407.

Zhong B, Zhang Y, Tan B, Liu TT, Wang YY, Shu HB. 2010. The E3 ubiquitin ligase RNF5 targets virus-induced signaling adaptor for ubiquitination and degradation. J Immunol 184: 6249–6255.

Zhou P, Goldstein S, Devadas K, Tewari D, Notkins AL. 1997. Human CD4+ cells transfected with IL-16 cDNA are resistant to HIV-1 infection: inhibition of mRNA expression. Nat Med 3: 659–664.

