## Supplemental figures and legends for "An eQTL-based Approach Reveals Candidate Regulators of LINE-1 RNA Levels in Lymphoblastoid Cells"

**SUPPLEMENTARY MATERIAL**

**Supplementary figures**

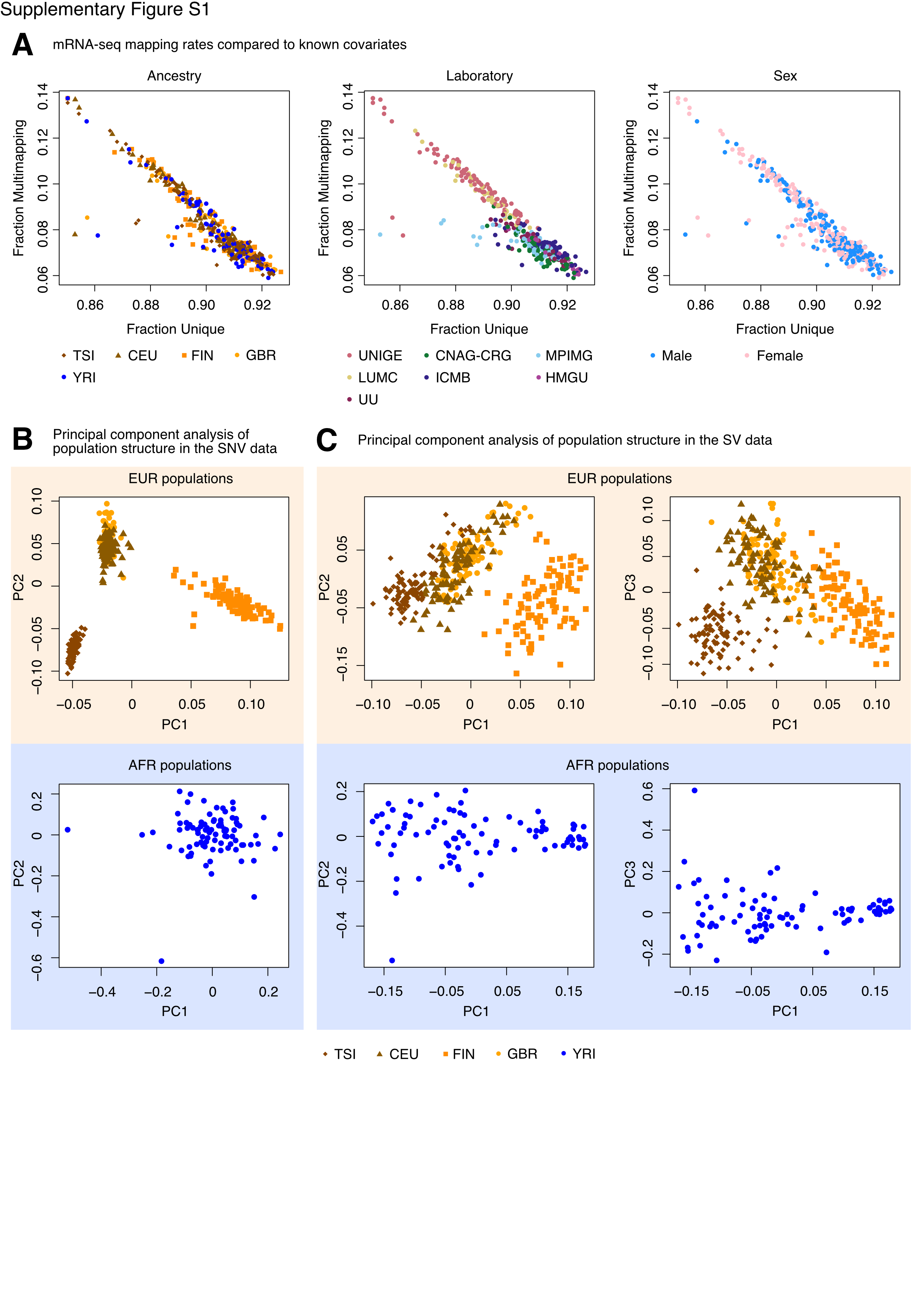

**Supplementary Figure S1. Quality control of GEUVADIS and 1000Genomes data.**

**(A)** Unique and multimapping fractions for the mRNA-sequencing data, color-coded by ancestry group, laboratory, or biological sex. For visual clarity, the plots were limited to samples with a minimum of 84% uniquely mapped reads, a limit which still captured the majority of samples analyzed. For ancestry, the following groups were analyzed: Tuscan (TSI), Northern Europeans from Utah (CEU), Finnish (FIN), British (GBR), and Yoruba (YRI). **(B)** PCA plots for pruned SNV genotype data from European or African samples, color-coded and shaped according to ancestry group. **(C)** PCA plots for pruned SV genotype data from European or African samples, color-coded and shaped according to ancestry group.

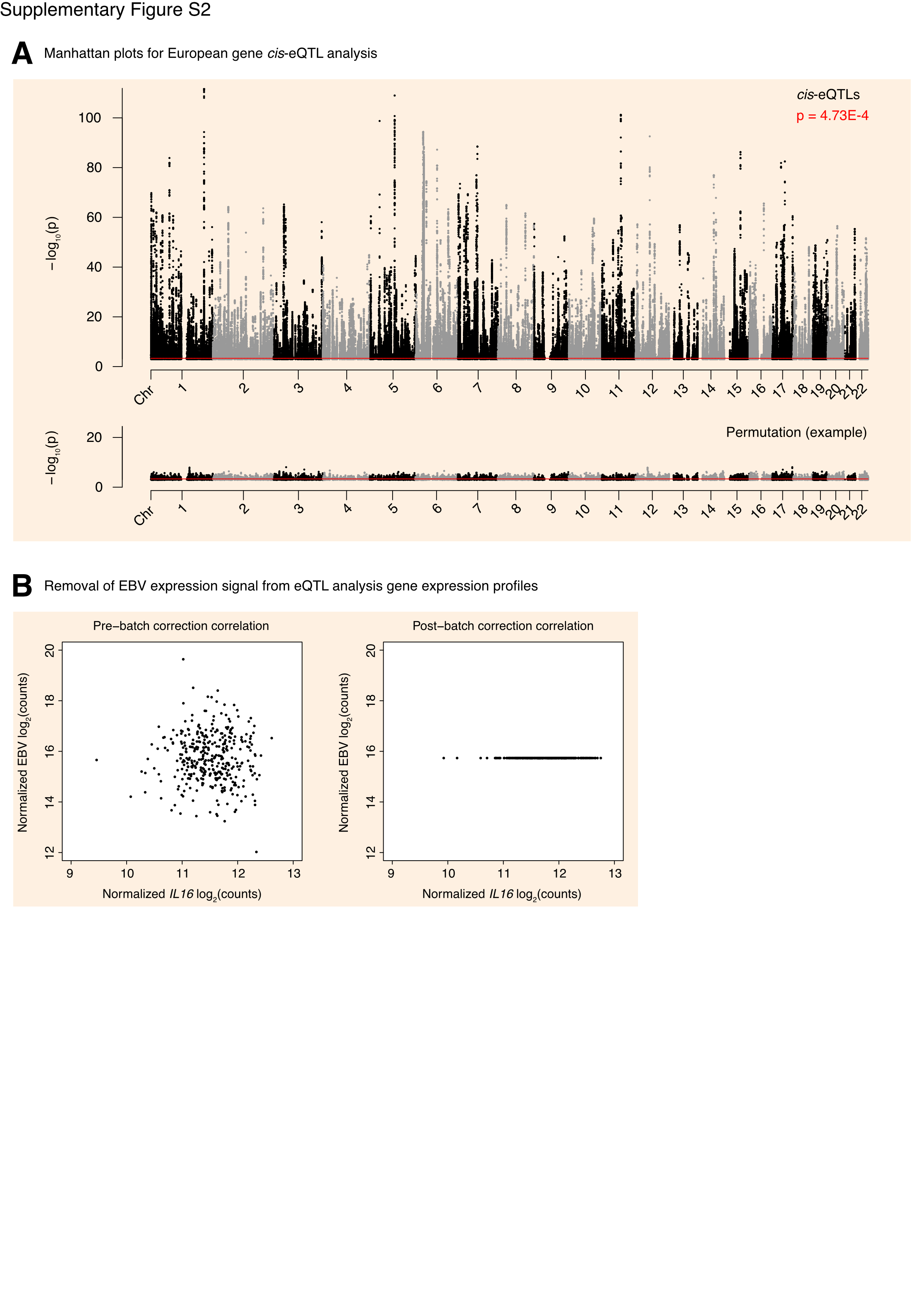

**Supplementary Figure S2.** **Annotation of genome-wide SNVs with *cis*-eQTLs in the European cohort.**

**(A)** A Manhattan plot for the gene *cis*-eQTL analysis in the European cohort. The dashed line at p = 4.75E-4 corresponds to an average empirical FDR < 0.05, based on 20 random permutations. One such permutation is illustrated in the bottom panel. The solid line at p = 4.73E-4 corresponds to a Benjamini-Hochberg FDR < 0.05. The stricter of the two thresholds, p = 4.73E-4, was used to define significant *cis*-eQTLs. **(B)** EBV expression as a function of *IL16* expression prior to (left) and after (right) correcting for EBV expression. This correction was applied prior to the eQTL scan in order to avoid confounding SNV associations with differences in EBV expression. FDR: False Discovery Rate.

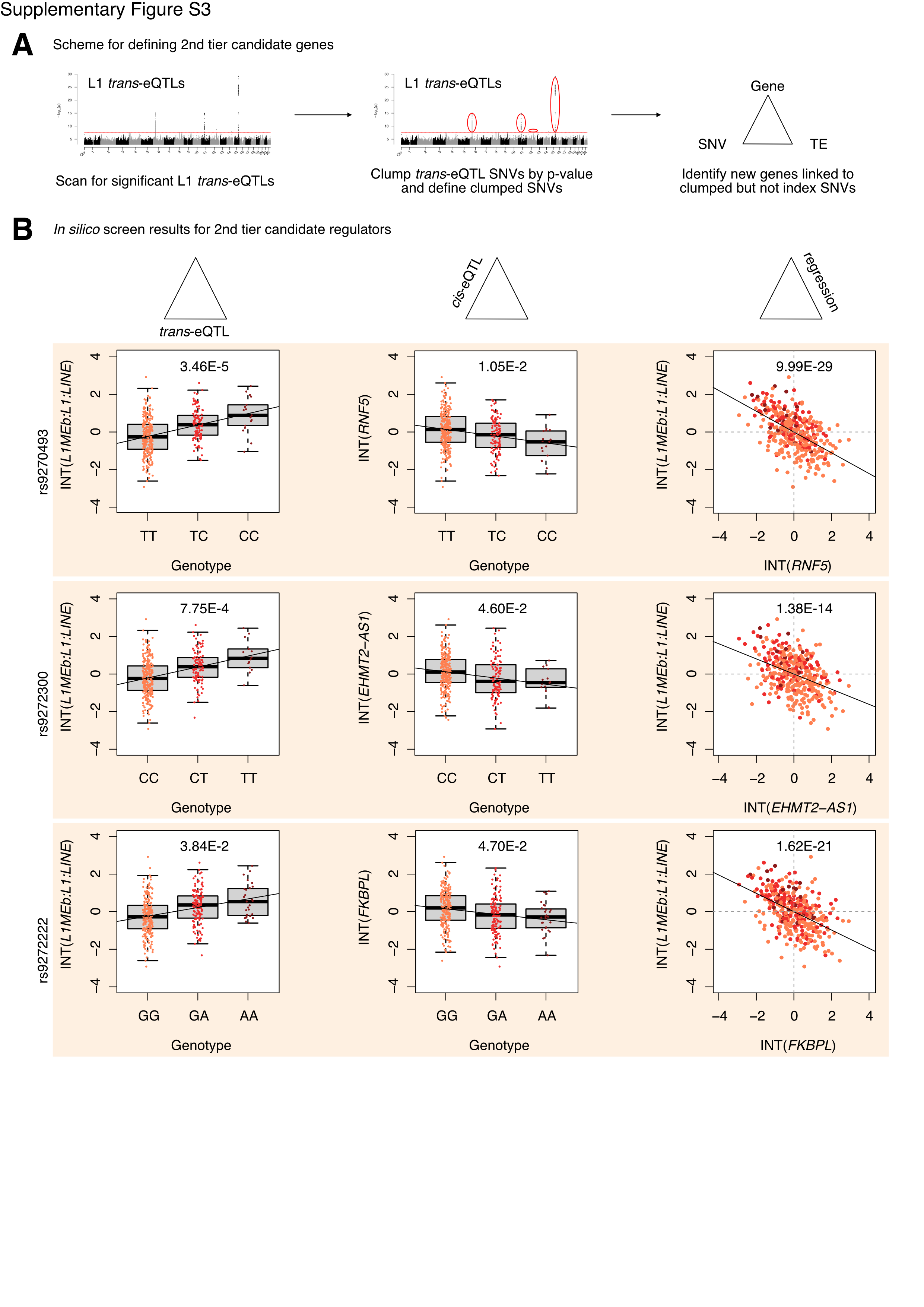

**Supplementary Figure S3. Identification of 2^nd^ tier candidate L1 RNA level regulators in the European cohort.**

**(A)** A schematic for how 2^nd^ tier candidate genes were defined. In short, these were genes in trios with clumped SNVs but not index SNVs at the top of each peak. **(B)** The three-part integration results for three genes—*RNF5*, *EHMT2-AS1*, *FKBPL*—that we considered second tier candidates for functional, *in vitro* testing. In the left column are the ­*trans*-eQTLs, in the middle column are the *cis*-eQTLs, and in the right column are the linear regressions for gene expression against L1 subfamily RNA levels. Expression values following an inverse normal transform (INT) are shown. The FDR for each analysis is listed at the top of each plot. FDR: False Discovery Rate.

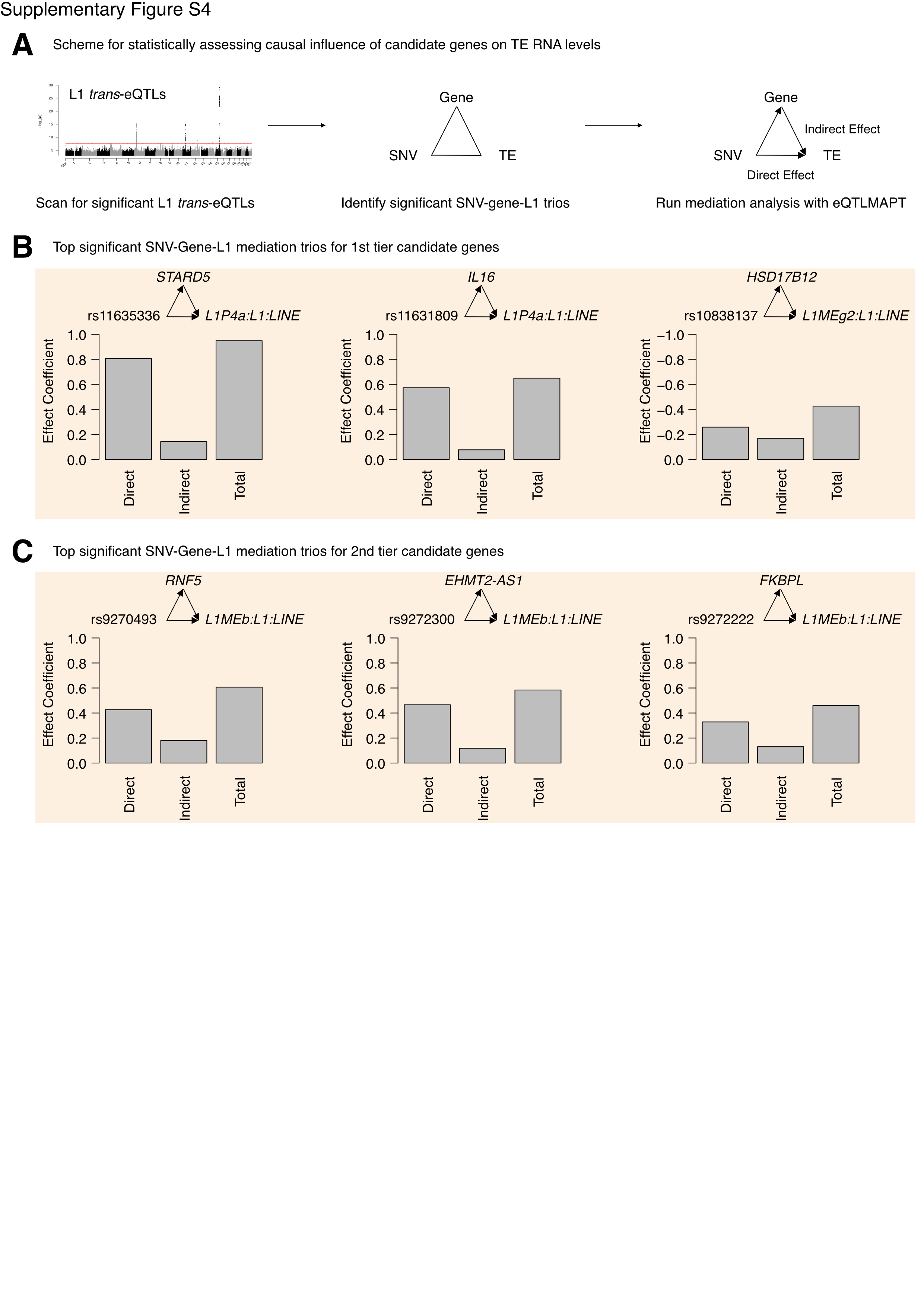

**Supplementary Figure S4. 1^st^ and 2^nd^ tier candidate regulators are partial mediators of SNV effects on L1 RNA levels.**

**(A)** Scheme for the mediation analysis. Mediation analysis tests the mechanistic model where a given gene, *in cis* to a given SNV, partially or fully mediates the effect that SNV has on TE RNA levels *in trans*. The direct, indirect, and total effects for **(B)** 1st tier candidate gene trios and **(C)** 2^nd^ tier candidate gene trios are shown. Mediation was considered significant if the FDR-adjusted empirical p-value, calculated from 30,000 permutations, was < 5%. FDR: False Discovery Rate.

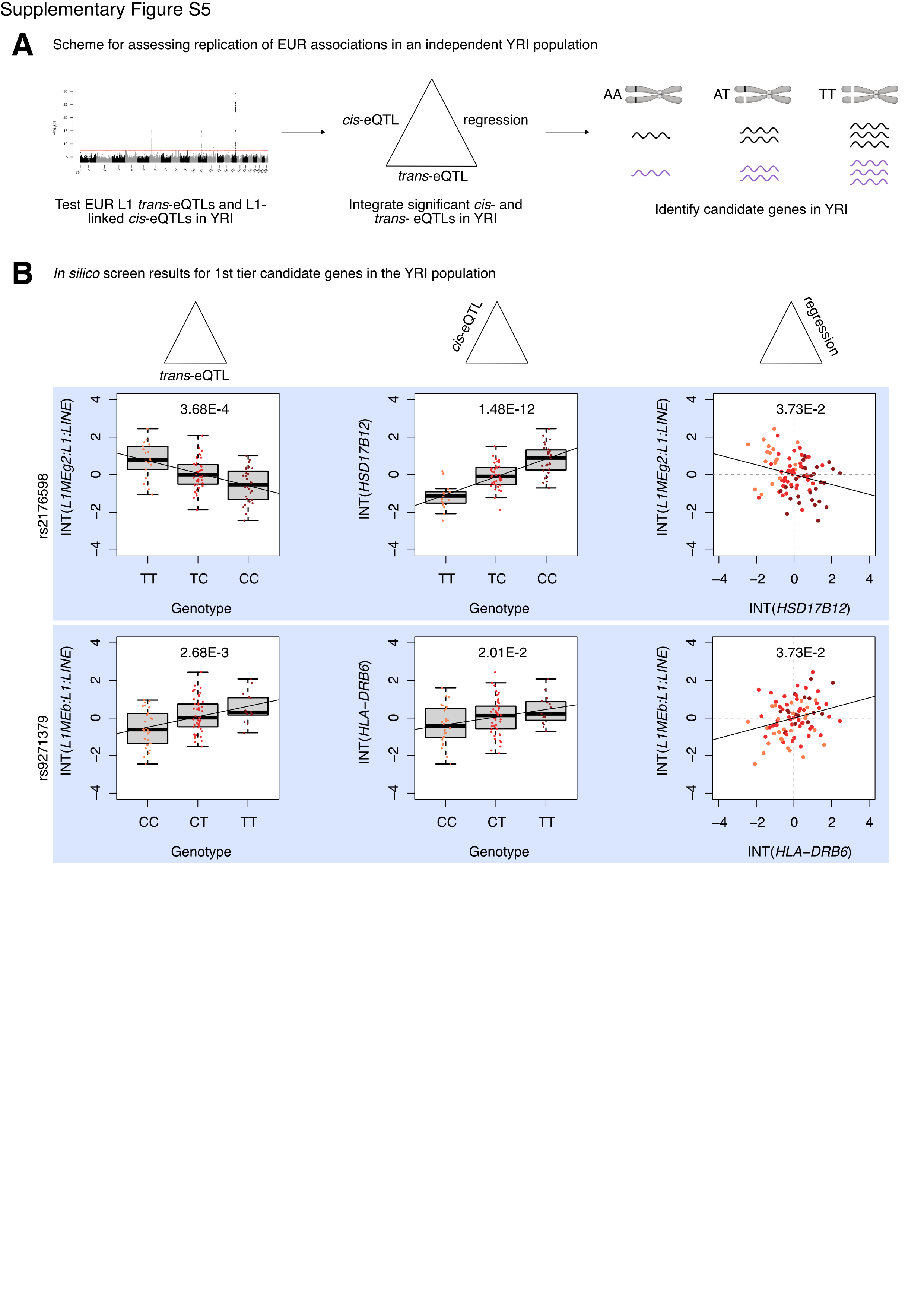

**Supplementary Figure S5**. ***In silico* scanning for candidate L1 RNA level regulators in an independent, African population.**

**(A)** A scheme for the *in silico* analysis carried out with an African cohort, which was similar to the analysis with the European cohort. Since the sample size was smaller than the European cohort, a targeted eQTL approach was undertaken, where we 1) only checked for replication of significant L1 *trans*-eQTLs and 2) only tested significant *trans*-eQTL SNVs for *cis*-association with genes that were L1-linked in the European cohort analysis. **(B)** The three-part integration results for two 1^st^ tier candidate regulators in the African cohort— *HSD17B12* and *HLA-DRB6*. In the left column are the ­*trans*-eQTLs, in the middle column are the *cis*-eQTLs, and in the right column are the linear regressions for gene expression against L1 subfamily expression. Expression values following an inverse normal transform (INT) are shown. The FDR for each analysis is listed at the top of each plot. FDR: False Discovery Rate. Some panels were created with BioRender.com.

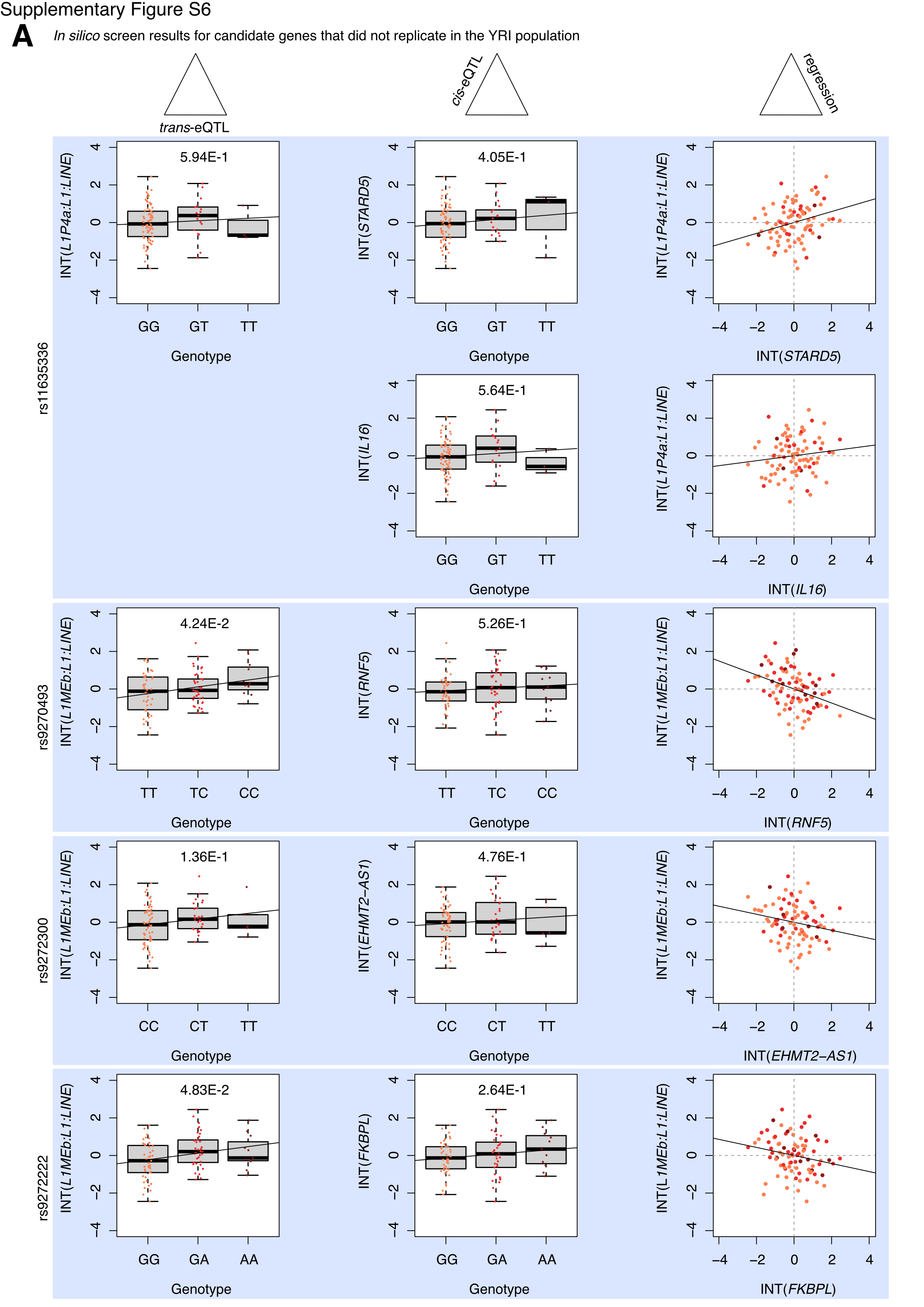

**Supplementary Figure S6. Some European-derived candidate TE-regulator genes do not replicate in the African cohort.**

Many candidate genes from the *in silico* screen in the European cohort did not replicate in the African cohort, likely due to the much smaller sample size and relative rarity of homozygotes carrying 2 alternate alleles. **(A)** The three-part integration results for 1^st^ and 2^nd^ tier candidate regulators identified in the European cohort analysis and tested in the African cohort. In the left column are the ­*trans*-eQTLs, in the middle column are the *cis*-eQTLs, and in the right column are the linear regressions for gene expression against L1 subfamily expression. Expression values following an inverse normal transform (INT) are shown. The FDR is listed for each integration step except for the linear regressions since trios were filtered out before the regression step. FDR: False Discovery Rate.

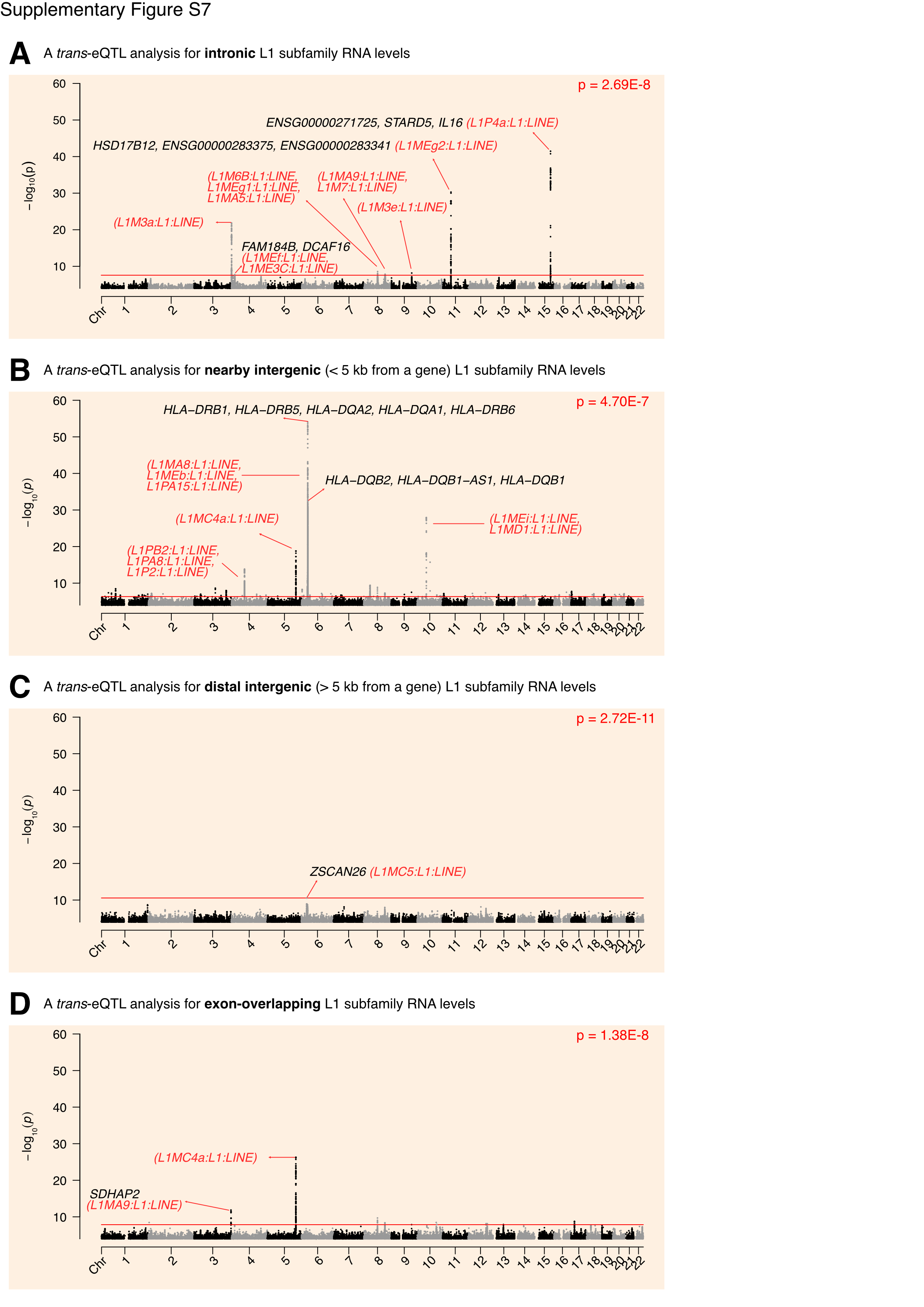

**Supplementary Figure S7. Scanning for candidate regulators of intronic, intergenic, or exonic L1 RNA levels.**

Transposon locus-specific quantifications were obtained using the TElocal package, and these were stratified by genomic region (i.e. intronic, nearby intergenic for loci < 5 kb from a gene, distal intergenic for loci > 5 kb from a gene, and exon-overlapping). Counts were then aggregated at the subfamily level, and the L1 eQTL scan was re-run using each of the four L1 expression profiles using the European cohort. The Manhattan plots for the **(A)** intronic L1 subfamily, **(B)** nearby intergenic L1 subfamily, **(C)** distal intergenic L1 subfamily, and **(D)** exon-overlapping L1 subfamily *trans*-eQTL analyses. For readability, only a subset of associated genes and L1s are highlighted in each plot. For regions with *trans*-eQTLs for multiple L1 subfamilies, we used an un-pointed line to depict the association between the listed L1 subfamilies and at least one SNV in that region. The solid red line in each plot corresponds to a Benjamini-Hochberg FDR < 0.05. FDR: False Discovery Rate.

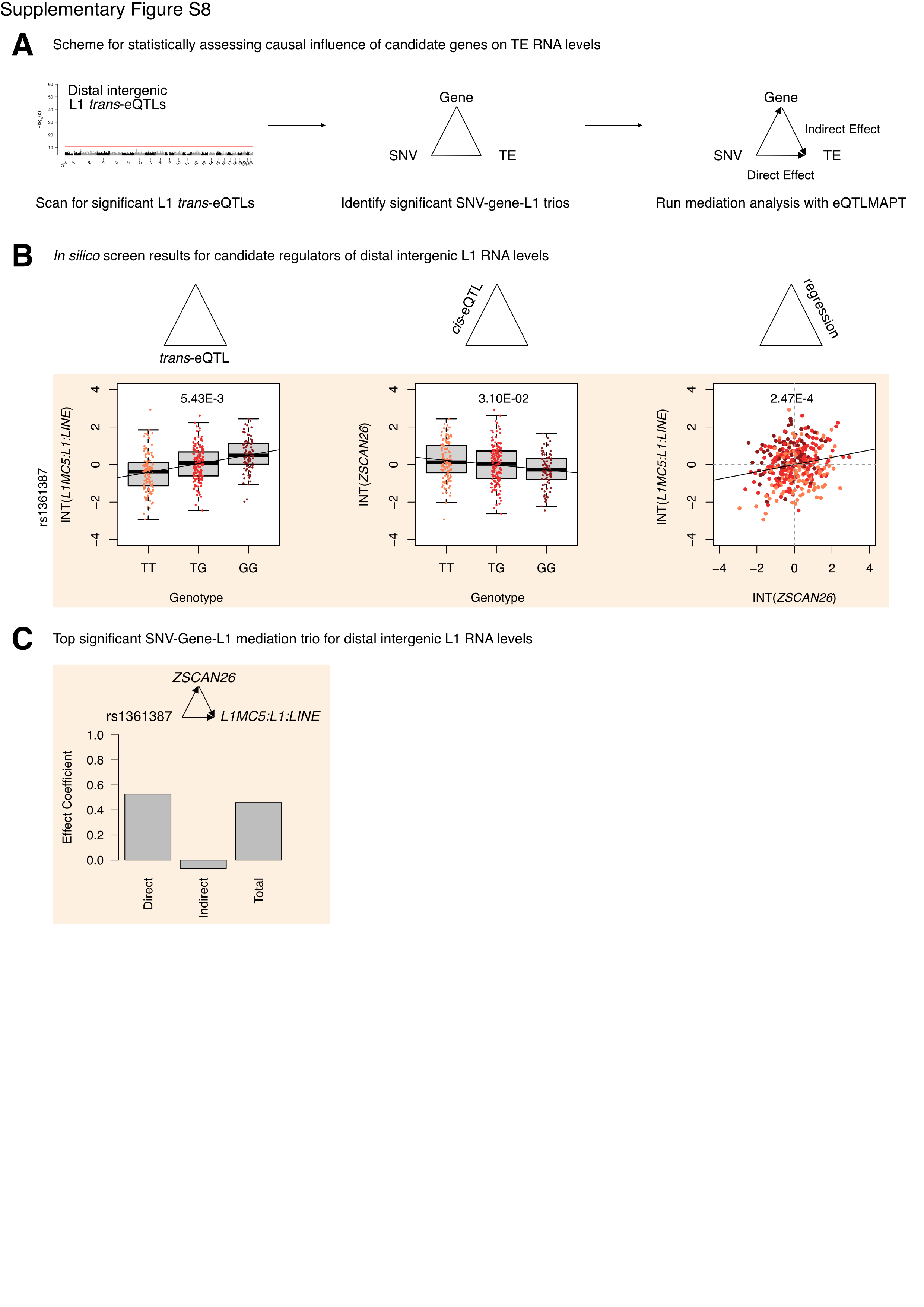

**Supplementary Figure S8. Identifying mediators of SNV effects on distal intergenic L1 RNA levels.**

**(A)** Scheme for the mediation analysis. Mediation analysis tests the mechanistic model where a given gene, *in cis* to a given SNV, partially or fully mediates the effect that SNV has on TE expression *in trans*. **(B)** The three-part integration results for one protein-coding gene—*ZSCAN26*—that we considered a candidate regulator of distal intergenic L1 RNa levels. In the left column are the ­*trans*-eQTLs, in the middle column are the *cis*-eQTLs, and in the right column are the linear regressions for gene expression against L1 subfamily RNA levels. Expression values following an inverse normal transform (INT) are shown. The FDR for each analysis is listed at the top of each plot. **(C)** The *ZSCAN26* SNV-gene-TE mediation trio results. Mediation was considered significant if the FDR-adjusted empirical p-value, calculated from 30,000 permutations, was < 5%. FDR: False Discovery Rate.

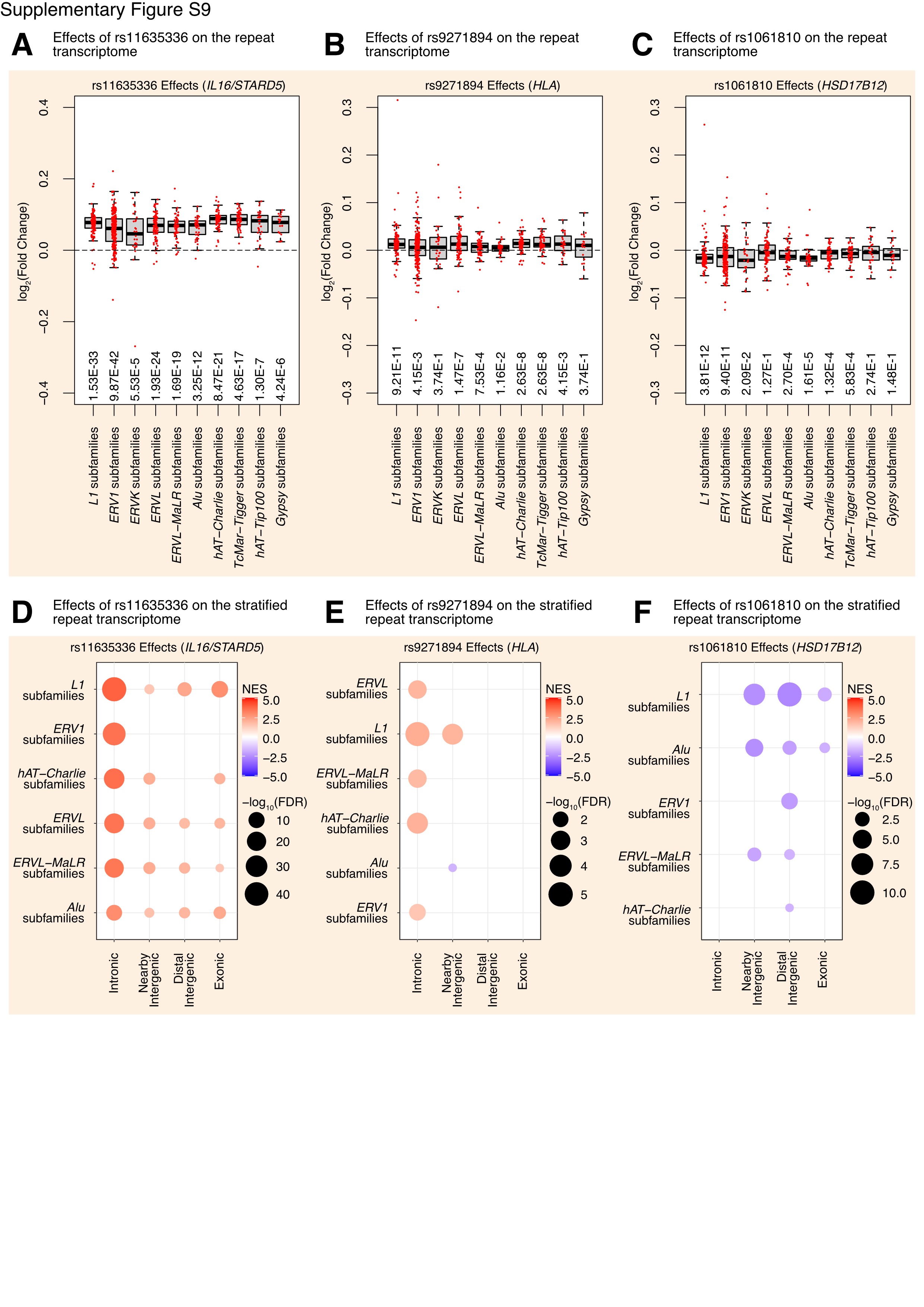

**Supplementary Figure S9.** **Total L1 *trans*-eQTLs alter the expression of TEs in distinct genomic regions.**

Box and whisker plots for the log_2_ fold changes of TE subfamilies (red dots), grouped by TE family, across genotypes for **(A)** rs11635336 (*IL16/STARD5*), **(B)** rs9271894 (*HLA*), and **(C)** rs1061810 (*HSD17B12*). A one-sample Wilcoxon test was run to determine whether changes were significantly different from 0. The FDR values from this test are listed at the bottom. GSEA analysis for top, differentially regulated TE family gene sets in different genomic regions (intronic, intergenic, exon-overlapping) across genotypes. The results for **(D)** rs11635336 (*IL16/STARD5*), **(E)** rs9271894 (*HLA*), and **(F)** rs1061810 (*HSD17B12*) are shown. In each bubble plot, the size of the dot represents the -log_10_(FDR) and the color reflects the normalized enrichment score. FDR: False Discovery Rate.

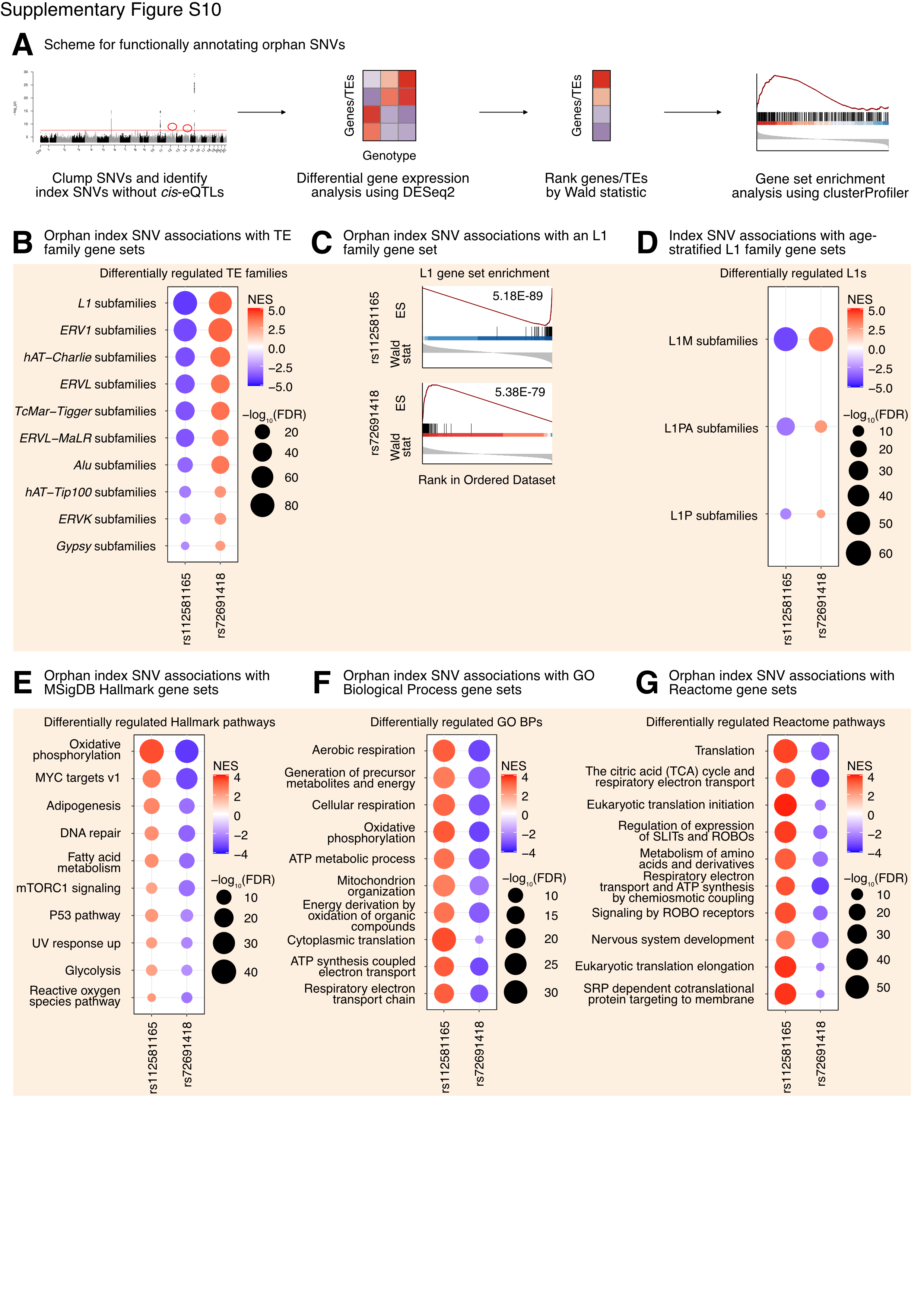

**Supplementary Figure S10. L1 *trans*-eQTL orphan SNVs are associated with differences in TE families and TE-associated pathways.**

**(A)** Scheme for functionally annotating orphan index SNVs by GSEA. **(B)** GSEA analysis for shared, significantly regulated TE family gene sets across genotypes for rs112581165 and rs72691418. **(C)** GSEA plots for the L1 family gene set results summarized in **(B)**. For these plots, the FDR value is listed. **(D)** GSEA analysis for shared, significantly regulated, evolutionary-age-stratified L1 gene sets across genotypes for rs112581165 and rs72691418. L1M subfamilies are the oldest, L1P subfamilies are intermediate, and L1PA subfamilies are the youngest. GSEA analysis for top, shared, concomitantly regulated **(E)** MSigDB Hallmark pathway, **(F)** GO Biological Process, and **(G)** Reactome pathway gene sets across genotypes for rs112581165 and rs72691418. Shared gene sets were ranked by combining p-values from each individual SNV analysis using Fisher’s method. In each bubble plot, the size of the dot represents the -log_10_(FDR) and the color reflects the normalized enrichment score. FDR: False Discovery Rate.

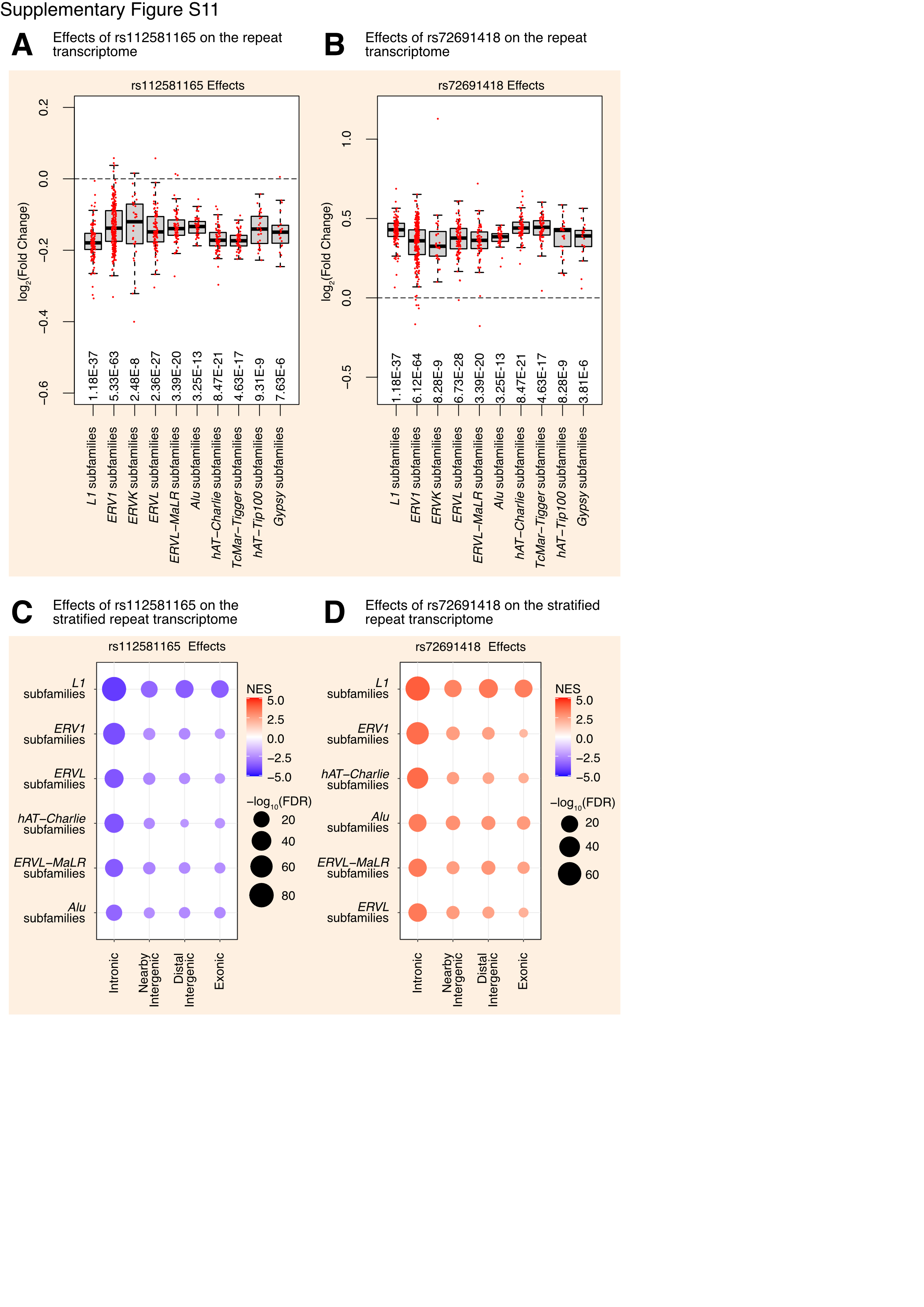

**Supplementary Figure S11.** **L1 *trans*-eQTLs, in the absence of a known mediator, alter the levels of TEs in distinct genomic regions.**

Box and whisker plots for the log_2_ fold changes of TE subfamilies (red dots), grouped by TE family, across genotypes for **(A)** rs112581165 and **(B)** rs72691418. A one-sample Wilcoxon test was run to determine whether changes were significantly different from 0. The FDR values from this test are listed at the bottom. GSEA analysis for top, differentially regulated TE family gene sets in different genomic regions (intronic, intergenic, exon-overlapping) across genotypes. The results for **(C)** rs112581165 and **(D)** rs72691418 are shown. In each bubble plot, the size of the dot represents the -log_10_(FDR) and the color reflects the normalized enrichment score. FDR: False Discovery Rate.

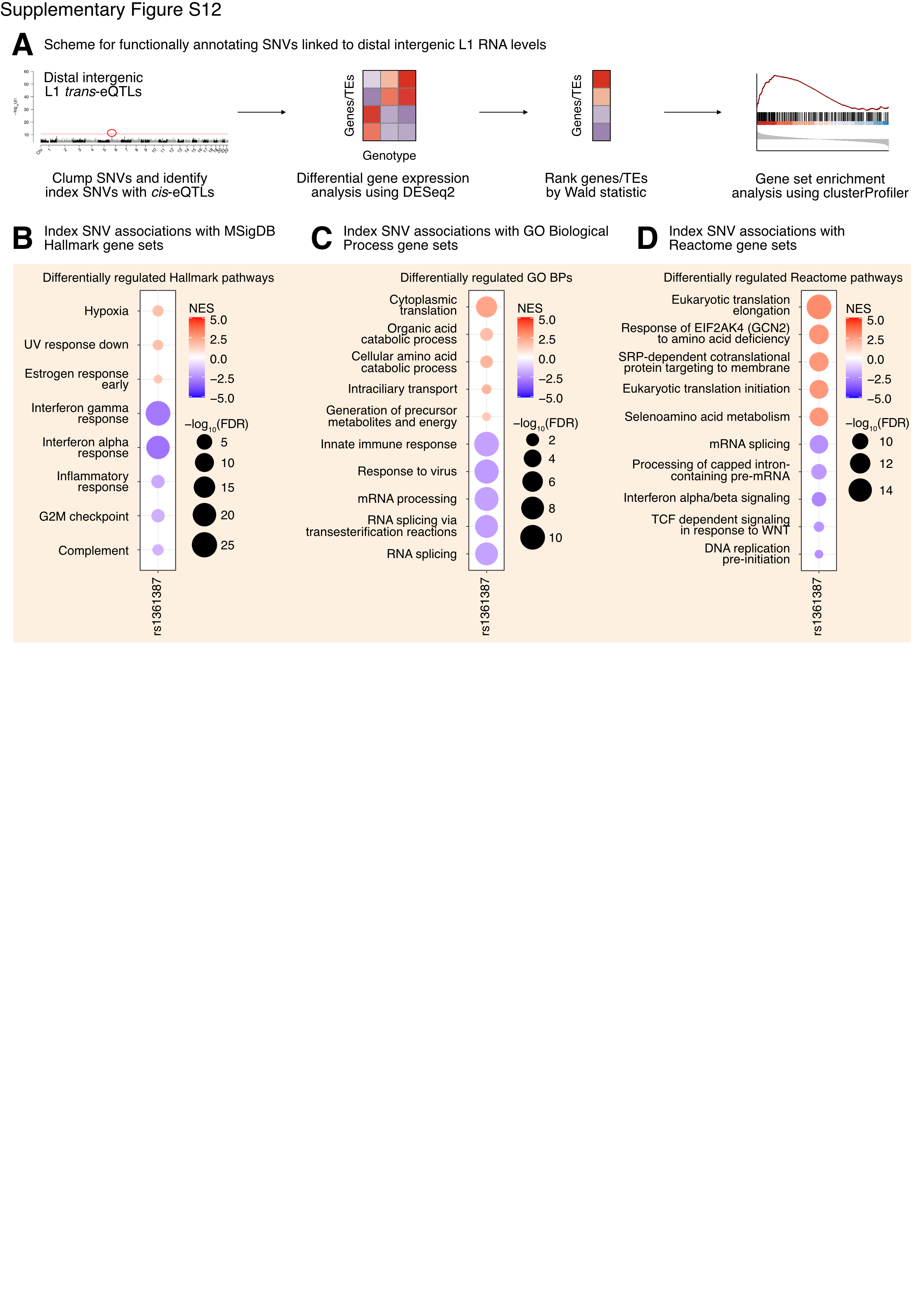

**Figure S12. Distal intergenic L1 *trans*-eQTLs are associated with alterations in inflammatory pathways.**

**(A)** Scheme for functionally annotating gene-linked index SNVs for distal intergenic L1 expression by GSEA. GSEA analysis for top regulated **(B)** MSigDB Hallmark pathway, **(C)** GO Biological Process, and **(D)** Reactome pathway gene sets across genotypes for rs1361387 (*ZSCAN26)*. In each bubble plot, the size of the dot represents the -log_10_(FDR) and the color reflects the normalized enrichment score. FDR: False Discovery Rate.

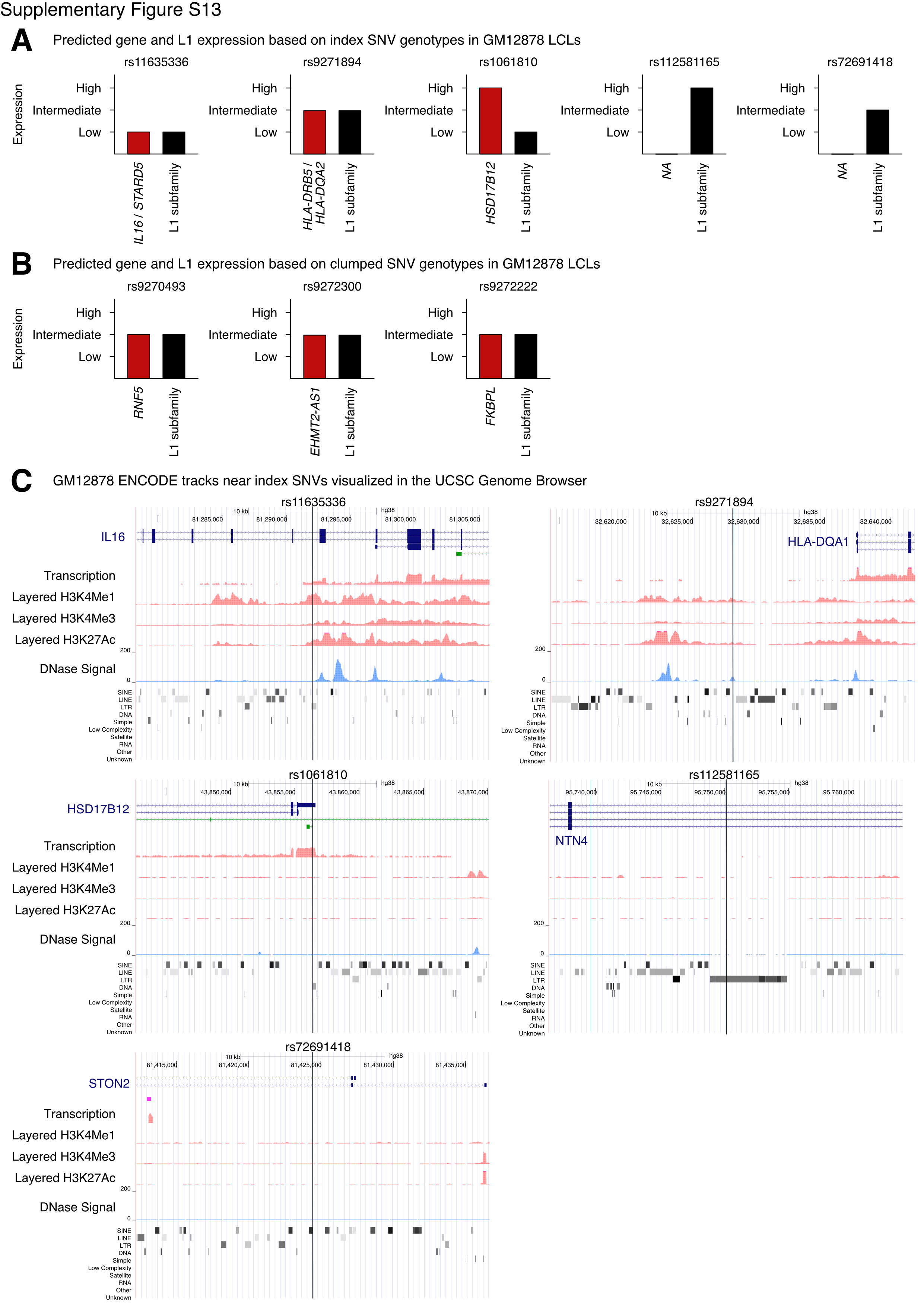

**Supplementary Figure S13. GM12878 LCLs as a model for assessing the roles of candidate genes in L1 regulation.**

Though transcriptomic data is not available for these LCLs, the relative expression of candidate genes and linked L1 subfamilies can be predicted from GM12878 genotypes at either **(A)** index SNVs or **(B)** clumped SNVs. **(C)** ENCODE project epigenetic data available for GM12878 highlights regulatory markers near some *trans*-eQTL index SNVs. Data is visualized on the UCSC Genome Browser.

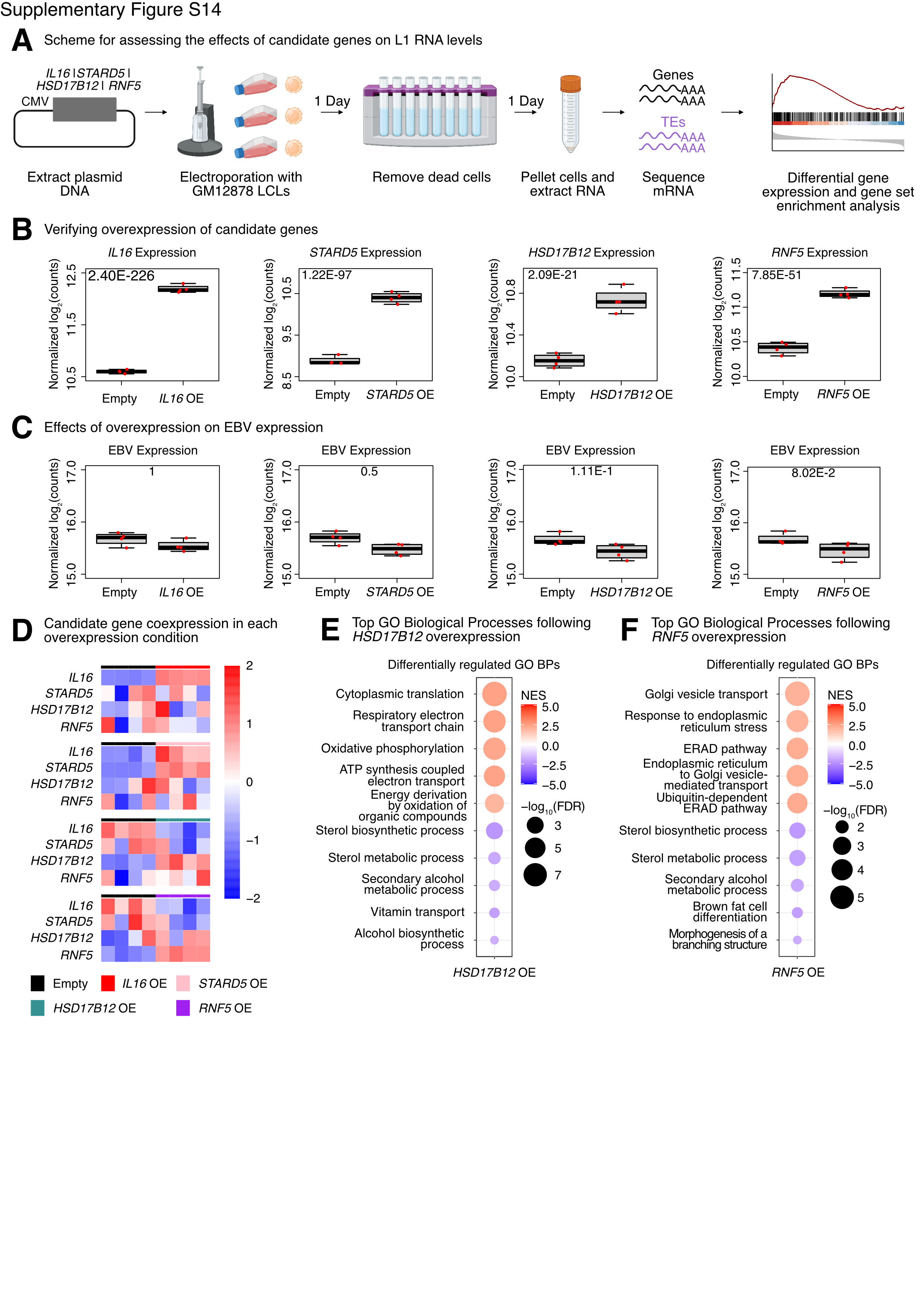

**Supplementary Figure S14.** **Though *HSD17B12* and *RNF5* were overexpressed, no L1 gene set changes were detected.**

**(A)** Scheme for experimentally validating the roles of *IL16,* *STARD5, HSD17B12,* and *RNF5* in L1 regulation. **(B)** VST-normalized log_2_ counts were quantified by DESeq2 for each gene being overexpressed. Each dot represents an independent transfection, with n = 4 per condition. The FDR for each comparison is listed at the top. **(C)** VST-normalized log_2_ counts for EBV were quantified by DESeq2 for each condition. Each dot represents an independent transfection, with n = 4 per condition. The FDR for each comparison is listed at the top. **(D)** Expression heatmaps for the four candidate genes tested, under each overexpression condition. GSEA analysis for top, differentially regulated **(E)** GO Biological Process gene sets following *HSD17B12* overexpression. GSEA analysis for top, differentially regulated **(F)** GO Biological Process gene sets following *RNF5* overexpression. In each bubble plot, the size of the dot represents the -log_10_(FDR) and the color reflects the normalized enrichment score. FDR: False Discovery Rate. Some panels were created with BioRender.com.

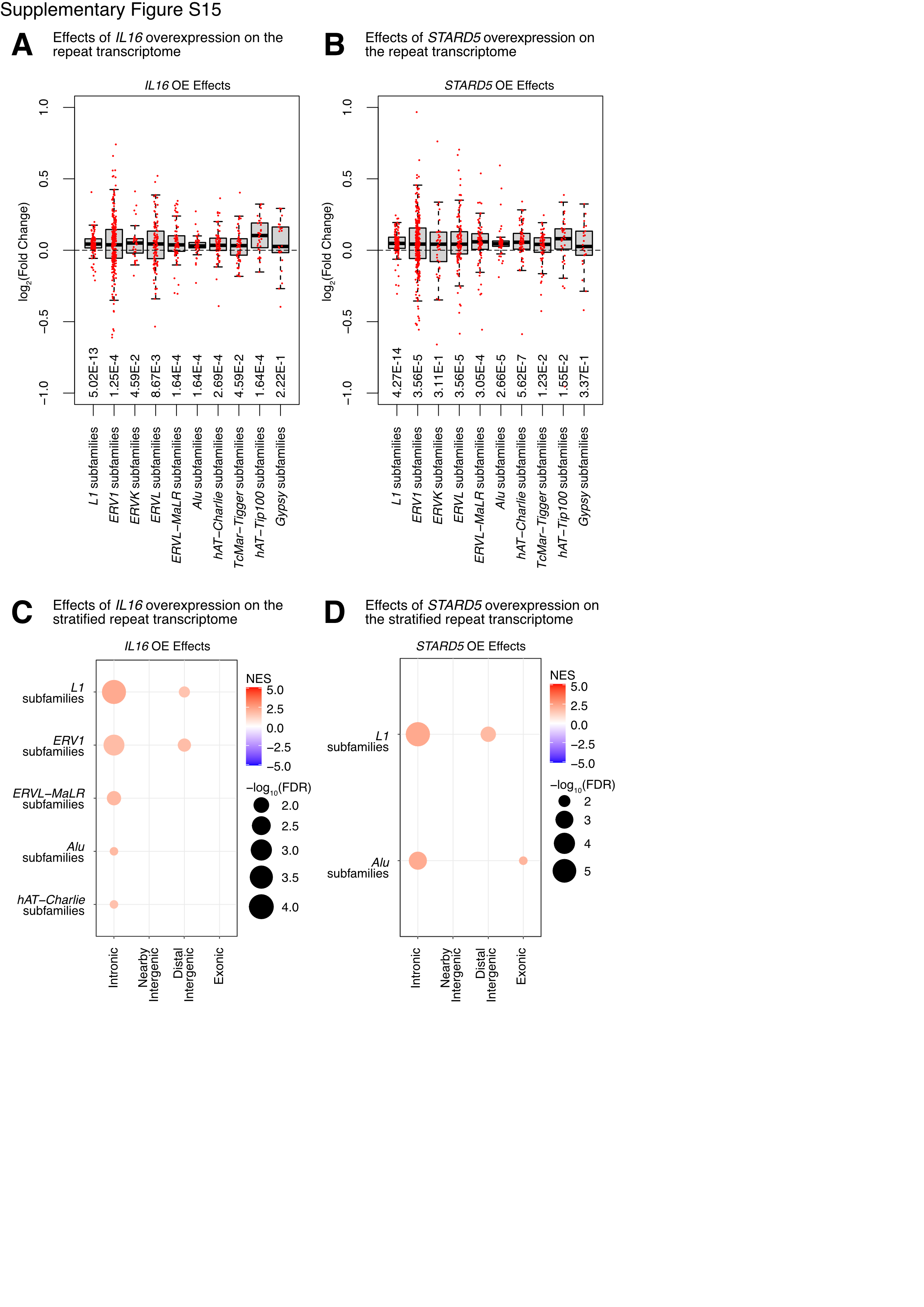

**Supplementary Figure S15. *IL16* and *STARD5* overexpression alters the levels of TEs in distinct genomic regions.**

Box and whisker plots for the log_2_ fold changes of TE subfamilies (red dots), grouped by TE family, following **(A)** *IL16* overexpression and **(B)** *STARD5* overexpression. A one-sample Wilcoxon test was run to determine whether changes were significantly different from 0. The FDR values from this test are listed at the bottom. GSEA analysis for top, differentially regulated TE family gene sets in different genomic regions (intronic, intergenic, exon-overlapping) across overexpression condition. The results for **(C)** *IL16* overexpression and **(D)** *STARD5* overexpression are shown. In each bubble plot, the size of the dot represents the -log_10_(FDR) and the color reflects the normalized enrichment score. FDR: False Discovery Rate.

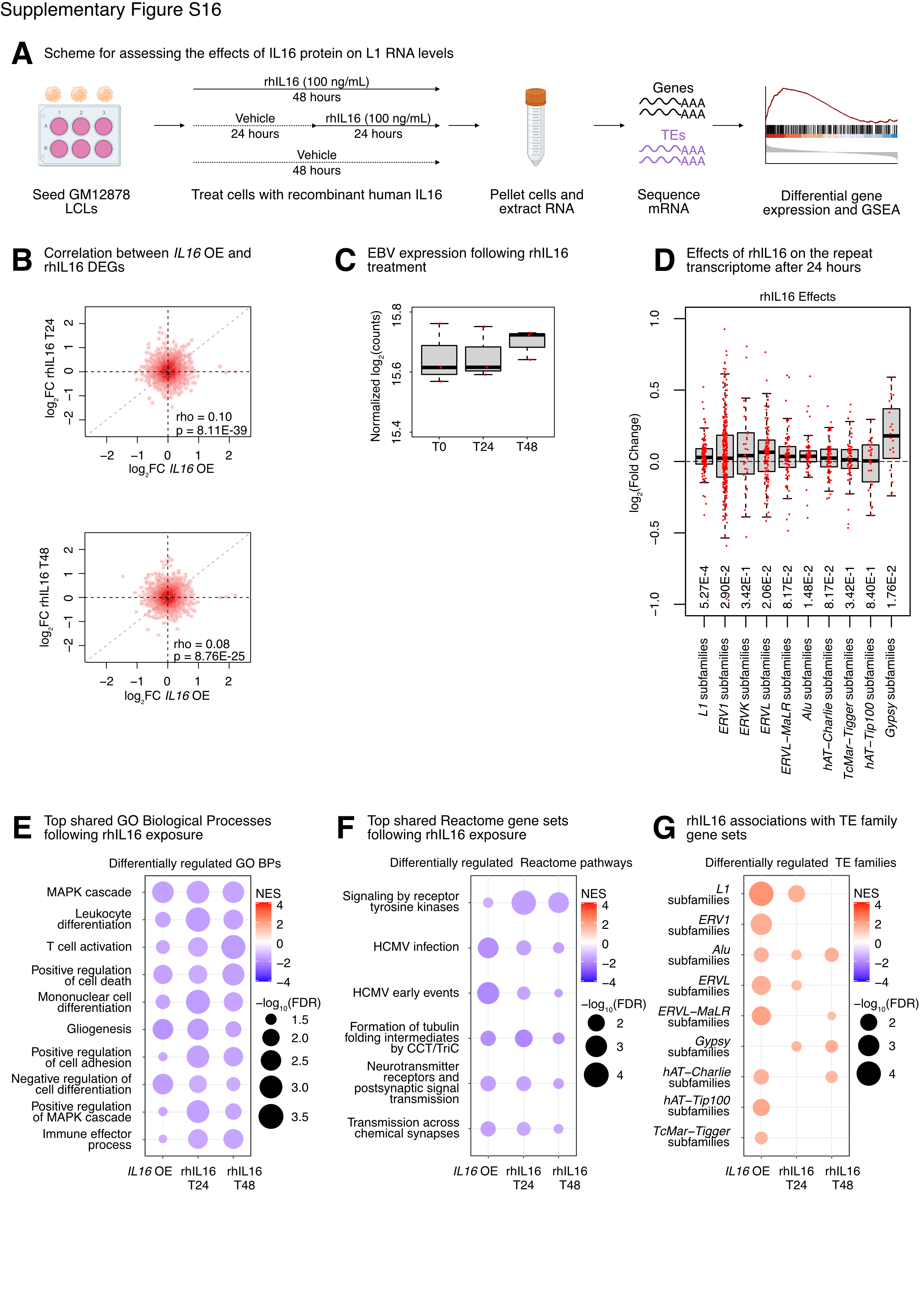

**Supplementary Figure S16. The effects of rhIL16 on an L1 family gene set are diminished after 48 hours.**

**(A)** Scheme for experimentally validating the role of rhIL16 in L1 regulation. **(B)** Spearman rank correlation between (*top)* 24 hour rhIL16 treatment differential expression or (*bottom)* 48 hour rhIL16 treatment differential expression and *IL16* overexpression differential expression. **(C)** VST-normalized log_2_ counts for EBV were quantified by DESeq2 for each peptide treatment condition. Each dot represents an independent exposure, with n = 3 per condition. FDR > 0.05 at both the 24 and 48 hour treatment time points. **(D)** Box and whisker plots for the log_2_ fold changes of TE subfamilies (red dots) grouped by TE family after treatment with rhIL16 for 24 hours. A one-sample Wilcoxon test was run to determine whether changes were significantly different from 0. The FDR values from this test are listed at the bottom. GSEA analysis for top, shared, concomitantly regulated **(E)** GO Biological Process, **(F)** Reactome pathway, and **(G)** TE family gene sets following *IL16* overexpression, rhIL16 exposure for 24 hours, and rhIL16 exposure for 48 hours. Shared gene sets were ranked by combining p-values from each individual treatment analysis using Fisher’s method. In each bubble plot, the size of the dot represents the -log_10_(FDR) and the color reflects the normalized enrichment score. FDR: False Discovery Rate. Some panels were created with BioRender.com.

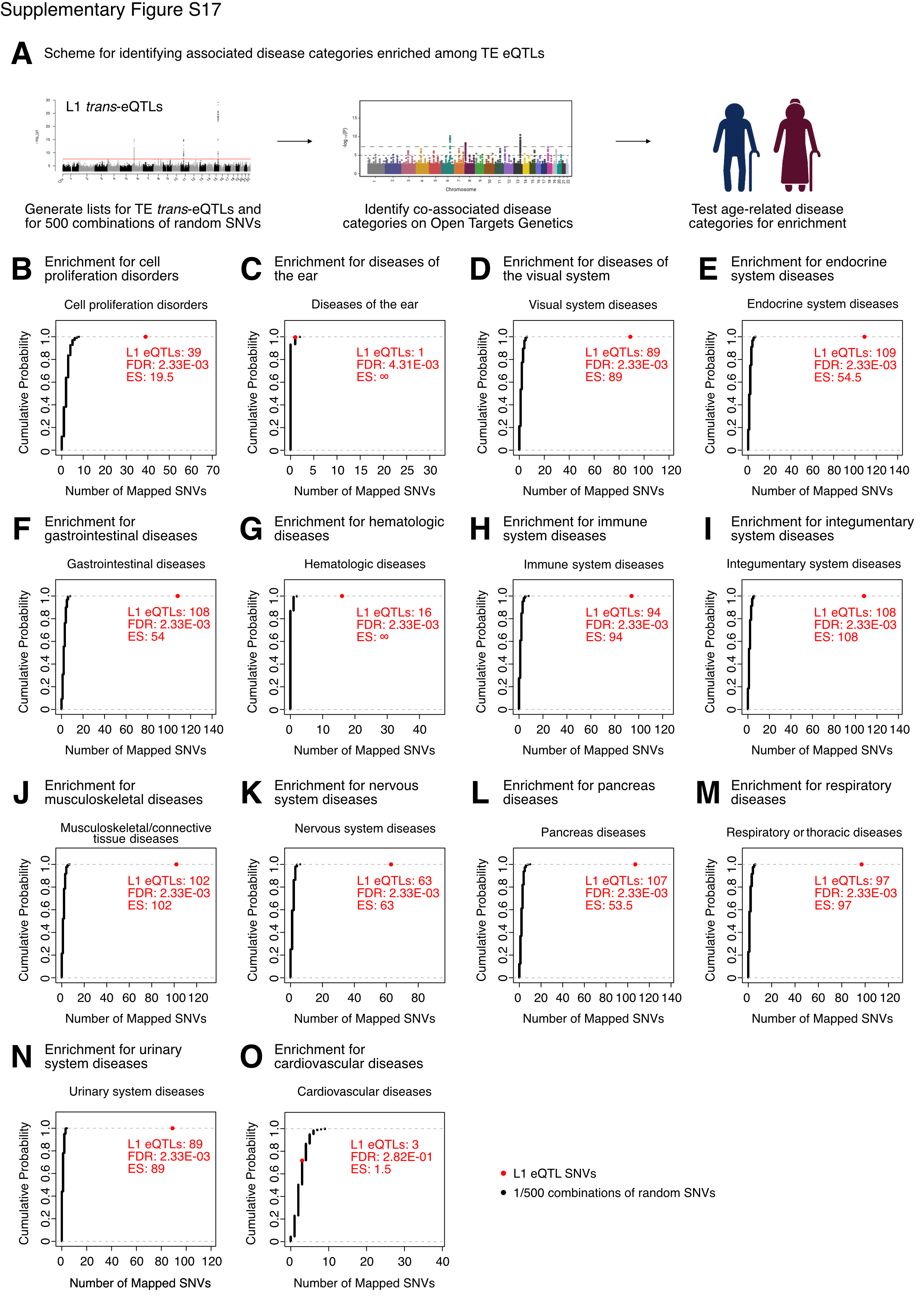

**Supplementary Figure S17. Aging-related disease categories are enriched among L1 *trans*-eQTL PheWAS associations.**

**(A)** Scheme for assessing enrichment of disease category associations among L1 *trans*-eQTLs. L1 eQTLs or combinations of random SNVs were queried on the Open Targets Genetics platform, and the number of SNVs mapping to 14 disease categories annotated by the platform were calculated. An empirical cumulative distribution function (ecdf) was defined for each disease category using the associations for the random SNVs, and this function was used to calculate an enrichment p-value, defined as p = 1 – ecdf(mapped eQTLs). Afterwards, p-values were FDR-corrected. An enrichment score (ES) was calculated by taking the number of mapped L1 eQTLs for a category and dividing by the median number of mapped SNVs among the random SNV combinations. Categories with an ES > 1 and FDR < 0.05 were considered significantly enriched. The number of mapped SNVs relative to the cumulative probabilities from the simulations are shown for **(B)** cell proliferation disorders, **(C)** diseases of the ear, **(D)** diseases of the visual system, **(E)** endocrine system diseases, **(F)** gastrointestinal diseases, **(G)** hematologic diseases, **(H)** immune system diseases, **(I)** integumentary system diseases, **(J)** musculoskeletal diseases, **(K)** nervous system diseases, **(L)** pancreas diseases, **(M)** respiratory diseases, **(N)** urinary system diseases, and **(O)** cardiovascular diseases. FDR: False Discovery Rate. Some panels were created with BioRender.com.

**Inventory of Supplementary Tables**

**Supplementary table S1. Results for eQTL scans and mediation analysis in the European cohort.**

**(A)** L1 subfamily *trans*-eQTLs passing FDR < 0.05 in the European populations. **(B)** Gene *cis*-eQTLs passing FDR < 0.05 in the European populations. **(C)** SNV-Gene-L1 trios passing FDR < 0.05 in the *cis*-eQTL, *trans*-eQTL, and linear regression analyses in the European cohort. **(D)** European L1 *trans*-eQTLs clumped by p-value. **(E)** Significant SNV-Gene-L1 trios that survived clumping in the European populations. **(F)** Mediation analysis results for all SNV-Gene-L1 trios in the European cohort. FDR: False Discovery Rate.

**Supplementary table S2. Results for eQTL scans and mediation analysis in the African cohort.**

**(A)** L1 subfamily *trans*-eQTLs passing FDR < 0.05 in the Yoruban population. **(B)** Gene *cis*-eQTLs passing FDR < 0.05 in the Yoruban population. **(C)** SNV-Gene-L1 trios passing FDR < 0.05 in the *cis*-eQTL, *trans*-eQTL, and linear regression analyses in the Yoruban population. **(D)** Yoruban L1 *trans*-eQTLs clumped by p-value. **(E)** Significant SNV-Gene-L1 trios that survived clumping in the Yoruban population. **(F)** Mediation analysis results for all SNV-Gene-L1 trios in the Yoruban cohort. FDR: False Discovery Rate.

**Supplementary table S3. Results for eQTL scans using intronic, intergenic, or exon-overlapping L1 subfamily RNA levels.**

**(A)** Intronic L1 subfamily trans-eQTLs passing FDR < 0.05 in the EUR population. **(B)** Nearby Intergenic L1 subfamily trans-eQTLs passing FDR < 0.05 in the EUR population. **(C)** Distal Intergenic L1 subfamily trans-eQTLs passing FDR < 0.05 in the EUR population. **(D)** Exon overlapping L1 subfamily trans-eQTLs passing FDR < 0.05 in the EUR population. **(E)** Significant SNV-Gene-L1 trios for intronic L1 subfamilies. **(F)** Significant SNV-Gene-L1 trios for nearby intergenic L1 subfamilies. **(G)** Significant SNV-Gene-L1 trios for distal intergenic L1 subfamilies. **(H)** Significant SNV-Gene-L1 trios for exon-overlapping L1 subfamilies. **(I)** Mediation analysis results for distal intergenic L1-SNV-gene trios.

**Supplementary table S4. Results for the differential gene expression analysis and GSEA comparing genotypes in SNVs with an attributed *cis*-mediator.**

**(A)** All DESeq2 results for alternating the allele of rs11635336. **(B)** All DESeq2 results for alternating the allele of rs9271894. **(C)** All DESeq2 results for alternating the allele of rs1061810. **(D)** All GSEA results for rs11635336 using TE family gene sets. **(E)** All GSEA results for rs9271894 using TE family gene sets. **(F)** All GSEA results for rs1061810 using TE family gene sets. **(G)** All GSEA results for rs11635336 using MSigDB Hallmark gene sets. **(H)** All GSEA results for rs9271894 using MSigDB Hallmark gene sets. **(I)** All GSEA results for rs1061810 using MSigDB Hallmark gene sets. **(J)** All GSEA results for rs11635336 using GO Biological Process gene sets. **(K)** All GSEA results for rs9271894 using GO Biological Process gene sets. **(L)** All GSEA results for rs1061810 using GO Biological Process gene sets. **(M)** All GSEA results for rs11635336 using Reactome gene sets. **(N)** All GSEA results for rs9271894 using Reactome gene sets. **(O)** All GSEA results for rs1061810 using Reactome gene sets. **(P)** Overlapping TE family gene sets for rs11635336, rs9271894, rs1061810. **(Q)** Overlapping age-stratified TE family gene sets for rs11635336, rs9271894, rs1061810. **(R)** All DESeq2 results for alternating the allele of rs9270493. **(S)** All GSEA results for rs9270493 using TE family gene sets. **(T)** GSEA results for genomic region-stratified TE family gene sets for rs11635336. **(U)** GSEA results for genomic region-stratified TE family gene sets for rs9271894. **(V)** GSEA results for genomic region-stratified TE family gene sets for rs1061810. **(W)** Overlapping MSigDB Hallmark gene sets for rs11635336, rs9271894, rs1061810. **(X)** Overlapping GO Biological Process gene sets for rs11635336, rs9271894, rs1061810. **(Y)** Overlapping Reactome gene sets for rs11635336, rs9271894, rs1061810.

**Supplementary table S5. Results for the differential gene expression analysis and GSEA comparing genotypes in orphan index SNVs and distal intergenic-associated SNVs.**

**(A)** All DESeq2 results for alternating the allele of rs112581165. **(B)** All DESeq2 results for alternating the allele of rs72691418. **(C)** All GSEA results for rs112581165 using TE family gene sets. **(D)** All GSEA results for rs72691418 using TE family gene sets. **(E)** All GSEA results for rs112581165 using MSigDB Hallmark gene sets. **(F)** All GSEA results for rs72691418 using MSigDB Hallmark gene sets. **(G)** All GSEA results for rs112581165 using GO Biological Process gene sets. **(H)** All GSEA results for rs72691418 using GO Biological Process gene sets. **(I)** All GSEA results for rs112581165 using Reactome gene sets. **(J)** All GSEA results for rs72691418 using Reactome gene sets. **(K)** Overlapping TE family gene sets for rs112581165 and rs72691418. **(L)** Overlapping age-stratified TE family gene sets for rs112581165 and rs72691418. **(M)** GSEA results for genomic region-stratified TE family gene sets for rs112581165. **(N)** GSEA results for genomic region-stratified TE family gene sets for rs72691418. **(O)** Overlapping MSigDB Hallmark gene sets for rs112581165 and rs72691418. **(P)** Overlapping GO Biological Process gene sets for rs112581165 and rs72691418. **(Q)** Overlapping Reactome gene sets for rs112581165 and rs72691418. **(R)** All DESeq2 results for alternating the allele of rs1361387. **(S)** All GSEA results for rs1361387 using MSigDB Hallmark gene sets. **(T)** All GSEA results for rs1361387 using GO Biological Process gene sets. **(U)** All GSEA results for rs1361387 using Reactome gene sets.

**Supplementary table S6. Results for the differential gene expression analysis and GSEA assessing the effects of overexpressing *IL16, STARD5, HSD17B12, or RNF5*.**

**(A)** All DESeq2 results for *IL16* overexpression. **(B)** All DESeq2 results for *STARD5* overexpression. **(C)** All DESeq2 results for *HSD17B12* overexpression. **(D)** All DESeq2 results for *RNF5* overexpression. **(E)** All GSEA results for *IL16* overexpression using MSigDB Hallmark gene sets. **(F)** All GSEA results for *STARD5* overexpression using MSigDB Hallmark gene sets. **(G)** All GSEA results for *HSD17B12* overexpression using MSigDB Hallmark gene sets. **(H)** All GSEA results for *RNF5* overexpression using MSigDB Hallmark gene sets. **(I)** All GSEA results for *IL16* overexpression using GO Biological Process gene sets. **(J)** All GSEA results for *STARD5* overexpression using GO Biological Process gene sets. **(K)** All GSEA results for *HSD17B12* overexpression using GO Biological Process gene sets. **(L)** All GSEA results for *RNF5* overexpression using GO Biological Process gene sets. **(M)** All GSEA results for *IL16* overexpression using Reactome gene sets. **(N)** All GSEA results for *STARD5* overexpression using Reactome gene sets. **(O)** All GSEA results for *HSD17B12* overexpression using Reactome gene sets. **(P)** All GSEA results for *RNF5* overexpression using Reactome gene sets. **(Q)** All GSEA results for *IL16* overexpression using TE family gene sets. **(R)** All GSEA results for *STARD5* overexpression using TE family gene sets. **(S)** All GSEA results for *HSD17B12* overexpression using TE family gene sets. **(T)** All GSEA results for *RNF5* overexpression using TE family gene sets. **(U)** GSEA with age-stratified L1 family gene sets following *IL16* and *STARD5* OE. **(V)** GSEA results for genomic region-stratified TE family gene sets for *IL16* OE. **(W)** GSEA results for genomic region-stratified TE family gene sets for *STARD5* OE.

**Supplementary table S7. Results for the differential gene expression analysis and GSEA assessing the effects of rhIL16 exposure.**

**(A)** All DESeq2 results for rhIL16 exposure for 24 hours. **(B)** All GSEA results for 24 hour rhIL16 exposure using MSigDB Hallmark gene sets. **(C)** All GSEA results for 24 hour rhIL16 exposure using GO Biological Process gene sets. **(D)** All GSEA results for 24 hour rhIL16 exposure using Reactome gene sets. **(E)** All GSEA results for 24 hour rhIL16 exposure using TE family gene sets. **(F)** Overlapping MSigDB Hallmark gene sets for *IL16* overexpression and rhIL16 24 hour exposure. **(G)** Overlapping GO Biological Process gene sets for *IL16* overexpression and rhIL16 24 hour exposure. **(H)** Overlapping Reactome gene sets for *IL16* overexpression and rhIL16 24 hour exposure. **(I)** GSEA with age-stratified L1 family gene sets following rhIL16 T24 exposure. **(J)** GSEA results for genomic region-stratified TE family gene sets for the T24 rhIL16 exposure. **(K)** All DESeq2 results for rhIL16 exposure for 48 hours. **(L)** All GSEA results for 48 hour rhIL16 exposure using MSigDB Hallmark gene sets. **(M)** All GSEA results for 48 hour rhIL16 exposure using GO Biological Process gene sets. **(N)** All GSEA results for 48 hour rhIL16 exposure using Reactome gene sets. **(O)** All GSEA results for 48 hour rhIL16 exposure using TE family gene sets. **(P)** Overlapping MSigDB Hallmark gene sets for *IL16* overexpression, rhIL16 24 hour exposure, and rhIL16 48 hour exposure. **(Q)** Overlapping GO Biological Process gene sets for *IL16* overexpression, rhIL16 24 hour exposure, and rhIL16 48 hour exposure. **(R)** Overlapping Reactome gene sets for *IL16* overexpression, rhIL16 24 hour exposure, and rhIL16 48 hour exposure. **(S)** Overlapping TE family gene sets for *IL16* overexpression, rhIL16 24 hour exposure, and rhIL16 48 hour exposure.

**Supplementary table S8. Shared gene sets that are concomitantly and significantly regulated across all conditions with upregulation of an L1 gene set.**

**(A)** Overlapping TE family gene sets for *IL16* overexpression, *STARD5* overexpression, and rhIL16 24 hour exposure. **(B)** Overlapping MSigDB Hallmark gene sets for *IL16* overexpression, *STARD5* overexpression, and rhIL16 24 hour exposure. **(C)** Overlapping GO Biological Process gene sets for *IL16* overexpression, *STARD5* overexpression, and rhIL16 24 hour exposure. **(D)** Overlapping Reactome gene sets for *IL16* overexpression, *STARD5* overexpression, and rhIL16 24 hour exposure.

**Supplementary table S9. Age-related associations with L1 *trans*-eQTLs.**

**(A)** Statistics for SNVs mapping to aging MeSH traits through the Open Targets Genetics platform. **(B)** Number of SNVs mapping to unique MeSH ID disease terms. **(C)** Circulating IL16 concentrations in aging mouse serum.
